# Genome-wide association study of multiple yield components in a diversity panel of polyploid sugarcane (*Saccharum* spp.)

**DOI:** 10.1101/387001

**Authors:** Xiping Yang, Ziliang Luo, James Todd, Sushma Sood, Jianping Wang

**Affiliations:** Agronomy Department, University of Florida, Gainesville, FL 32610, USA; Sugarcane Research Unit, USDA-ARS, Houma, LA 70360; Sugarcane Field Station, USDA, ARS, Canal Point, FL 33438; Center for Genomics and Biotechnology, Key Laboratory of Genetics, Breeding and Multiple Utilization of Corps, Ministry of Education; Fujian Provincial Key Laboratory of Haixia Applied Plant Systems Biology, Fujian Agriculture and Forestry University, Fuzhou 350002, Fujian, China

**Keywords:** Sugarcane germplasm, polyploidy, yield components, target enrichment sequencing, GWAS

## Abstract

Sugarcane (*Saccharum* spp.) is an important economic crop, contributes up to 80% of sugar and approximately 60% bio-fuel globally. To meet the increased demand for sugar and bio-fuel supplies, it is critical to breed sugarcane cultivars with robust performance in yield components. Therefore, dissection of causal DNA sequence variants is of great importance by providing genetic resources and fundamental information for crop improvement. In this study, we evaluated and analyzed nine yield components in a sugarcane diversity panel consisting of 308 accessions primarily selected from the “world collection of sugarcane and related grasses”. By genotyping the diversity panel using target enrichment sequencing, we identified a large number of sequence variants. Genome-wide association study between the markers and traits were conducted with dosages and gene actions taken into consideration. In total, 217 non-redundant markers and 225 candidate genes were identified to be significantly associated with the yield components, which can serve as a comprehensive genetic resource database for future gene identification, characterization, and selection for sugarcane improvement. We further investigated runs of homozygosity (ROH) in the sugarcane diversity panel. We characterized 282 ROHs, and found that the occurrence of ROH in the genome were non-random and probably under selection. ROHs were associated with total weight and dry weight, and high ROHs resulted in decrease of the two traits. This study approved that genomic inbreeding has led to negative impacts on sugarcane yield.

## Introduction

As an important cash crop, sugarcane (*Saccharum* spp.) contributes up to 80% of sugar and approximately 60% bio-fuel globally (Dahlquist 2013). To meet the increasing global demand for sugar and bio-fuel, sugarcane harvest area in the world has increased from 22.7 to 26.8 million hectares in the last ten years (FAO 2016). However, cropping areas for sugarcane could not be increased indefinitely due to the limited farming land areas and competition with food crops. In order to sustainably improve sugarcane production, breeding sugarcane cultivars with high performance for yield components is critical. Thus, evaluating yield components in sugarcane germplasm and dissecting the genetic basis of causal sequence variants are of great importance to fulfill this strategy by providing tools and genetic resources.

The *Saccharum* genus consists of two wild species (*S. spontaneum* and *S. robustum*) and four cultivated species (*S. officinarum, S. barberi, S. sinence*, and *S. edule*) (Daniels and Roach 1987). All the *Saccharum* species are readily intercrossed except sterile S. *edule*. Modern sugarcane hybrids were primarily derived from inter-species hybridization between *S. spontaneum* (2n = 40-128, x = 8) and *S. officinarum* (2n = 80, x = 10) followed by several backcrosses with *S. officinarum*. Therefore, modern sugarcane hybrids have a chromosome number of 100-130 with approximately 80% chromosomes from *S. officinarum*, 10% chromosomes from *S. spontaneum*, and 10% recombinant chromosomes (D'Hont et al. 1996), and an estimated genome size of approximately 10 Gbp (D'Hont 2005). Due to the nature of huge genome size and high polyploidy level (Primarily aneuploidy), genetic studies of sugarcane have been extremely challenging with relatively slow progress compared with many other crops.

Modern sugarcane cultivars could be traced back to only a handful of *S. spontaneum* and *S. officinarum* ancestor clones (Acevedo et al. 2017; Deren 1995; Lima et al. 2002). This narrow genetic base hampered the sugarcane cultivar improvement on yield and tolerance. Therefore, maintenance, characterization, and utilization of genetic diversity in sugarcane germplasm collection are critical to broaden the genetic background of sugarcane for crop improvement. The “world collection of sugarcane and related grasses” (WCSRG), which assembled a significant amount of genetic diversity, is an essential resource for sustained sugarcane cultivar improvement (Nayak et al. 2014; Todd et al. 2014). The collection has ~ 1,200 accessions from 45 countries, including *Saccharum* germplasm and closely related grass species with the most abundant being *S. spontaneum, S. officinarum*, and interspecific hybrids. This collection preserves gene resources that could be used to presumably improve yield, fiber and abiotic and biotic stresses in sugarcane breeding programs. A core germplasm collection was created by selecting representative accessions from the WCSRG (Nayak et al. 2014; Todd et al. 2014), which showed significant amount of natural phenotypic variations, and can be valuable resources for identifying desirable alleles contributing to yield and tolerance for sugarcane improvement.

Genome-wide association study (GWAS) has been one of the most powerful tools to discover causal variants for complex traits in human, animal and plant systems. Compared with traditional linkage mapping strategy performed in bi-parental populations, GWAS explores historical and evolutionary recombination events at the population level, and thus achieves a high resolution to the gene level (Zhu et al. 2008). In addition, multiple genes and alleles involved in the same traits could be investigated and identified in a natural diversity panel. With continually decreasing cost for the next generation sequencing (NGS) technology, GWAS has been widely applied in complex trait dissection in the genomic era. However, GWAS might not be as promising in polyploid sugarcane (With up to 12 sets of chromosomes) as in diploids. There are several primary challenges that could hinder the use of GWAS in sugarcane. For example, 1) relatively high sequencing depth needed to call sequence variants due to their large genome sizes (~ 10 Gbp); 2) dosages of markers were difficult to be determined especially for single dose markers; and 3) most GWAS models and software are designed for diploids, in which dosages and gene actions could not be accounted for analyses. Hence in most cases, GWAS in polyploids treated molecular markers as diploids, which might not reflect the actual genetic effects. Since the first GWAS was conducted (Haines et al. 2005), a few GWAS have been performed in sugarcane (Débibakas et al. 2014; Gouy et al. 2015; Racedo et al. 2016; Wei et al. 2006, 2010). However, all of them were challenged by the difficulties mentioned above. GWASpoly in R is a software tailored for polyploids, which could model different types of polyploid gene actions in GWAS, including additive, simplex dominant, and duplex dominant (Rosyara et al. 2016). This software has been tested in a simulated tetraploid population, and successfully applied in potato, sunflower, orchardgrass and wheat for GWAS (Berdugo-Cely et al. 2017; Bock et al. 2018; Phan et al. 2018; Rosyara et al. 2016; Zhao et al. 2017). However, this software has not been utilized for GWAS in sugarcane.

A run of homozygosity (ROH) is a continuous homozygous segment of the genome in an individual due to haplotypes inherited from common ancestors in the past (Ceballos et al. 2018). ROHs capture population history of a species, such as inbreeding, change of population size, admixture and so on. ROH calling relies on high-density genome-wide markers (Ceballos et al. 2018). With the development of NGS technologies, large numbers of single polymorphisms (SNPs) can be generated at a relatively low price, making it achievable to perform ROH analysis to capture the genomic regions contributing to inbreeding and thus to assess the breeding history and to identify the genetic components for trait selection. ROHs are widely applied in human and animal genetics (Ceballos et al. 2018; Christofidou et al. 2015; Peripolli et al. 2018; Sud et al. 2015; Yang et al. 2015) for detecting inbreeding and loci associated with quantitative trait or disease. However, there is no report of ROHs in sugarcane genetic research.

In this study, we assembled a diversity panel consisted of 299 accessions based on phenotypic and genotypic data collected from the WCSRG, and nine sugarcane cultivars and breeding materials (Nayak et al. 2014; Todd et al. 2014). Nine yield components were evaluated in the diversity panel in three replicates. We deeply sequenced the coding regions of the 308 accessions in the diversity panel, and identified large amounts of sequence variants with a confident dosage resolution. We performed GWAS using GWASpoly in sugarcane with marker dosages and gene actions taken into consideration. A total of 217 non-redundant sequence variants and 225 genes were significantly associated with the traits evaluated. In addition, for the first time, we identified 280 ROHs, indicators of inbreeding in the diversity panel, and found that ROHs were associated with two yield components. The results of this study not only provided the genetic bases of multiple yield components in sugarcane but also suggested important tools and deposited valuable genomics resources for the sugarcane community. The new methods and concepts in this study shed light on research in sugarcane and polyploid species.

## Materials and methods

### Plants in the association panel

A core sugarcane diversity panel consisting of 299 accessions collected were selected from the WCSRG, maintained at the USDA-ARS Subtropical Horticulture Research Station, Miami, Florida, to represent the wide genetic diversity of the WCSRG (Nayak et al. 2014; Todd et al. 2014). Nine modern sugarcane hybrids frequently used as parental materials in the USA Florida sugarcane breeding program (Table S1) were also included in this study. In total, 308 accessions were established at USDA-ARS Sugarcane Field Station at Canal Point, FL in 2013 in a field nursery in a randomized complete block design with three replicates. Regular fertilizer and daily water were applied to maintain healthy plants in the nursery.

### Phenotyping and data treatment

During August 2014, nine yield components were measured for the sugarcane diversity with three replicates on ratoon plants including stalk diameter, leaf width, leaf length, stalk number, internode length, brix, total weight, dry weight and water content (Todd et al. 2017). In brief, brix, stalk number, stalk diameter, internode length, leaf width, and leaf length of the leaf at the top visible dewlap were recorded for each plant before harvesting. Immediately prior to harvest, three or more stalks (To roughly equal 1 kg dry mass) were shredded, and the total weight was recorded per pot. Shredded cane was subsequently dried at 60 °C, and re-weighted to determine dry weight. Water content was calculated by subtracting dry weight from total weight.

The normality of the trait distribution was checked by their phenotype distribution. For the traits not following normal distribution, a Box Cox transformation was performed using MASS package in R (Ripley et al. 2013) to satisfy model assumptions. A pairwise Pearson correlation among yield components was calculated using the ‘Hmisc’ package in R3.0.2 (Harrell 2018). The total correlation coefficients of yield components to sugarcane dry weight and sugarcane content (Brix) were participated into direct and indirect effects following the method (Dewey and Lu, 1959), and the computation was conducted using ‘lavaan’ and ‘semPlot’ package in R (Epskamp 2015; Rosseel 2012). Analysis of variance was performed using the ‘nlme’ package in R (Pinheiro et al. 2009). Broad sense heritability (*H*^2^) was estimated using the formula:

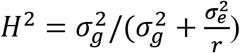

Where 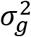 is the genetic variance, 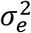 is the residual variance and *r* is the number of replicates. For each trait, the best linear unbiased predictor (BLUP) was estimated by modeling genotypes as random effects and replicates as fixed effects using *lme4* package in R (Bates et al. 2014), and the computed BLUPs were used for GWAS.

### Genotyping

Young leaves were collected from each accession, and the DNA was extracted by using the cetyltrimethyl ammonium bromide (CTAB) method (Wang et al. 2010). The target enrichment sequencing (TES) of each accession was conducted (Song et al. 2016). In brief, a set of 60 thousand 120-bp oligonucleotides probes were designed, and used for TES. The captured DNA libraries were prepared following the product manual of Agilent SureSelectXT reagents (Agilent Technologies, Santa Clara, CA), and then were indexed and cleaned using Ampure beads, quantified by qPCR using KAPA Library Quantification Kits (KAPA Biosystems.com) and sequenced in paired-end mode (2 x 100 bp) by Illumina HiSeq 2000.

The raw reads were sorted into reads per library according to their barcode sequences and then the barcodes were removed from raw data. The paired-end reads were trimmed from ends in pairs based on PHRED score of 20 using Trimommatic v0.36 (Bolger et al. 2014). Reads longer than 50 bp were aligned to sorghum genome v3.0 (Paterson et al. 2009). BWA-mem (Li 2013) with default settings was used for alignment. Uniquely mapped reads were extracted from sequence alignment BAM files, and used for sequence variant calling. SNPs were called by using Unified Genotyper implemented in Genome Analysis Tool Kit v3.30 (McKenna et al. 2010). The raw SNPs were subjected to a series of filtering for each individual accession: 1) mapping quality for SNPs > 30; 2) base quality for SNP sites > 20; 3) for a heterozygous genotype, at least two reads for minor allele were required; otherwise, missing genotypes were assigned; 4) for a given SNP locus with homozygous genotype, minimum reads were calculated according to ploidy of accessions to ensure single dose SNPs were not called as homozygotes (Yang et al. 2017, 2018). Based on the calculation, minimum reads of 5, 11, 17, 23, 29 and 35 were required for ploidy at 2, 4, 6, 8, 10 and 12 for homozygous genotype calling, respectively; otherwise, missing genotypes were assigned; 5) for a given SNP locus with heterozygous genotype, sequencing depth required to separate all SNP genotypes with a probability higher than 95% at determined ploidy, which were 31, 47, 70, 90 and 108 for ploidy at 4, 6, 8, 10 and 12, respectively. After the stringent filtering, genotypes of SNPs were scaled to the highest ploidy (12) according to ratio of reads for respective alleles. InDels were called using the same procedures as SNPs described above.

### Genome wide association study

We used discriminant analysis of principal components (DAPC) implemented in the adegent package for R to infer the population structure of the accessions in this study (Jombart, 2008, 2010). *k*-means clustering was used to identify groups, where the best *k* minimized the Bayesian Information Criterion (BIC) and was thus used for GWAS. GWASpoly is tailored for autopolyploids based on the Q + K mixed model, and could model gene actions for polyploids (Rosyara et al. 2016). Six models were used for GWAS by using GWASpoly, including general model, additive model, two simplex dominant models (1-dom-ref and 1-dom-alt), diplo-general model, and diplo-additive model. In additive model, effect of markers was proportional to the dosage of minor allele, while heterozygotes were equivalent to one of the homozygotes for simplex model (See (Rosyara et al. 2016) for detail). Diplo- means that genotypes of markers were diploidized for the model. Sequence variants for GWAS were further filtered with a call rate ≥ 95% and minor allele frequency (MAF) ≥ 0.05. Only samples with less than 20% missing data were included for the GWAS. We performed GWAS runs for nine yield components in the full set of the sugarcane diversity panel (with 300 accessions) in addition to several subsets separately, including a subpopulation 1 (Sacc) with 273 accessions from *S. spontaneum, S. officinarum*, modern *S*. hybrids and ancient S. hybrids, a subpopulation 2 (Spon) with 100 accessions from *S. spontaneum*, a subpopulation 3 (Hybrid) with 173 accessions from *S. officinarum*, modern *S*. hybrids and ancient S. hybrids, and a subpopulation 4 (P8) with 162 accessions at the same ploidy level of 8. The annotation of candidate genes with GWAS hits were obtained from the sorghum genome annotation (Paterson et al. 2009), and their domains were annotated using InterProScan 5.0 (Jones et al. 2014).

### Identification of runs of homozygosity

SNPs with confidence separating homozygote and heterozygote (After step 1, 2 and 3 filtering in ‘genotyping’ part) were used for identification of ROH. Additional quality control for SNPs included: individual missing rate < 10%, SNP call rate ≥ 90%, and minor allele frequency < 1%. ROHs were identified in each individual using Plink v1.90b3 using default settings by fixed scanning windows (Purcell et al. 2007). Specifically, the parameters used to define and identify a ROH were: 1) a sliding window contained 50 sequence variants with at most 1 heterozygous call and 5 missing calls; 2) for a variant to be eligible in a ROH, the hit rate of all scanning windows containing the variant must be at least 0.05; 3) maximum gap between two consecutive homozygous variants was 1,000 Kbp; 4) a ROH had a minimum length of 1,000 Kbp, and contained at least 100 sequence variants, and at least one variant per 50 Kbp on average. Linear regression was used to study association between ROHs and phenotypes with control for population structures. Associations were performed for total number of ROHs (NROHs), sum total length of ROHs (SROHs), and average length of ROHs (AROHs), separately.

### Data availability

The cleaned target enrichment sequencing reads generated in this study were deposited into NCBI Short Reads Archive with an accession number of SRP132365. The phenotype, genotype and population structure used in GWAS were deposited into Gatorcloud (https://bit.ly/2OcwsN4). The rest intermediate analysis data and the plant materials will be available upon request.

## Results

### Correlation and path coefficient analysis of yield components

All the traits were normally distributed except stalk number (Figure S1), which was further transformed. According to pairwise Pearson’s correlation coefficients for the nine traits, five pairs of traits were found to be highly correlated (Correlation coefficients r ≥ 0.6 or r ≤ −0.6) (*P* < 0.0001) (Table 1). Specifically, leaf width was positively correlated with stalk diameter (r = 0.82) but negatively correlated with stalk number (R = −0.62), and stalk diameter had negative correlation with stalk number (R = −0.69), reflecting there was a competition for carbon resources between stalk diameter and stalk number. Based on the correlations, leaf width seems to be more important to determine sugarcane stalk diameter and stalk number compared with leaf length. As expected, sugarcane total weight, dry weight and water content were highly correlated (With a correlation coefficient 0.78 between dry weight and total weight, and 0.92 between total weight and water content) (Table 1). We further partitioned the correlation into direct effects and indirect effects to understand the inter-relationships among these yield components. Stalk number and internode length had high direct effects on sugarcane yield (Dry weight) (Table 2), which were consistent with the results of overall correlation coefficients (Table 1). Moreover, indirect effects were relatively low for both stalk number and internode length. The results implied that selection on high stalk number and internode length would be beneficial for increasing sugarcane yield. On contrary, leaf width had negative direct and indirect effects on sugarcane yield, while stalk diameter had negative indirect effects (Table 2), which should be selected against for yield improvement. The situation is much complex for sugar content (Brix) because all the yield related traits, such as stalk diameter, stalk number and leaf width, had both relatively high negative and positive effects (Direct or indirect) on sugar content (Table 3). The overall correlation should be considered in breeding material selection. To breed sugarcane cultivars with high sugar content, the results suggested that genotypes with high stalk diameter, leaf width but low stalk number should be selected.

**Table 1.**
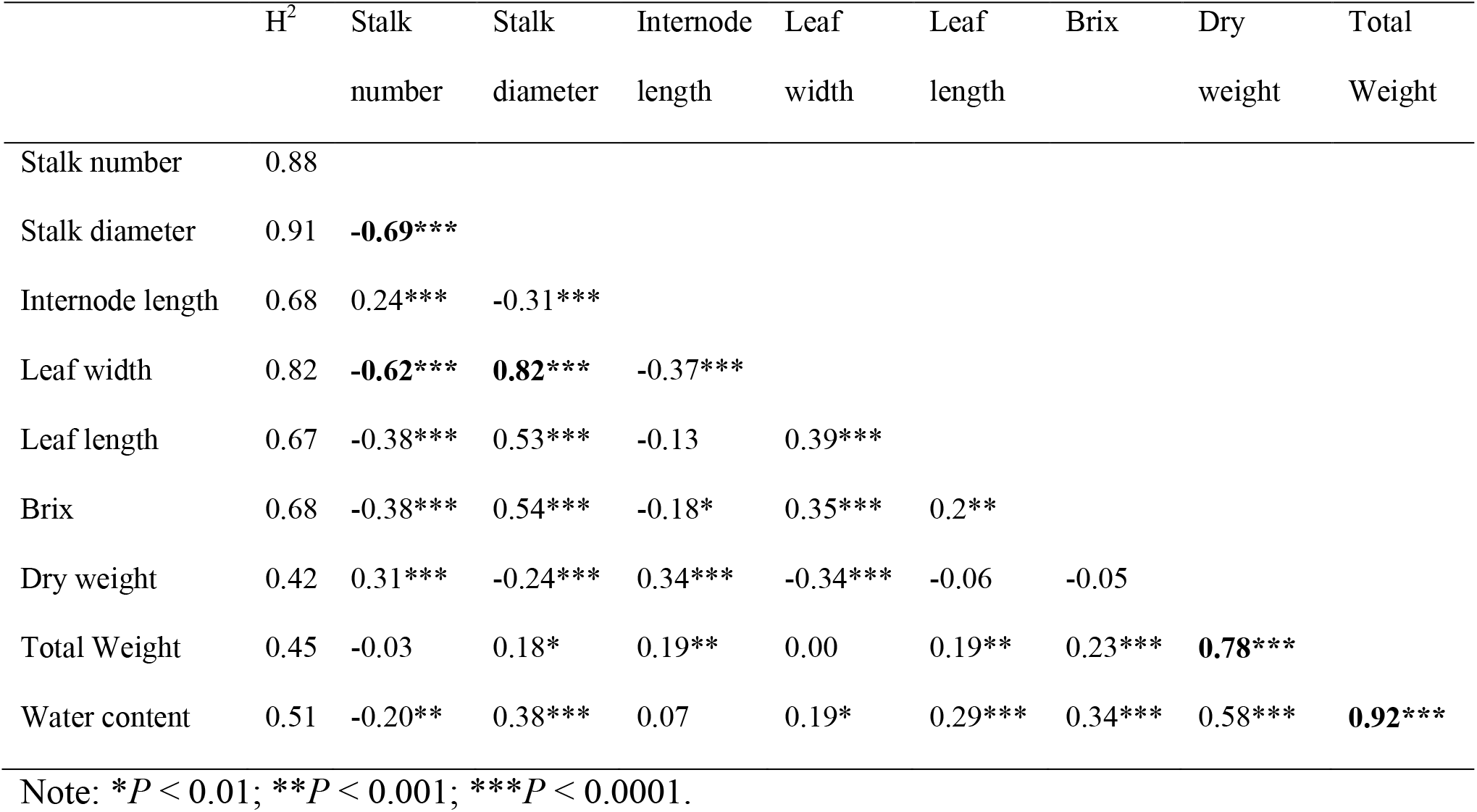
Pair-wise Pearson’s correlation coefficients and broad sense heritability (*H*^2^) for the nine yield components evaluated in the sugarcane diversity panel.

**Table 2.**
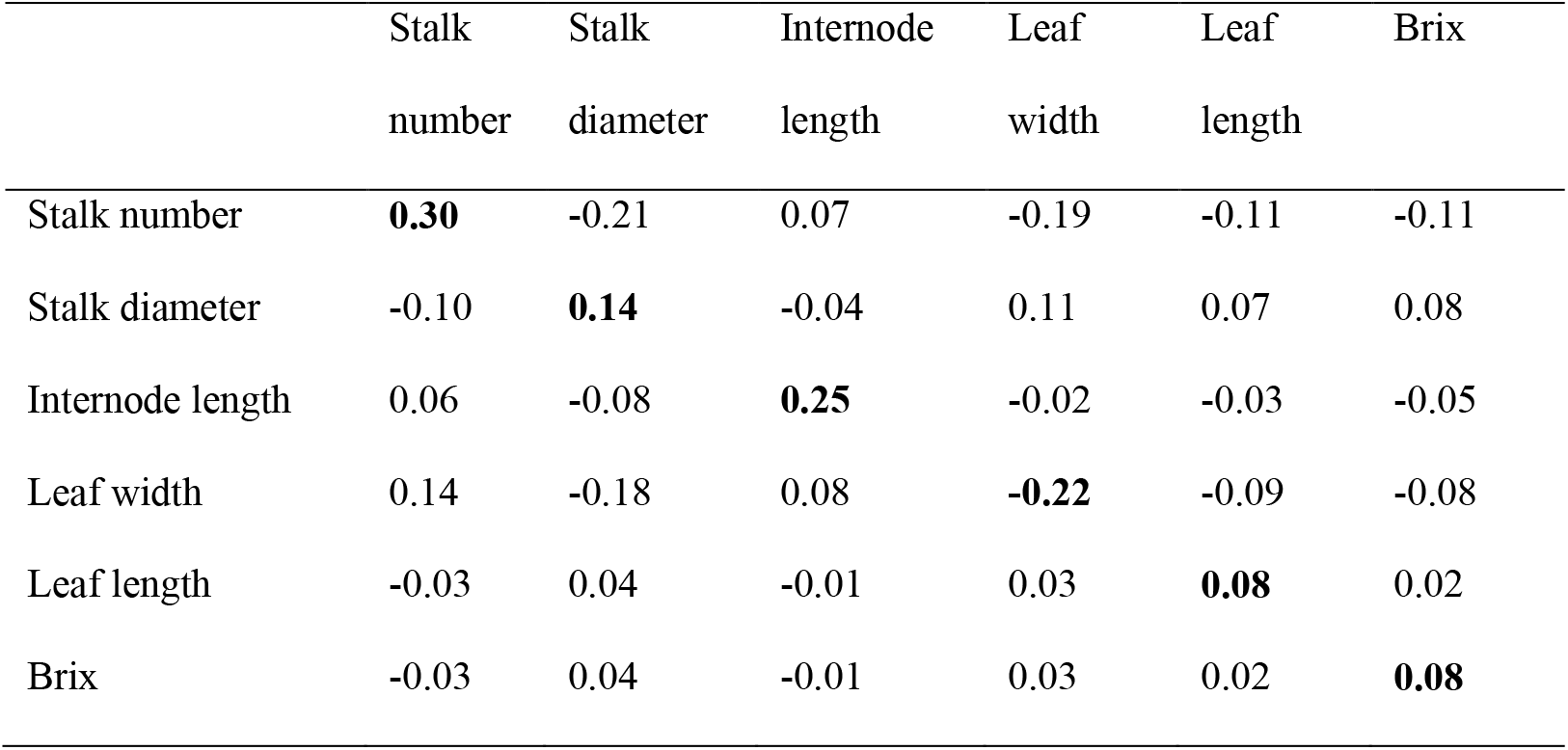
Path coefficients showing direct (Diagonal) and indirect effects (Non-diagonal) of six yield components on sugarcane dry weight.

|  | Stalk<br>number | Stalk<br>diameter | Internode<br>length | Leaf<br>width | Leaf<br>length | Brix |
| --- | --- | --- | --- | --- | --- | --- |
| Stalk number | **0.30** | -0.21 | 0.07 | -0.19 | -0.11 | -0.11 |
| Stalk diameter | -0.10 | **0.14** | -0.04 | 0.11 | 0.07 | 0.08 |
| Internode length | 0.06 | -0.08 | **0.25** | -0.02 | -0.03 | -0.05 |
| Leaf width | 0.14 | -0.18 | 0.08 | **-0.22** | -0.09 | -0.08 |
| Leaf length | -0.03 | 0.04 | -0.01 | 0.03 | **0.08** | 0.02 |
| Brix | -0.03 | 0.04 | -0.01 | 0.03 | 0.02 | **0.08** |

**Table 3.**
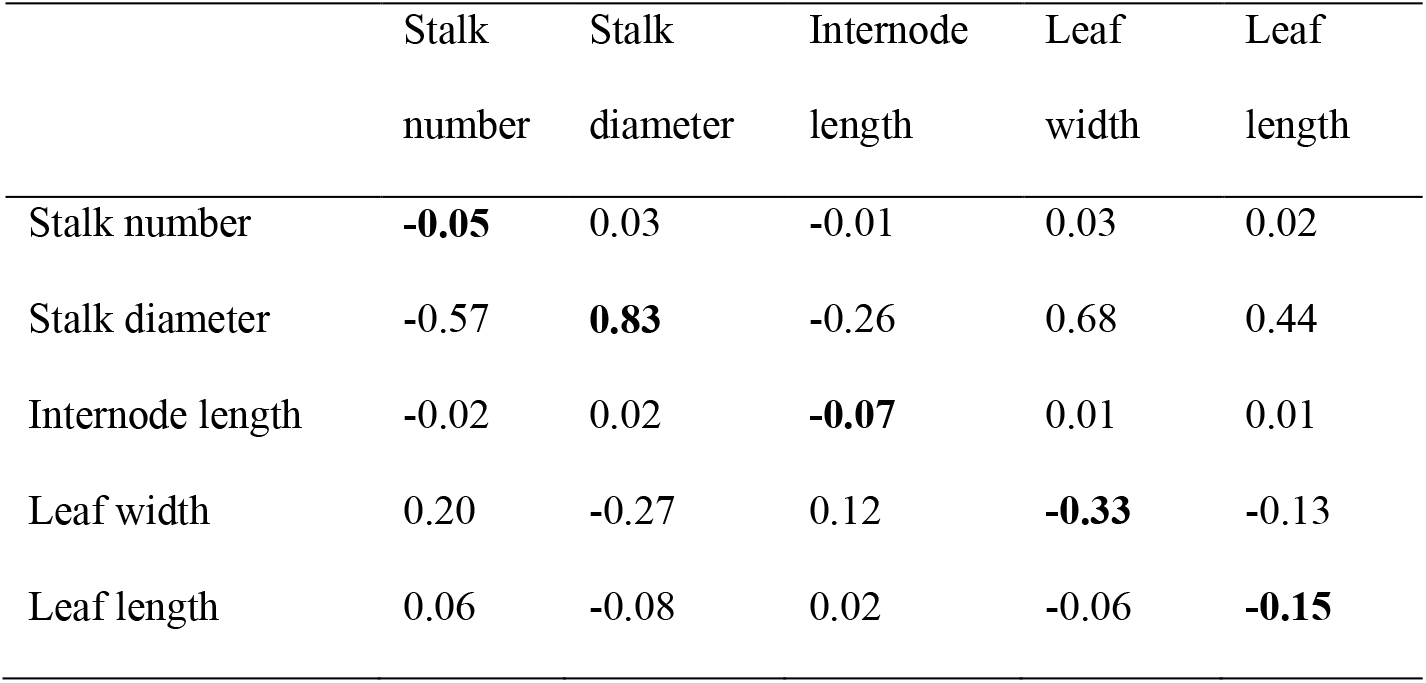
Path coefficients showing direct (Diagonal) and indirect effects (Non-diagonal) of five yield components on sugar content (Brix).

|  | Stalk<br>number | Stalk<br>diameter | Internode<br>length | Leaf<br>width | Leaf<br>length |
| --- | --- | --- | --- | --- | --- |
| Stalk number | **-0.05** | 0.03 | -0.01 | 0.03 | 0.02 |
| Stalk diameter | -0.57 | **0.83** | -0.26 | 0.68 | 0.44 |
| Internode length | -0.02 | 0.02 | **-0.07** | 0.01 | 0.01 |
| Leaf width | 0.20 | -0.27 | 0.12 | **-0.33** | -0.13 |
| Leaf length | 0.06 | -0.08 | 0.02 | -0.06 | **-0.15** |

The broad sense heritability (*H*^2^) ranged from 0.42 (Dry weight) to 0.91 (Stalk diameter) for the nine yield components (Table 1). The total weight, dry weight and water content had relative low broad sense heritability, probably because they were complex traits combining impacts from multiple yield components with significant environmental effects. Overall, the estimated broad sense heritability indicated that these yield components were largely controlled by genetic factors. Therefore, the nine yield components were used for GWAS analysis. Though some traits were highly correlated, GWAS may identify the same markers to be associated with the correlated traits. To maximum discovery of associations, we conducted GWAS for each of the nine traits separately without considering the correlation first. However, after the GWAS, we summarized and presented a unique set of associated markers subsequently.

### Sequence variants captured by target enrichment sequencing

Overall, 8.0 billion clean reads were generated after sequencing the target regions of the 308 accessions with 7.4 billion (92.1%) of the clean reads mapped to the sorghum genome, and 7.3 billion (90.7%) uniquely mapped. The average sequence depth of the whole diversity panel was 99x in a range of 9x to 375x. Using the uniquely mapped reads, 25.3 million raw SNPs were identified, of which 9.9 million (39.2%) were SNPs with high quality after the stringent sample-by-sample filtering (Table 4). The 9.9 million SNPs included 8.6 million bi-allelic SNPs and 1.3 million multiple-allelic SNPs. Majority of the bi-allelic SNPs (90.2%) were rare variants (MAF < 0.05), and thus were removed from further analysis. With a SNP call rate ≥ 95%, 65 thousand SNPs (0.7% of the 9.9 million high quality SNPs) were included as markers for GWAS. Similar patterns were observed for InDels with 880.0 K high quality InDels after sample-by-sample filtering, in which 130.2 K (14.8%) were multiple allelic, and majority of the bi-allelic InDels (562.9 K, 75.1% of the 880.0 K) were rare variants (Table 4). With the same call rates as applied to SNPs, 9.9 K InDels were included as markers for GWAS. Therefore, a total of 74.9 K DNA markers, including 65.0 K SNPs and 9.9 K InDels, were used for subsequent GWAS.

**Table 4.**
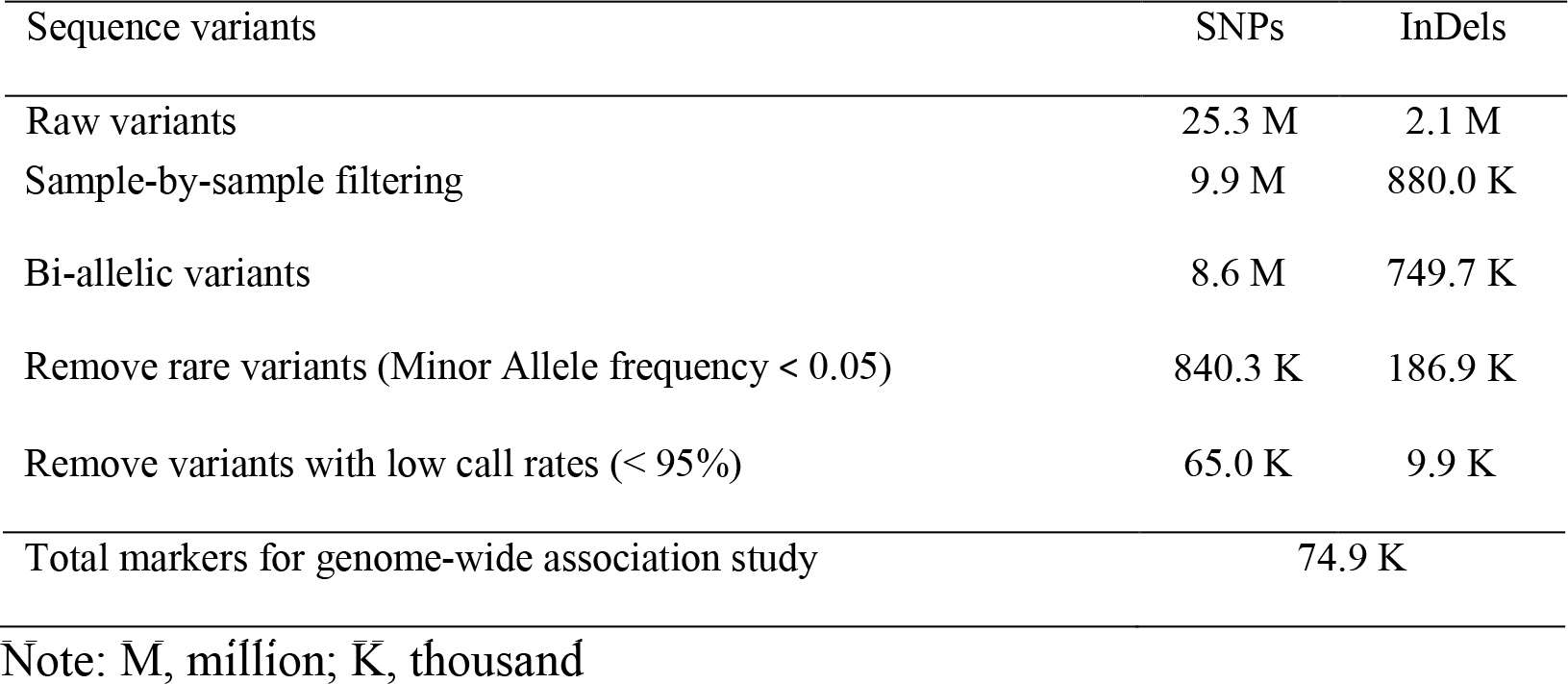
Summary of single nucleotide polymorphisms (SNPs) and insertions and deletions (InDels) identified in the sugarcane diversity panel.

According to the sorghum genome, the majority of the DNA markers used for GWAS were in genic regions (67.5 K, 90.1%) targeting 10,458 gene models. The number of markers per chromosome ranged from 2,758 (Chr05) to 12,836 (Chr01), with an average marker distance of 9,116 bp. Given the large block of linkage disequilibrium in sugarcane (~ 5 cM, (Raboin et al. 2008)), the marker density in this study should be high enough for an effective GWAS.

### Inferring population structure

We carried out population structure analysis with DAPC using 13.8 K SNPs, which were evenly distributed according to the sorghum genome (One SNP per 10 Kbp) and had call rates ≥ 95% in the sugarcane diversity panel. We selected *k* = 6 representing number of population groups in the diversity panel, which was within the curve minimizing the BIC value (Figure 1A). Among the 308 accessions, four groups belonging to *Saccharum* were identified with *S. spontaneum* group including 101 accessions, *S. officinarum* group including 47 accessions, modern *S*. hybrids group including 114 accessions, and ancient S. hybrids group (S. *barberi* and *S. sinence*) including 17 accessions (Figure 1B; Table S1). In addition to *Saccharum* spp. groups, two non*-Saccharum* groups were identified in the diversity panel as non-*Saccharum* 1 and non-*Saccharum* 2 with 10 and 19 accessions, respectively (Figure 1B, Table S1). Based on WCSRG records, *non-Saccharum* 1 most likely belonged to *Miscanthus*, while *non-Saccharum* 2 belonged to *Erianthus* (Table S1). Due to low sequencing depth (On average 24.3), eight accessions, including five accessions from modern S. hybrids, two from *non-Saccharum* 2, and one from *S. spontaneum*, with missing sequence variants > 20%, were removed from further analysis. As a result, 300 accessions from the sugarcane diversity panel were included for subsequent GWAS.

**Figure 1.**
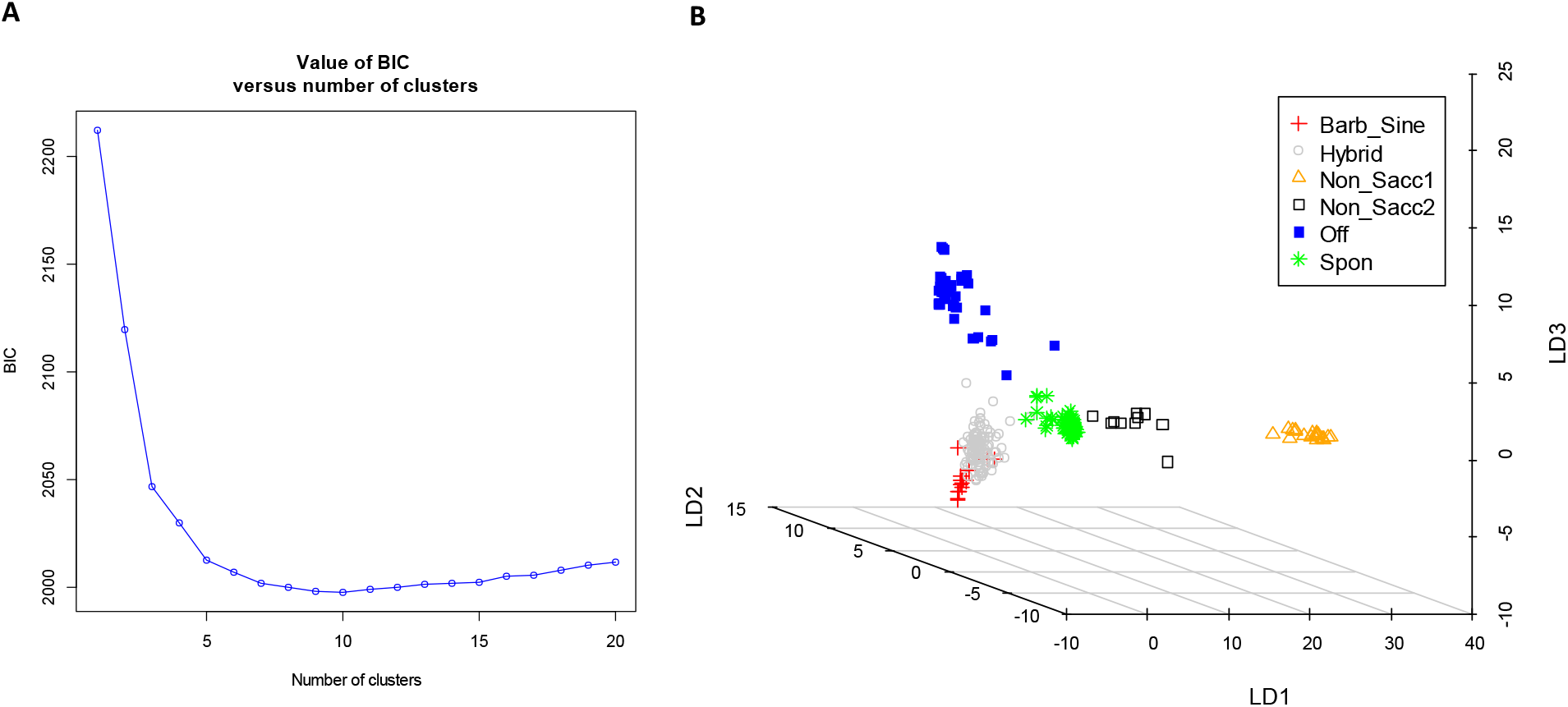
Population structure analysis of the sugarcane diversity panel based on 13.8 K single nucleotide polymorphisms using discriminant analysis of principal components. (A) Bayesian Information Criteria (BIC) vs. number of clusters in k-means clustering. (B) Projection of the sugarcane diversity panel using the first three linear discriminants (LDs). Spon = *S. spontaneum*; Off = *S. officinarum*; Hybrid = modern *S*. hybrid; Barb = *S. barberi*, Sine = *S. sinence*; Non sacc = Non *saccharum*.

### Genome-wide association study

Using a stringent threshold (*P* < 0.05 after Bonferroni correction), 217 non-redundant markers (191 SNPs and 26 InDels) were identified to be significantly associated (Figure 2, Figure S2, S3, S4, S5, S6, and Table S3). The 217 associated markers included 128, 77, 30, 13 and 58 from the full population, subpopulation 1, 2, 3, and 4, respectively (See method part), with some associations overlapped (Table 5). The genome-wide association rate was 0.55% (217/74,935 = 0.29%). The number of associated markers according to the sorghum gene models was 33 in intergenic regions, 60 in introns, 48 in untranslated regions, and 66 in coding regions. For the markers in coding regions, three had high effects (Stop gain or frameshift variant), 25 had moderate effects (Non-synonymous mutations) and 38 had low effects (Synonymous mutations). Only 28 markers (12.9%) led to new alleles with likely different functions, while the rest did not change the original gene function, which might perform the roles as regulatory elements or associate with other causal genes. Among the 217 associated sequence variants, 109 were detected in at least two GWAS runs or two models reflecting certain level of consistence in the trait-marker association. There were nine markers associated with more than two different traits. Specifically, six markers (2P8663239, 3P66737730, 3P66744355, 4P2535140, 5ind60876916, 10P55668582) were associated with both stalk diameter and leaf width, two markers (3P61470337 and 7P2769299) with water content and total weight, and one marker (3P63471814) with dry weight, water content, and total weight (Table S3). These single markers associated with multiple traits indicated that either these loci located in or be associated with pleiotropic genes, or these traits were highly correlated biochemically.

**Figure 2.**
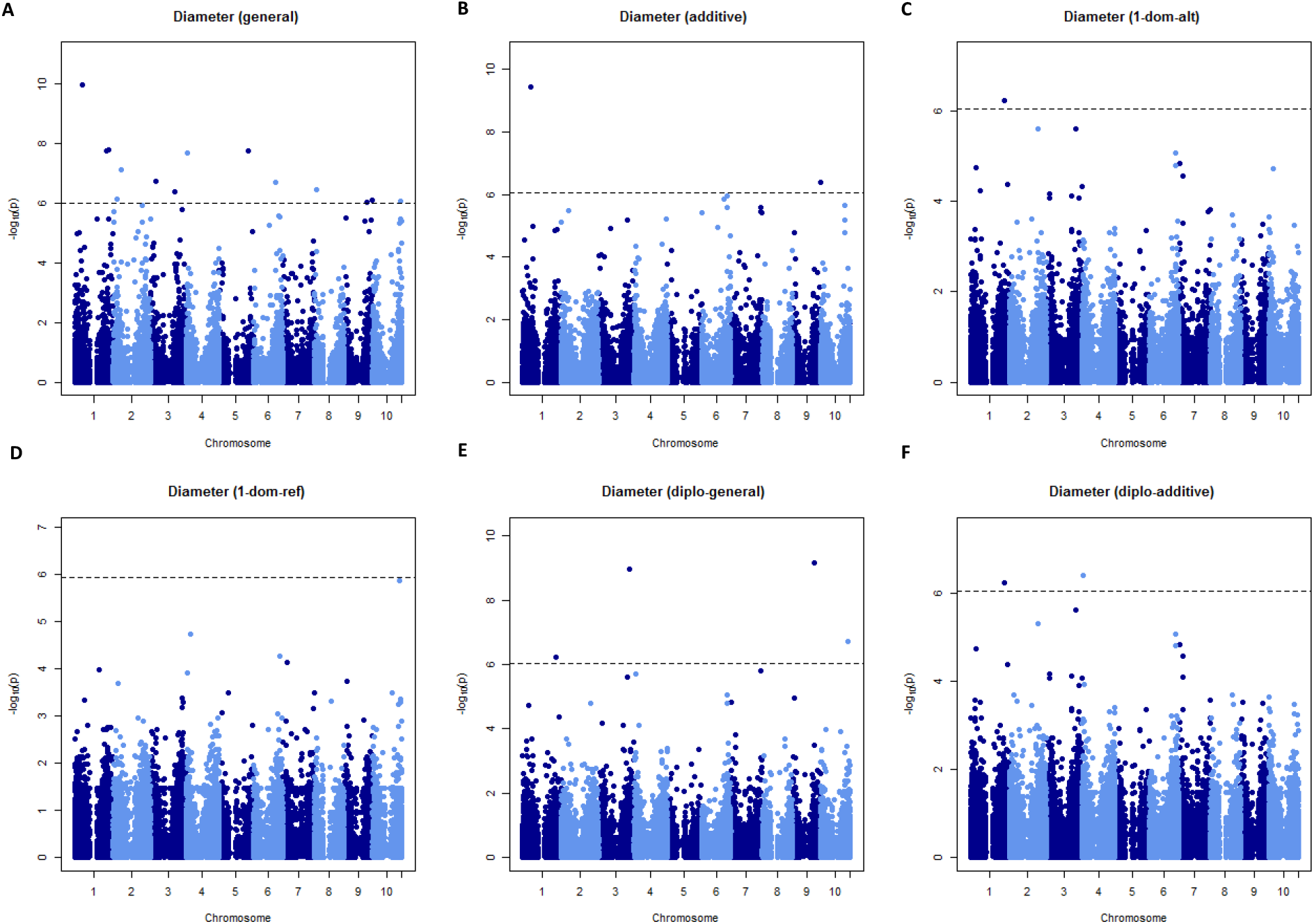
Example of significant marker-trait associations identified in the sugarcane diversity panel (Full set) for stalk diameter (Diameter) using general model (A), additive model (B), 1-dom-alt model (C), 1-dom-ref model (D), diplo-general model (E), and diplo-additive model (F).

**Figure 3.**
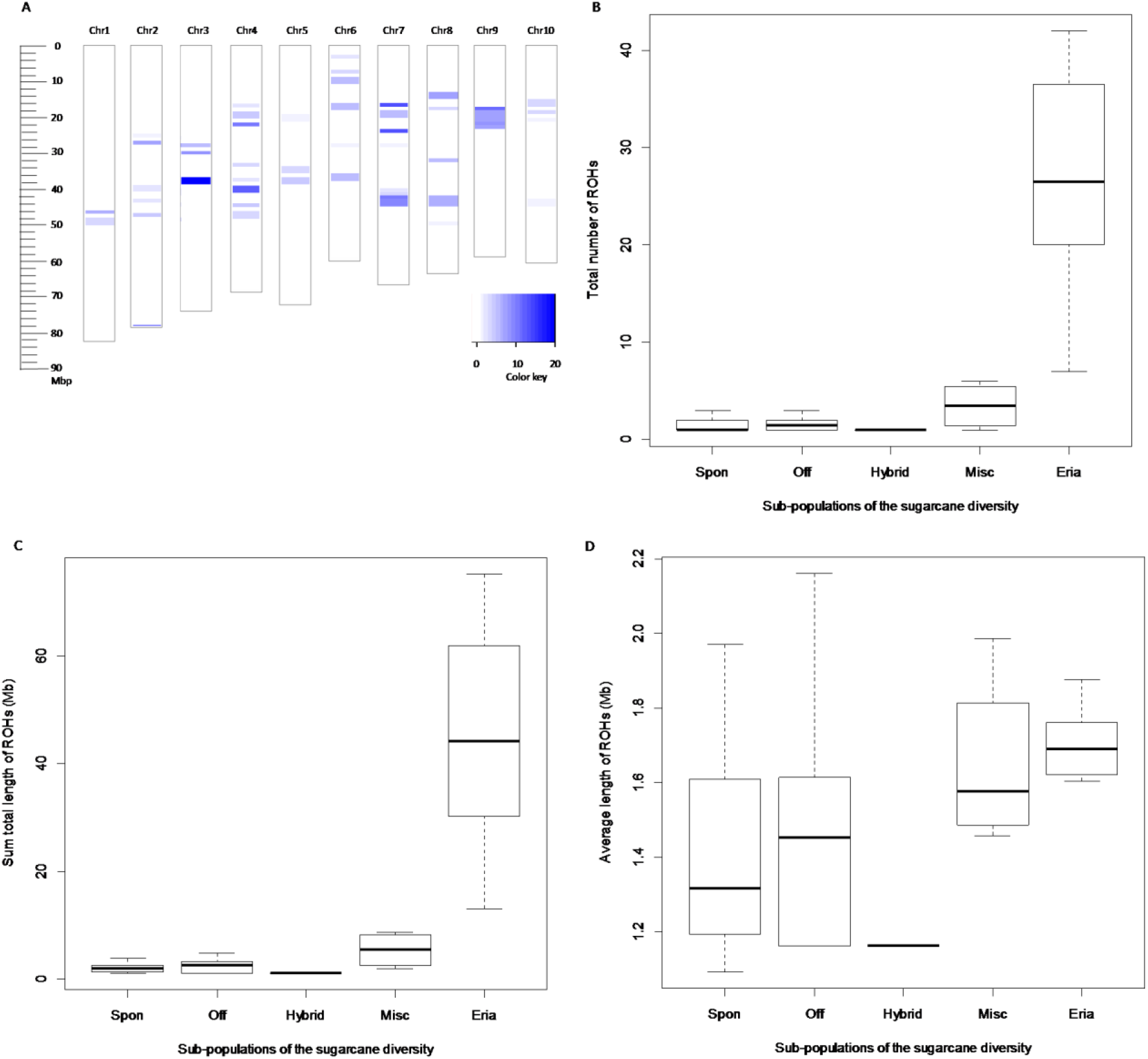
Summary of runs of homozygosity (ROHs) in the sugarcane diversity panel. A) Distribution of common ROHs according to the sorghum genome; B) Total number of ROHs; C) sum total length of ROHs; D) average length of ROHs. Spon = *S. spontaneum*; Off = *S. officinarum*; Hybrid = modern *S*. hybrid; Misc = *Miscanthus*; Eria = *Erianthus*.

**Table 5.** Number of associated sequence variants identified in different runs and models of genome-wide association study.

| Runs<sup>a</sup>/Models | General | Additive | 1-dom-alt | 1-dom-ref | diplo-general | diplo-additive | Total |
| --- | --- | --- | --- | --- | --- | --- | --- |
| Full | 99 | 9 | 25 | 14 | 35 | 27 | 128 |
| Sacc | 71 | 2 | 2 | 2 | 10 | 2 | 77 |
| Spon | 31 | 1 | 0 | 0 | 0 | 0 | 30 |
| Hybrid | 13 | 0 | 0 | 0 | 0 | 0 | 13 |
| P8 | 45 | 6 | 9 | 7 | 10 | 7 | 58 |

Note: <sup>a</sup> GWAS runs in Full set and several subests. Full = all accessions in the diversity panel; Sacc = 273 accessions from *S. spontaneum*, *S. officinarum*, modern *S. hybrids* and ancient *S. hybrids*; Spon = 100 accessions from *S. spontaneum*; Hybrid = 173 accessions from *S. officinarum*, modern *S. hybrids* and ancient *S. hybrids*; and P8 = 162 accessions with the same ploidy level (Ploidy = 8).

Among the 217 non-redundant associated sequence variants, 184 were in genic regions targeting 169 gene models, and 33 in intergenic regions according to the sorghum genome (Table S3). To further investigate the genes involved in the associations, a total 225 non-redundant genes were obtained by retrieving the gene models hit by the associated markers and flanked by associated markers, including 167 genes with GWAS direct hits in gene models, 56 were extracted from genomic regions flanked by GWAS hit markers, and two genes were shared between the two categories (Table S3). The average number of candidate genes was 25 for the evaluated yield components with a range of two (Leaf length) to 89 (Leaf width). Details of associated markers and genes for the nine evaluated yield components were deposited in Tables S2 and S3. Based on annotations and domain analysis, among the associated genes, 10 were involved in sugar metabolism, including one (*Sobic.009G245000*) significantly associated with stalk diameter, one (*Sobic.006G278500*) with dry weight, two (*Sobic.002G279900* and *Sobic.009G063400*) with leaf width, and six (*Sobic.001G466900, Sobic.001G529600, Sobic.002G241100, Sobic.003G180100, Sobic.009G233200* and *Sobic.009G073400*) with internode length (Table S3). These genes encoded enzymes such as sugar-related synthase, hydrolases and transferase, important for carbon participation regulation, thus contribute to the biomass growth. We further searched enriched Gene Ontology (GO) categories for the 225 non-redundant genes, and genes associated for each component separately. The biological process for signal transduction by phosphorylation was significantly enriched in the genes associated with stalk diameter (*P* < 0.05 adjusted with false discovery rate (FDR)) (Yi et al. 2013), indicating signal transduction might be critical for determining stalk diameter sizes. We specifically focused on 27 associated genes with markers in coding regions that caused large effects (Frameshift or stop gained) (Three genes) or moderate effects (missense or disruptive in-frame deletion) (24 genes). The mRNA surveillance pathway was significantly enriched in the 27 associated genes (*P* < 0.05 adjusted with FDR) (Yi et al. 2013), indicating that ensuring fidelity and quality of messenger RNA has significant impacts on sugarcane yield components.

### Investigation of ROHs in the sugarcane diversity panel

With stringent filtering, 1.07 million SNPs and 211 accessions were used for ROH identification. The total genotyping rate of the SNPs was 97.2% in the 211 accessions. In total, 282 ROHs were characterized in 46 of the 211 accessions (20.4%). The distribution of ROHs was uneven and not correlated with SNP density, with a minimum of 12 ROHs on chromosome 1 and 5, and 57 ROHs on chromosome 7. There were 270 ROHs forming 50 pools of overlapping ROHs (The pool size ranging from 2 to 20 ROHs with 4 ROHs as the most popular pool size) (Figure 4A). The non-random distribution of ROHs indicated the occurrence of ROHs was most likely under selection during domestication and breeding processes. The sizes of ROHs ranged from 1.0 Mbp to 4.5 Mbp with an average of 1.5 Mbp. The most common ROH (Detected in 20 accessions) was located at chr3: 37153462-39398419 bp, harboring 25 genes. By searching gene ontology and pathway analysis (Lyne et al. 2015), genes were enriched in five biological processes and three cellular components in photosynthesis (*P* < 0.05 adjusted with FDR).

The ROH detection rate was 29.8% for *S. spontaneum* group (Out of 74 accessions), 27.8% for *S. officinarum* group (Out of 36 accessions), 0 for ancient S. hybrids group (out of 14 accessions), 2.7% for modern *S*. hybrids group (Out of 75 accessions), and 100% for both non-*Saccharum* groups (Table 6). The average NROH and SROH was 26.9 and 45.1 Mbp for non-*Saccharum* 2, which were significantly higher than that of any other sub-populations in this study (*P* < 0.001) (Figure 4B, 4C and Table 6). No difference was observed for the AROHs (Figure 4D, Table 6). The association analysis between the ROHs and the nine yield related traits showed that two yield components, total weight and dry weight, were associated with NROHs and SROHs significantly (*P* < 0.05). The negative coefficients of ROHs for both traits indicated that increase of genomic inbreeding in sugarcane had negative influence on these two yield components.

**Table 6.**
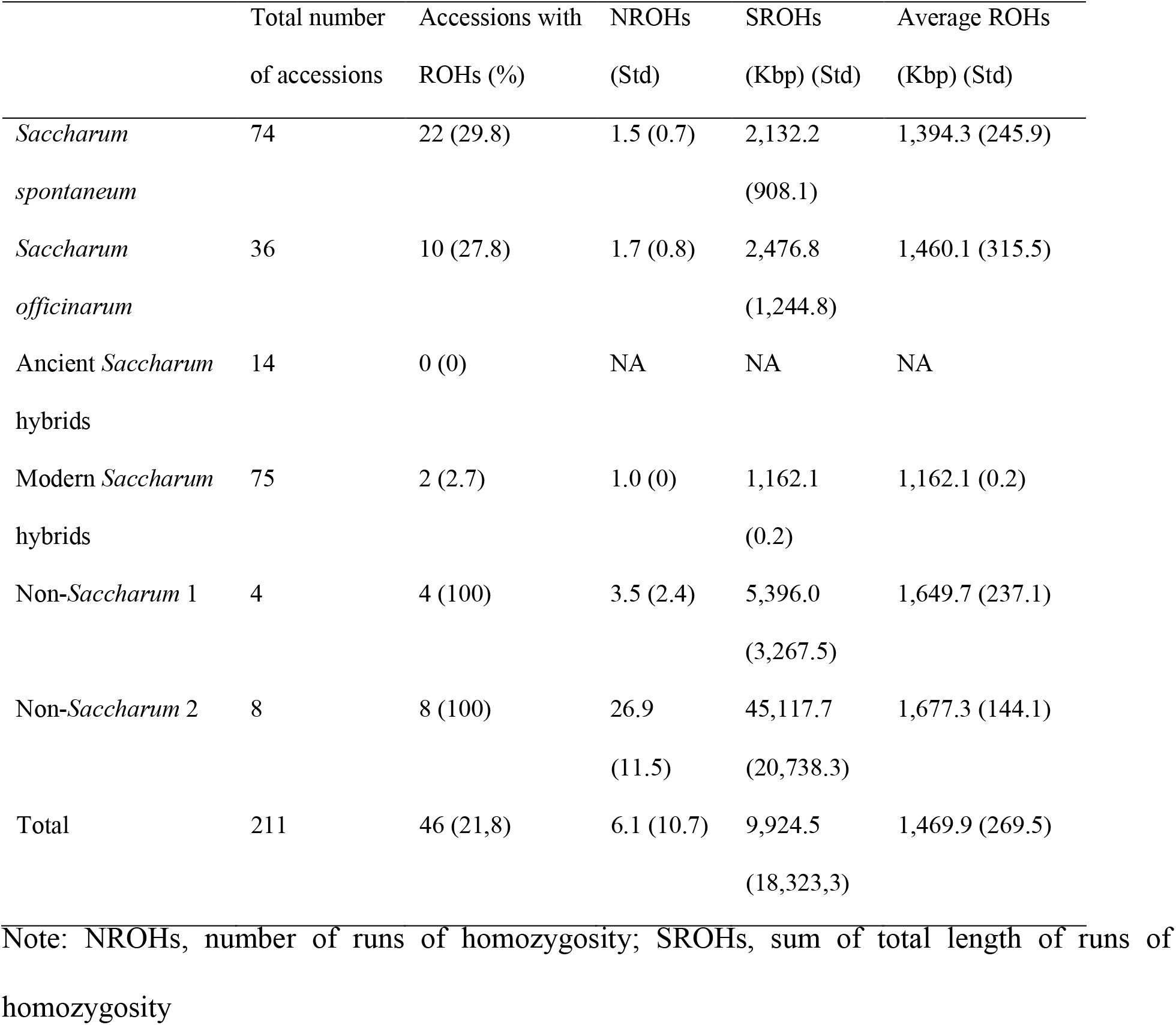
Statistical summary of runs of homozygosity (ROHs) identified in the sugarcane diversity panel.

## Discussion

Dissection of genetic loci controlling yield components is of great importance for sugarcane improvement. In this study, we performed GWAS analyses in a large representative diversity panel using high-density NGS-based DNA markers with dosages and allele interactions taken into consideration. We identified 217 non-redundant markers and 225 genes significantly associated with the nine yield components evaluated. In addition, for the first time, we investigated ROHs in sugarcane, and found that ROHs were negatively associated with total weight and dry weight. This research provided valuable and novel genomic resources and tools for sugarcane breeding programs, and empowered GWAS analyses in this polyploid species.

### High throughput genotyping sugarcane clones empowered by TES

Sugarcane is a highly polyploidy species with up to 12 sets of chromosomes, and has an estimated genome size of approximately 10 Gbp (D'Hont 2005). Due to the nature of huge and highly polyploidy genomes of sugarcane, intensive sequencing depth is required to accurately call genotypes with allele dosage resolution, which is the prerequisite to perform GWAS with allele dosages under consideration. In this study, we applied TES in a representative diversity panel for a high throughput genotyping. TES could capture a subset of sugarcane genome for sequencing, and therefore effectively investigate sequence variants at limited sequencing resources. This strategy to reduce genome complexity is very helpful for genomic studies in species like sugarcane with complex genomes. By selectively sequencing mostly the coding regions, we achieved an average sequencing depth of 99x, which is difficult to obtain using other type of NGS-enabled methods. At this sequencing depth, we could differentiate all genotypes of sequence variants in sugarcane, even for dodeca-ploid species with up to 13 possible genotypes (Yang et al., 2017). After applying a stringent sample-by-sample filtering, we obtained 74.9 K markers with an average marker distance of 9 Kbp, which is much higher than any of other published GWAS study in sugarcane. This powerful NGS-based technology has maximized the identification of markers associated with the traits evaluated. However, we still had only 39.2% of the raw SNPs pass our stringent quality filtering (Mainly due to sequence depth), and only 0.7% of the SNPs had a decent call rate across the diversity panel. This significant reduction of SNP numbers during filtering informed us that the sequence depth is still the limiting factor for NGS application in this polyploid species.

### Population structure in sugarcane

Population structure is an important factor needed to be controlled for GWAS. Several software are available for population structure analysis specifically for diploid species, such as Structure (Pritchard et al. 2000) and Admixture (Alexander et al. 2009), which were not suitable for sugarcane due to its nature of polyploids, outcrossing and clonal propagation, thus not fitting the assumptions in these software. We applied DAPC to capture population structures in this sugarcane diversity panel, which is a multivariate method for the analysis of the population genetic structure not relying on assumptions about Hardy-Weinberg equilibrium or linkage disequilibrium (Jombart et al. 2010). Compared to principal component analysis, DAPC can partition genetic variation into a between-group and a within-group components, and thus achieves clear discrimination of individuals into pre-defined groups. Using 13.8 K high quality and evenly distributed SNPs, we have inferred six groups existed in this diversity panel, including two non*-Saccharum* (*Miscanthus* and *Erianthus*) and four *Saccharum* groups (*S. spontaneum, S. officinarum*, modern *S*. hybrids, and ancient S. hybrids), which aligned perfectly well with biological classification of this diversity panel according to the records of WCSRG. Moreover, DAPC could clearly separate S. *officinarum* from *S*. hybrids, and modern *S*. hybrids from ancient S. hybrids, which are highly genetically similar, supporting the sensitivity of this method to capture miniature population structures in sugarcane.

### GWAS on yield components in a sugarcane diversity panel

We evaluated nine yield components, and performed correlation and path coefficients analyses in a representative sugarcane diversity panel. The path coefficients analyses could suggest appropriate selection strategies to breeding programs by revealing inter-relationship among yield components (Dewey and Lu 1959). For example, we recommended selecting large stalk diameter and leaf width for improving sugar content but for different reasons: a positive direct effect has been estimated for stalk diameter but an indirect effect for leaf width. The net direct effects come from the balance of contributions and competitions among these components. The results of path coefficients analyses informed us that the sizes of stalk diameter might be critical for accumulating high sugar content through improving sugar storage and supporting functions, while only competition for carbon resources existed for leaf width and sugar content (Reflected by negative direct effect). The indirect effect of leaf width on sugar content derived from its influence on stalk diameter. Our results also demonstrated that the same yield component can be favored or against depends on the purpose of breeding programs. For example, genotypes with high stalk number had a high biomass yield but relatively low sugar content. Therefore, comprehensively evaluating and analyzing inter-relationships among yield components in a representative sugarcane diversity panel provided key information for sugarcane breeding programs.

Current studies had three major improvements compared with previous GWAS performed in sugarcane (Débibakas et al. 2014; Gouy et al. 2015; Racedo et al. 2016; Wei et al., 2006, 2010). First, a representative diversity panel with a large number of accessions derived from WCSRG was assembled and used in sugarcane GWAS. This diversity panel consisted of 308 accessions including 279 *Saccharum* accessions and 29 non*-Saccharum* accessions (*Miscanthus* and *Erianthus*). The panel was not limited in sugarcane elite germplasm as previous studies, primarily parental clones and sugarcane hybrids in breeding programs, and had a decent number of accessions from *Saccharum* and non*-Saccharum* species. Therefore, the statistical power of GWAS has been improved in this diversity panel. Moreover, our study has the capability to identify novel genetic resources, which might be not existed in modern sugarcane cultivars due to their narrow genetic basis. More importantly, these novel robust traits could be quickly introgressed into sugarcane breeding programs through controlled crosses, especially with help of associated DNA markers. Second, we applied TES, an NGS-enabled genome complexity reduction sequencing technology, to genotype this diversity panel, which was proven to be very efficient for investigation of sequence variants in sugarcane (Song et al. 2016). With a stringent sample-by-sample and population-level filtering, 74.9 K markers were retained for GWAS, which was a much larger number of markers than any of other GWAS in sugarcane. Also, TES achieved an average sequencing depth of 99x in this study, which allowed us to differentiate all dosages, even for dodeca-ploid species with up to 13 possible genotypes (Yang et al. 2017). The accurate genotypes were the prerequisite to perform GWAS with allele dosages under consideration in this polyploidy species, which were usually simplified as diploids in previous studies. Third, thanks to the development of GWASpoly (Rosyara et al. 2016), GWAS was performed in sugarcane with six models regarding different gene actions. Depending on the traits and loci analyzed, different gene actions may exist. Without further experimental support, we couldn’t evaluate which models are more suitable. To be inclusive, all the six models were tested to maximize the discovery of associated sequence variants.

We identified 217 non-redundant markers and 225 genes associated with the nine yield components. We compared our results with GWAS or linkage mappings in sugarcane. Unfortunately, comparisons were relatively limited since only researches on the same traits and with the associated markers or genes aligned to the sorghum genome can be compared, and only a few such studies were published previously. For the sugar content (Brix), two (*2P5849859* and *2P65250224*) out of three associated markers were found with a distance of 333.1 Kbp and 1,631.3 Kbp, respectively from the associated gene (*Sb02g004780* and *Sb02g028450*) identified previously (Racedo et al. 2016). Recently, quantitative trait loci (QTLs) mapping has been conducted in a bi-parental population of 173 progeny with multiple QTLs detected for Brix, water content and stalk diameter (Islam et al. submitted). We had associated markers with a distance of 1,615.6 Kbp, 6,556.7 Kbp, 3,165.5 Kbp and 591.1 Kbp from their closest markers (3SNP463 for stalk diameter, 2SNP3040 for Brix, 1SNPUN315 and 3SNP3118 for water content), respectively. However, there were several QTLs from this QTL mapping not detected in our study. Multiple reasons could led to this discrepancy, including plant materials (bi-parental population vs. diversity panel), statistical methods (linkage mapping vs. GWAS), and stringent threshold for significance declared (Logarithm of odd < 3 vs. *P* < 0.05 after Bonferroni correction) etc. We also compared our results with selection genes identified in another study (Yang et al 2018). Eight of our associated genes were also under selection in sugarcane breeding programs. Interestingly, the candidate gene (*Sobic.009G000100*), also under selection and encoding SIN3 histone deacetylase protein, contained four sequence variants (one InDel in intron, and three SNPs in coding region causing one synonymous mutation and two non-synonymous mutations), which were identified for dry weight at least in two GWAS runs or models with -log (*P*) ranging from 5.71 to 8.15. The combination of different haplotypes and dosages of this gene might explain the segregation of dry weight in the diversity panel. The comparisons corroborated the reliability of the results. These markers and genes associated with yield components should be fully utilized in sugarcane breeding programs.

### ROHs in sugarcane

For the first time, we assessed ROHs in this outcrossing sugarcane using high-density markers. The results of ROHs revealed several interesting phenomena in this polyploidy species. First, the investigation of ROHs could reflect population migration, mating history and structure of sugarcane. *Non-Saccharum* 2 had high NROHs and SROHs but the same AROHs as other sub-populations. SROHs were correlated with NROHs in *non-Saccharum* 2 (Correlation coefficient 0.99), suggesting increase of the number of ROHs leading to a large SROH in this species. A sudden reduction of population size might happen in non*-Saccharum* 2 and crosses in a population with limited number of individuals could result in a drastic increase of NROHs. In addition, ROHs were not detected in ancient *S*. hybrids and only two ROHs in modern *S*. hybrids. Ancient S. hybrids and modern S. hybrids were derived from the hybridization between the wild species *S. spontaneum* and the cultivated *S. officinarum* (D'Hont et al. 2002). Compared with their ancestors, the ROH results showed the inter-species crosses broadened the genetic diversity and then reduce ROHs in sugarcane hybrids. The two ROHs in modern S. hybrids (One for each accession) were located at Chr07:15.9 Mbp according to the sorghum genome, and were the common ROHs detected in 12 accessions in this diversity panel. The results supported the increase of genomic inbreeding in modern sugarcane cultivars due to multiple backcrosses to *S. officinarum*, and probably their narrow genetic basis. Second, the occurrence of ROHs were likely under selection because the majority of ROHs were common (96.4% were detected in at least two accessions). More interestingly, the genes landing in ROHs were enriched for certain categories with specific gene functions. In our case, the enriched genes in the most common ROHs were involved in plant photosynthesis, suggesting that increase of genomic inbreeding of the regions harboring genes related with photosynthesis would have negative impacts on sugarcane yield. Third, we observed that ROHs were significantly associated with total weight and dry weight, and increase of ROHs had negative influence on these traits. The results further emphasize the importance to broad genetic basis in order to maintain sugarcane production specifically the biomass.

## Author contribution statement

JW conceived the experiment. XY performed the research and analyzed the data. JT and SS measured the yield components. ZL helped prepare marker for GWAS. XY and JW wrote the manuscript. All authors reviewed and approved this submission.

## Acknowledgments

This research is financially supported by Florida Sugar Cane League and the Office of Science, U.S. Department of Energy (Award number: DE-SC0006995).

### Compliance with ethical standards

#### Conflict of interest

The authors declare that they have no conflict of interest.

## Online supporting information

**Figure S1.**
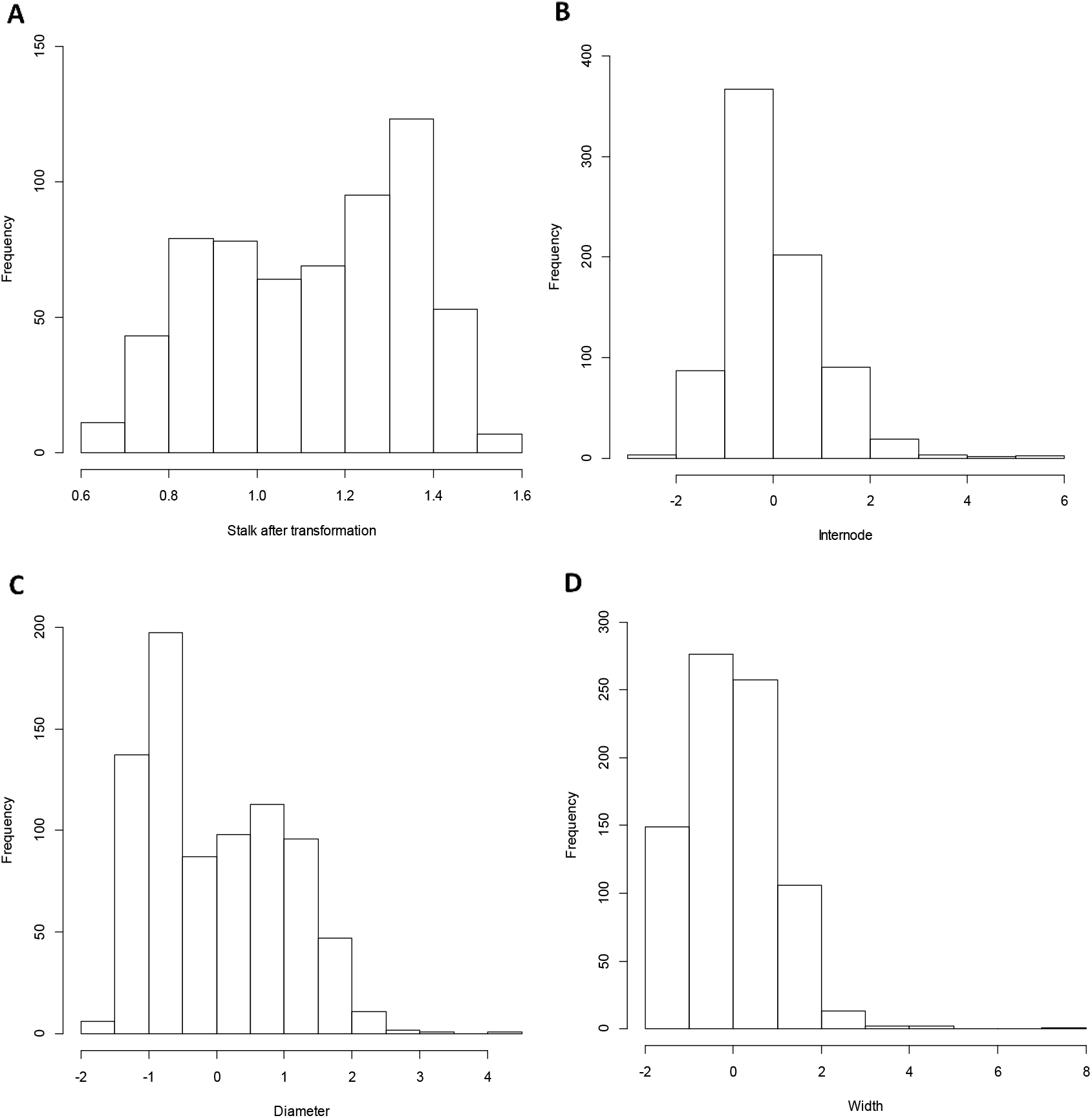
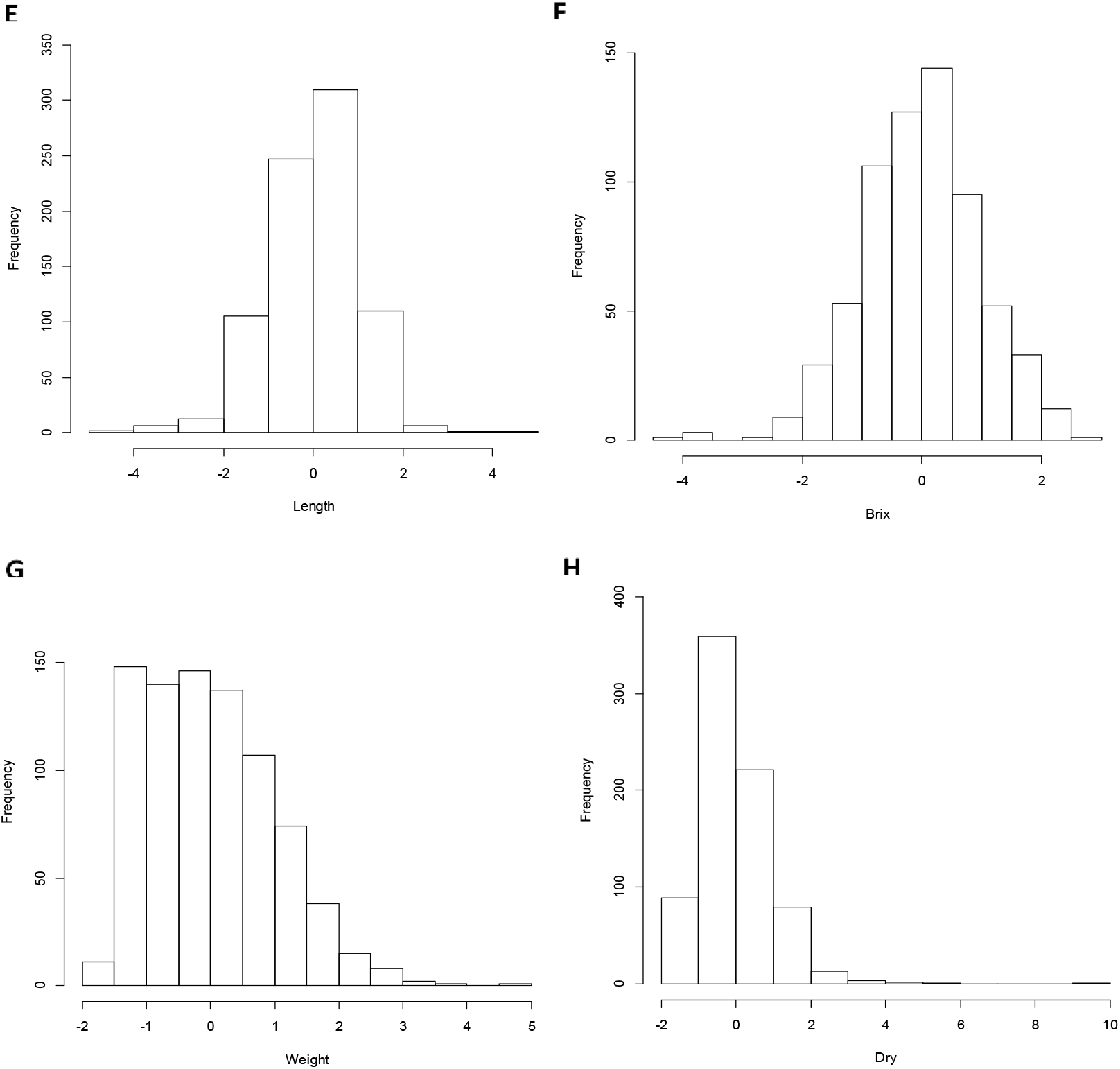
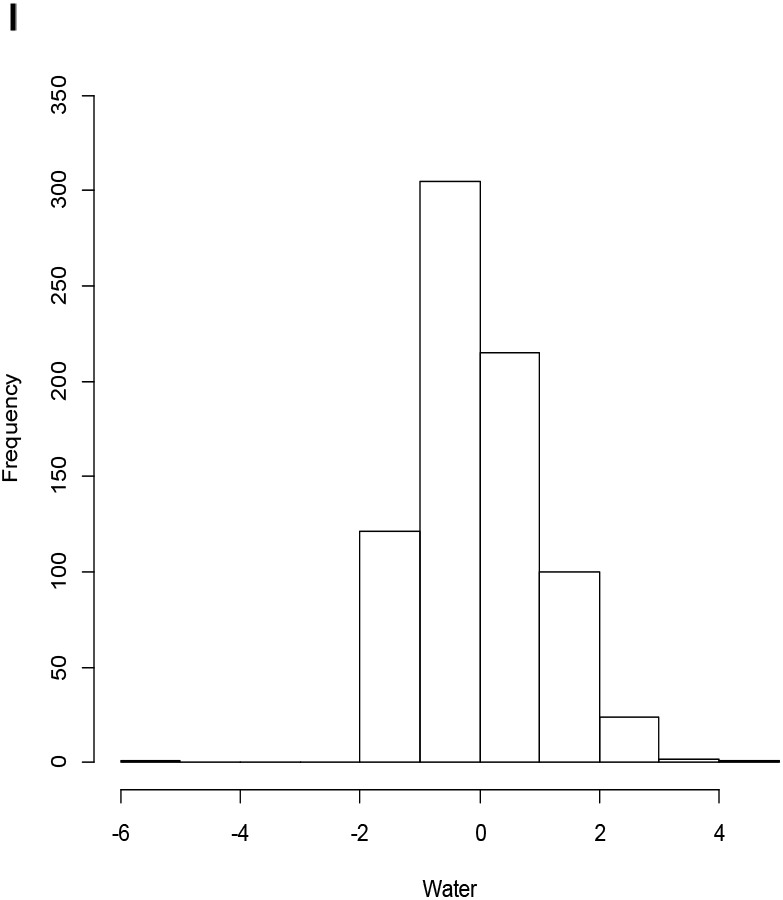
Distribution of the nine yield components in the sugarcane diversity panel, (**A**) Stalk number (Stalk); (**B**) Internode length (cm) (Internode); (**C**) Stalk diameter (mm) (Diameter); (**D**) Leaf width (mm) (Width); (**E**) Leaf length(cm) (length); (**F**) Brix; (**G**) Total weight(kg) (Weight); (**H**) Dry weight(kg) (Dry); (**I**) Water content(kg) (Water). For each trait, Z-score for each accession was calculated in each replicate, and Z acores of the diversity panel in three replicates were plotted.

**Figure S2.**
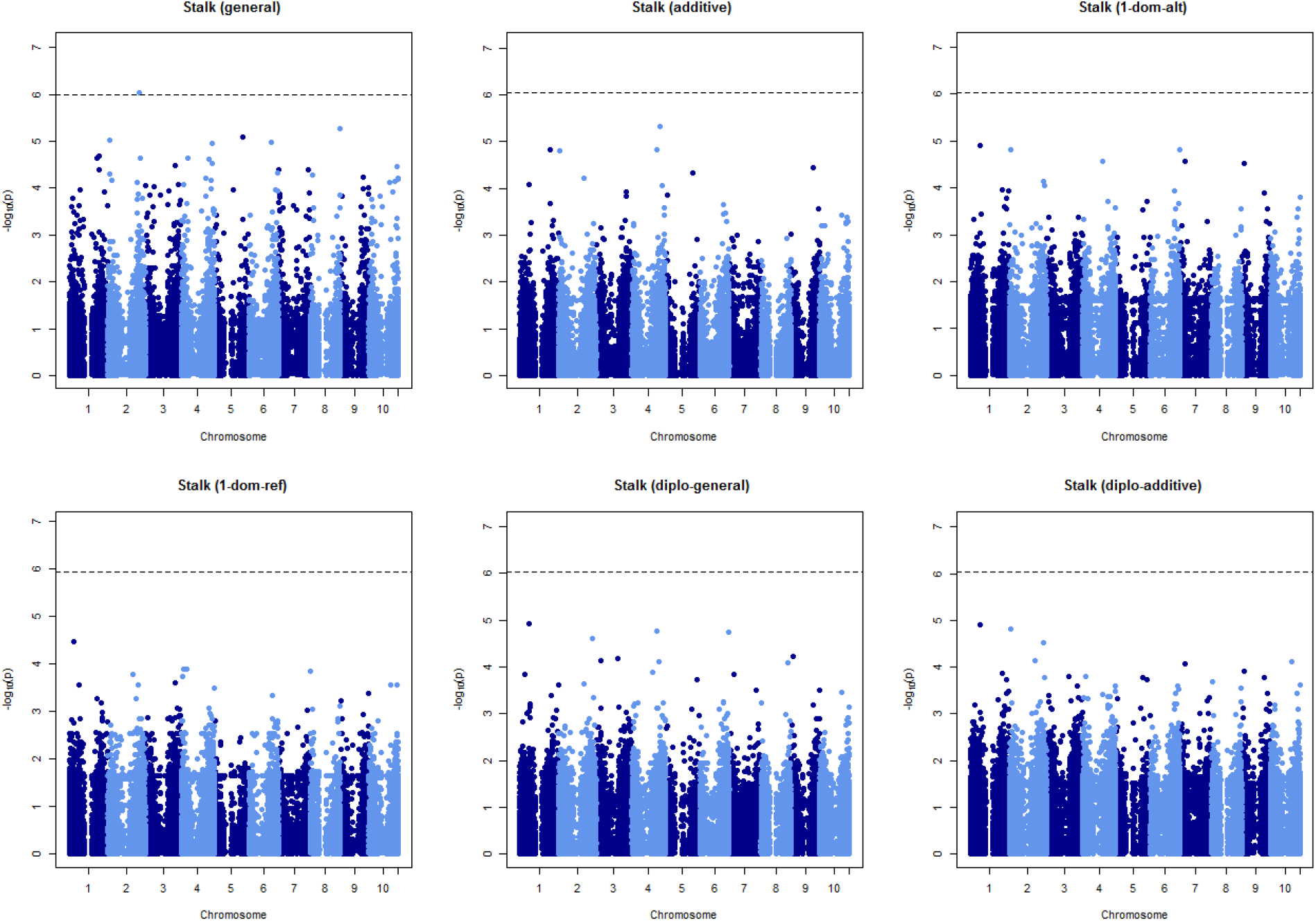
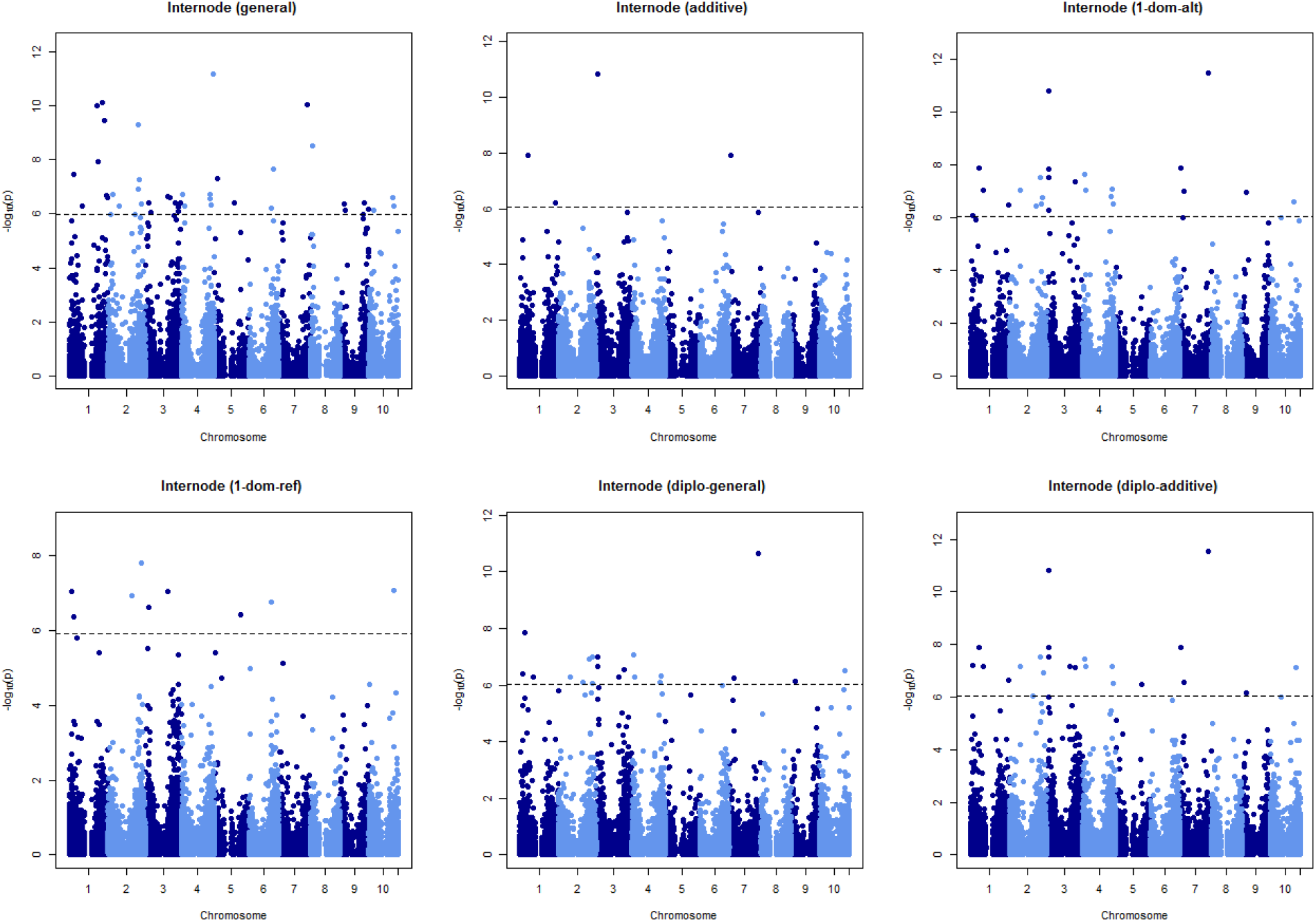
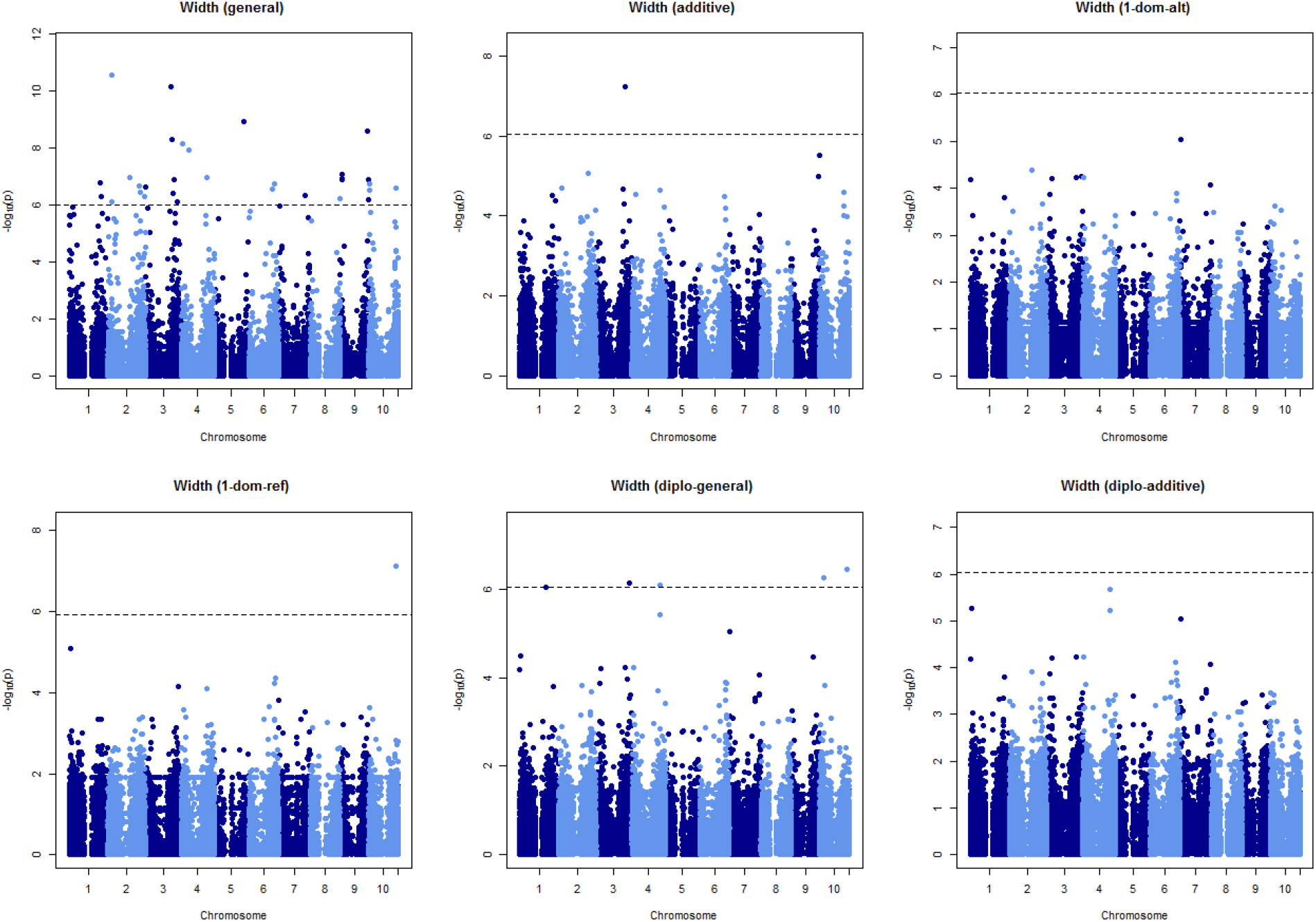
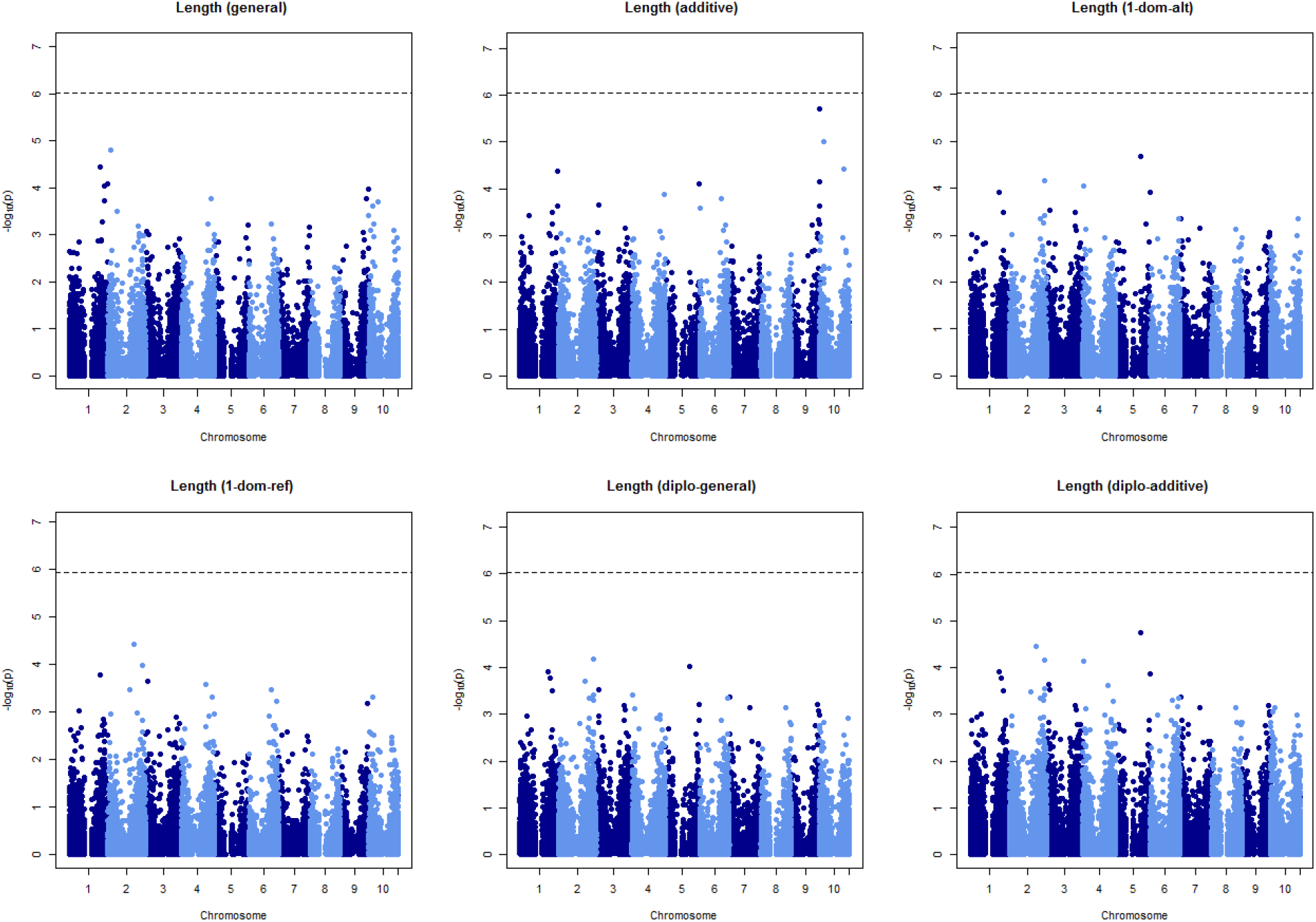
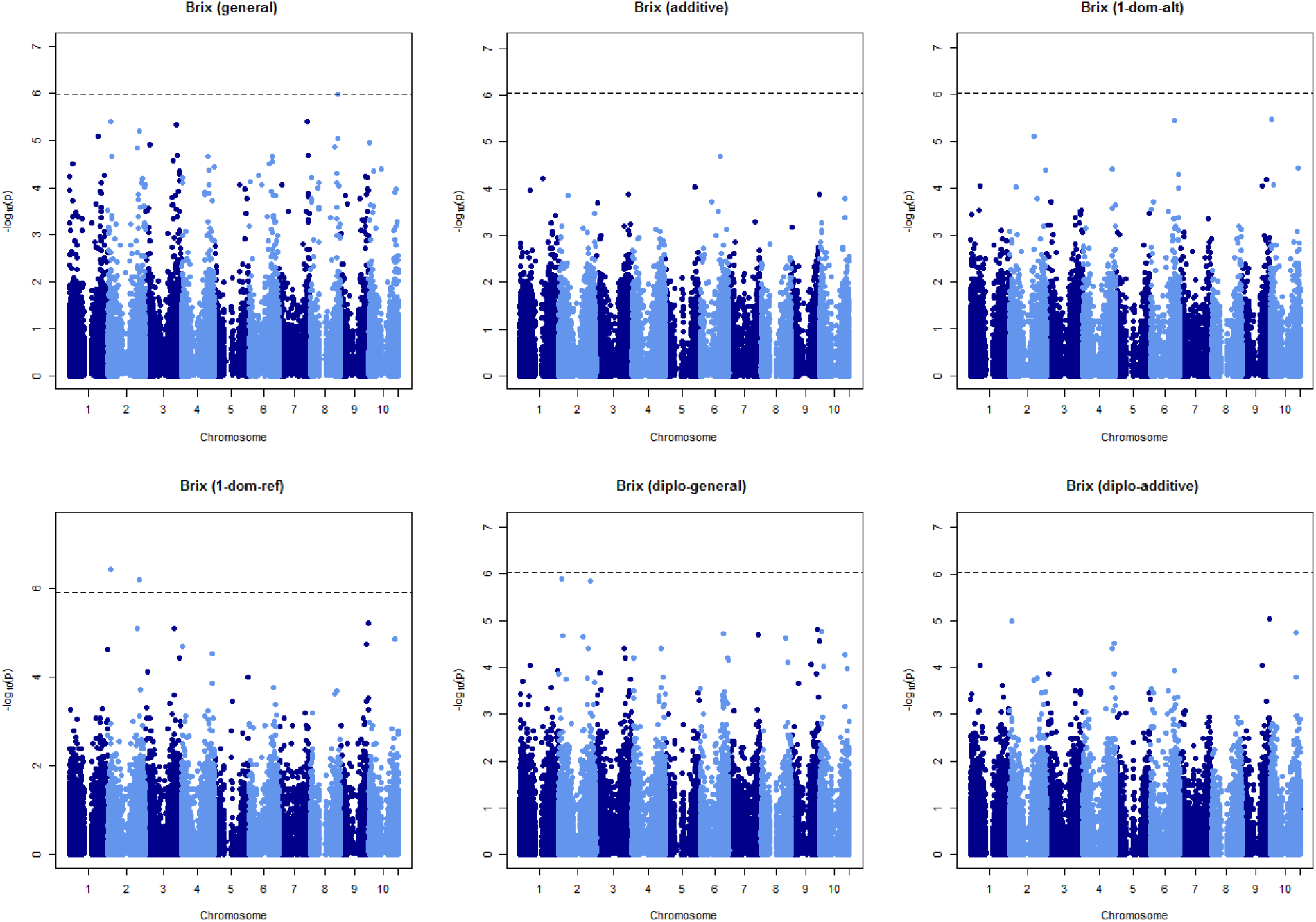
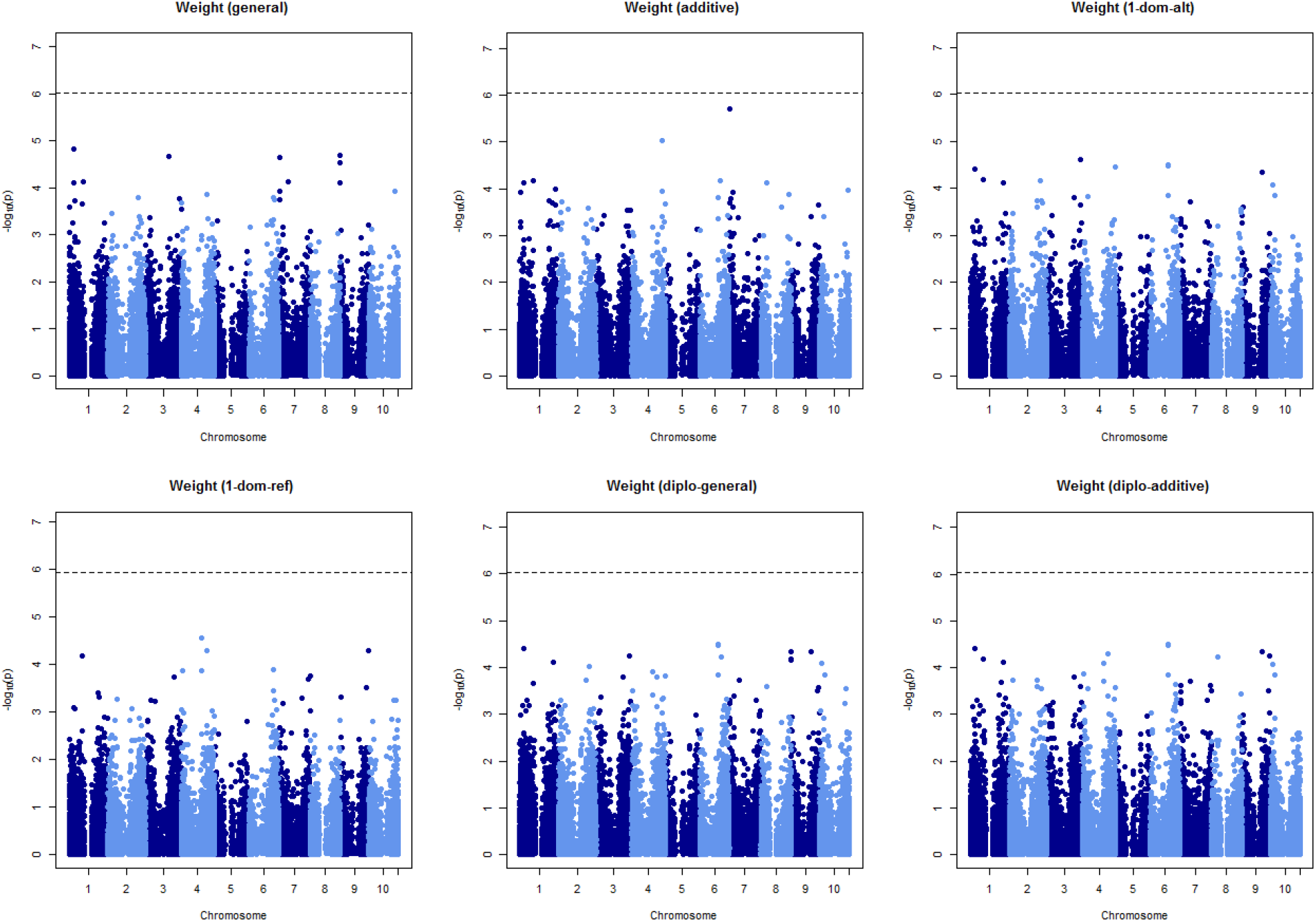
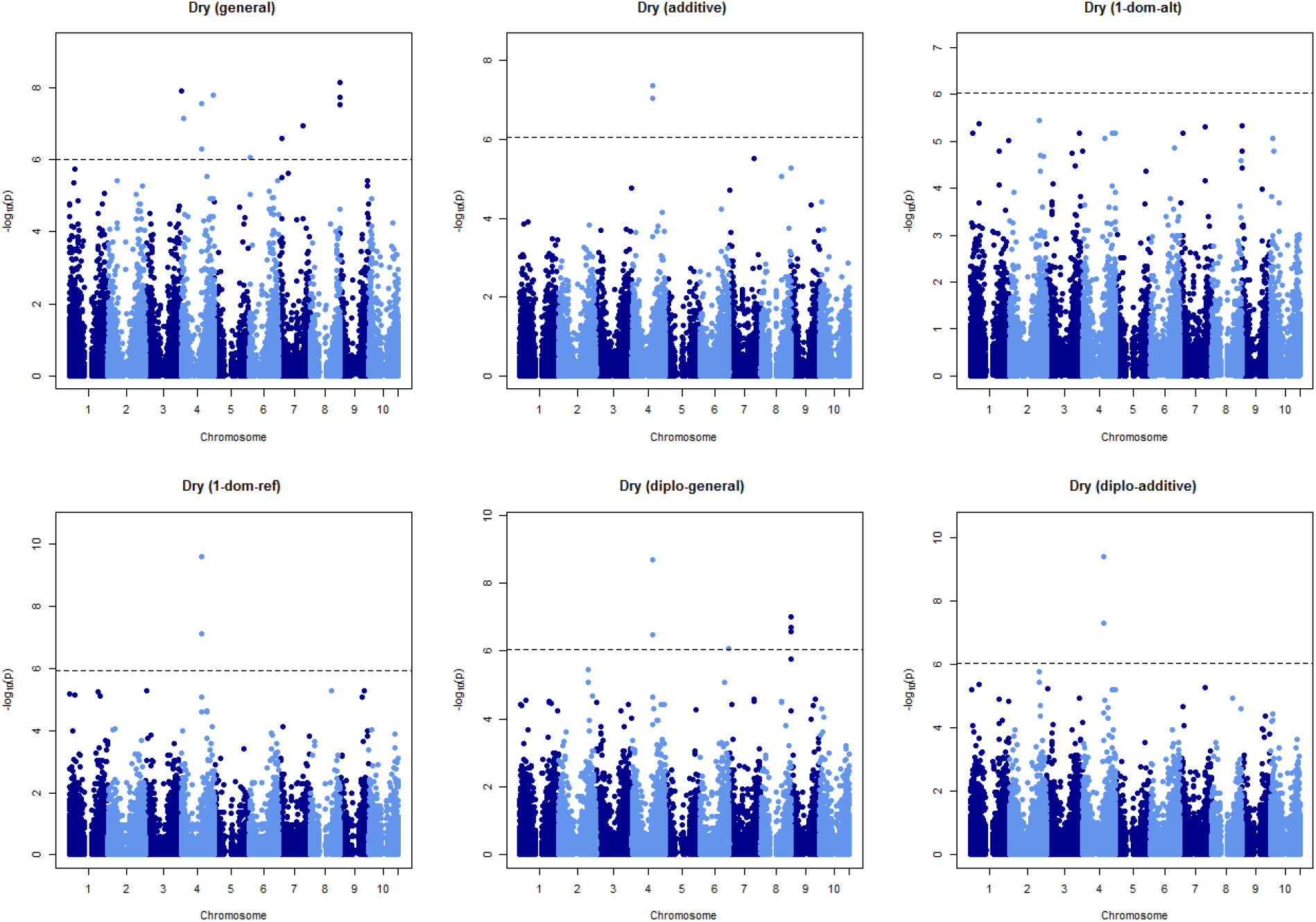
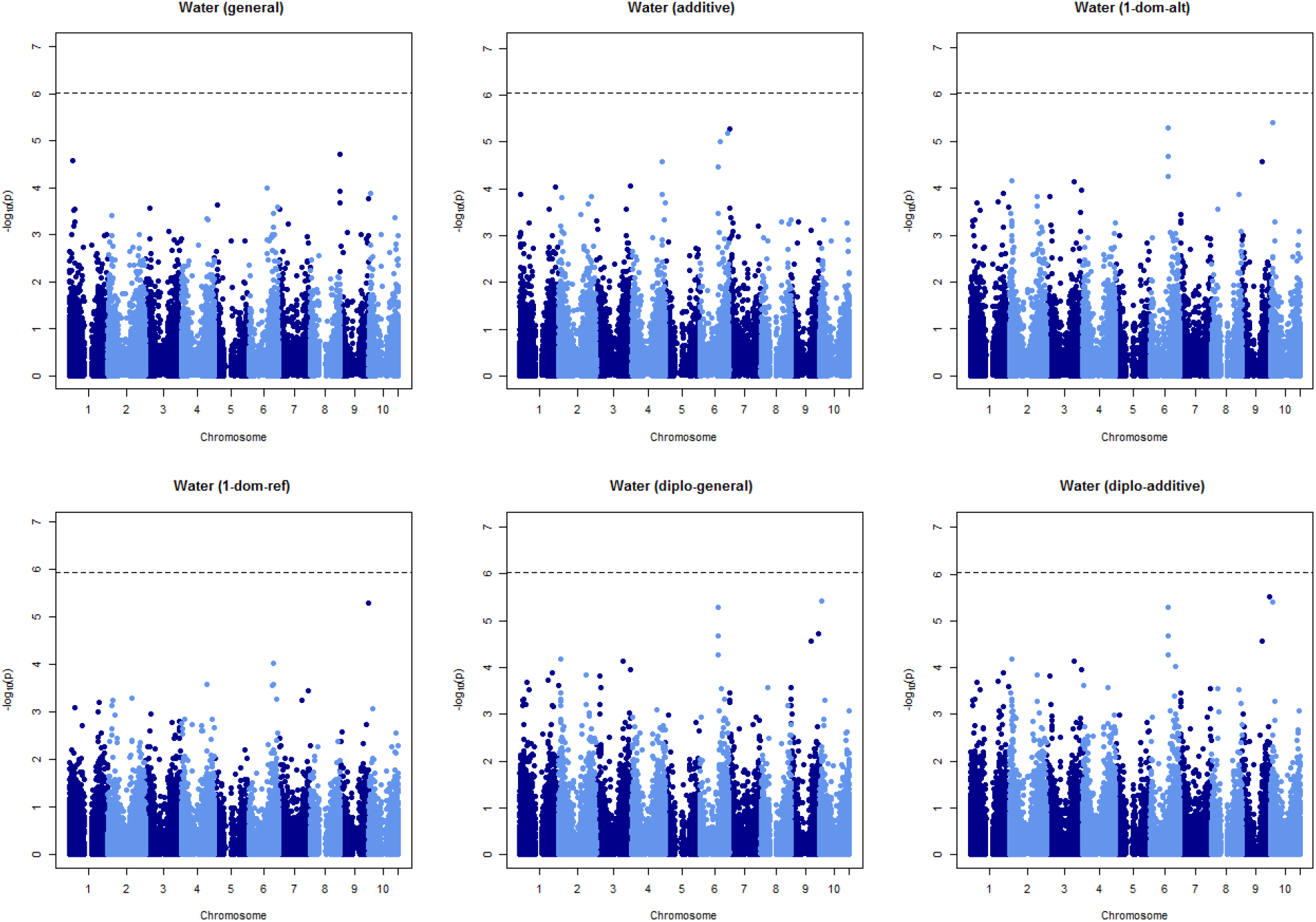
Significant marker-trait associations identified in full set of the sugarcane diversity panel with 300 accessions. Stalk = Stalk number; Diameter = Stalk diameter; Width = Leaf width; Length = Leaf length; Weight = Total weight; Dry = Dry weight; Water = Water content; Internode = Internode length.

**Figure S3.**
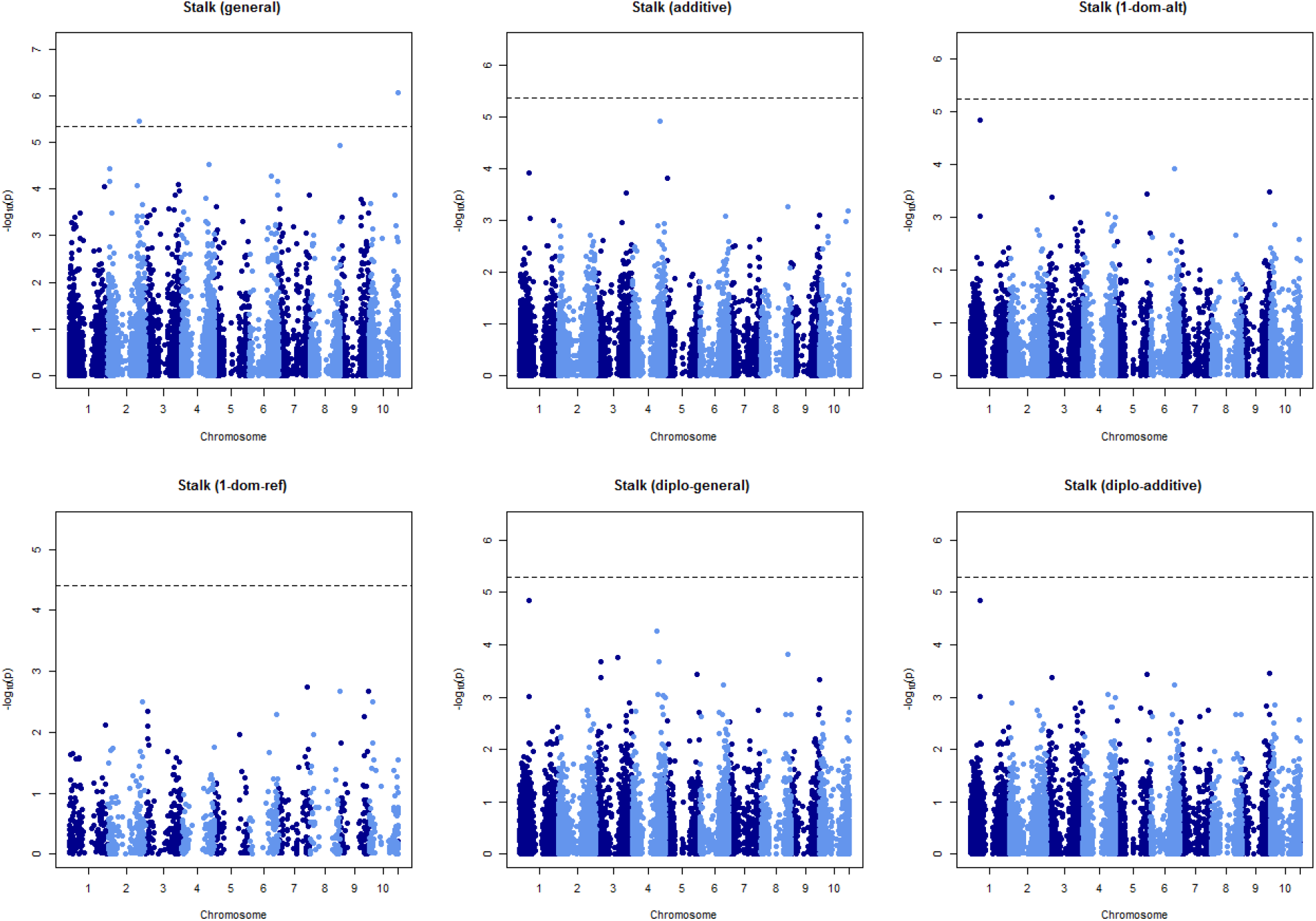
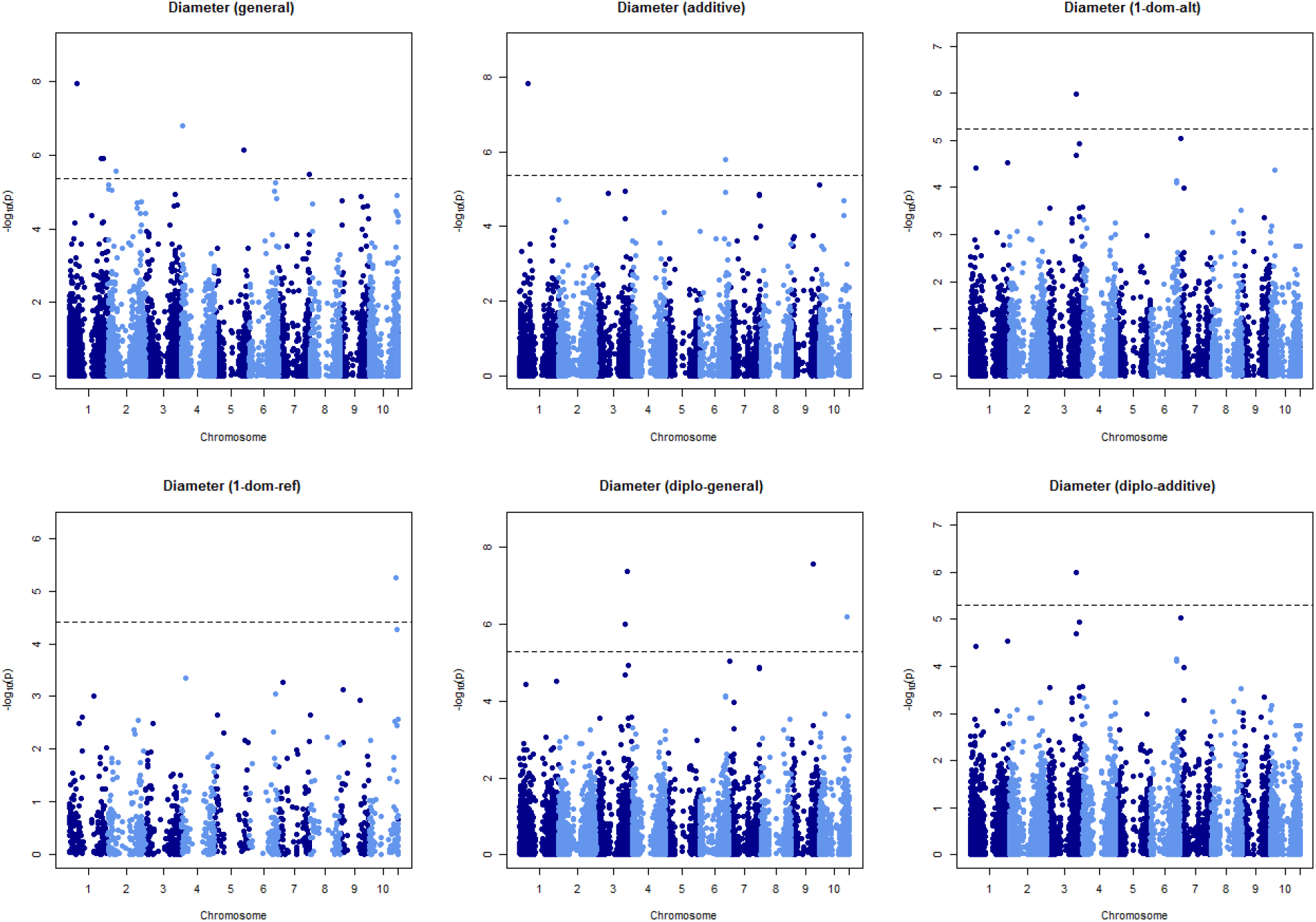
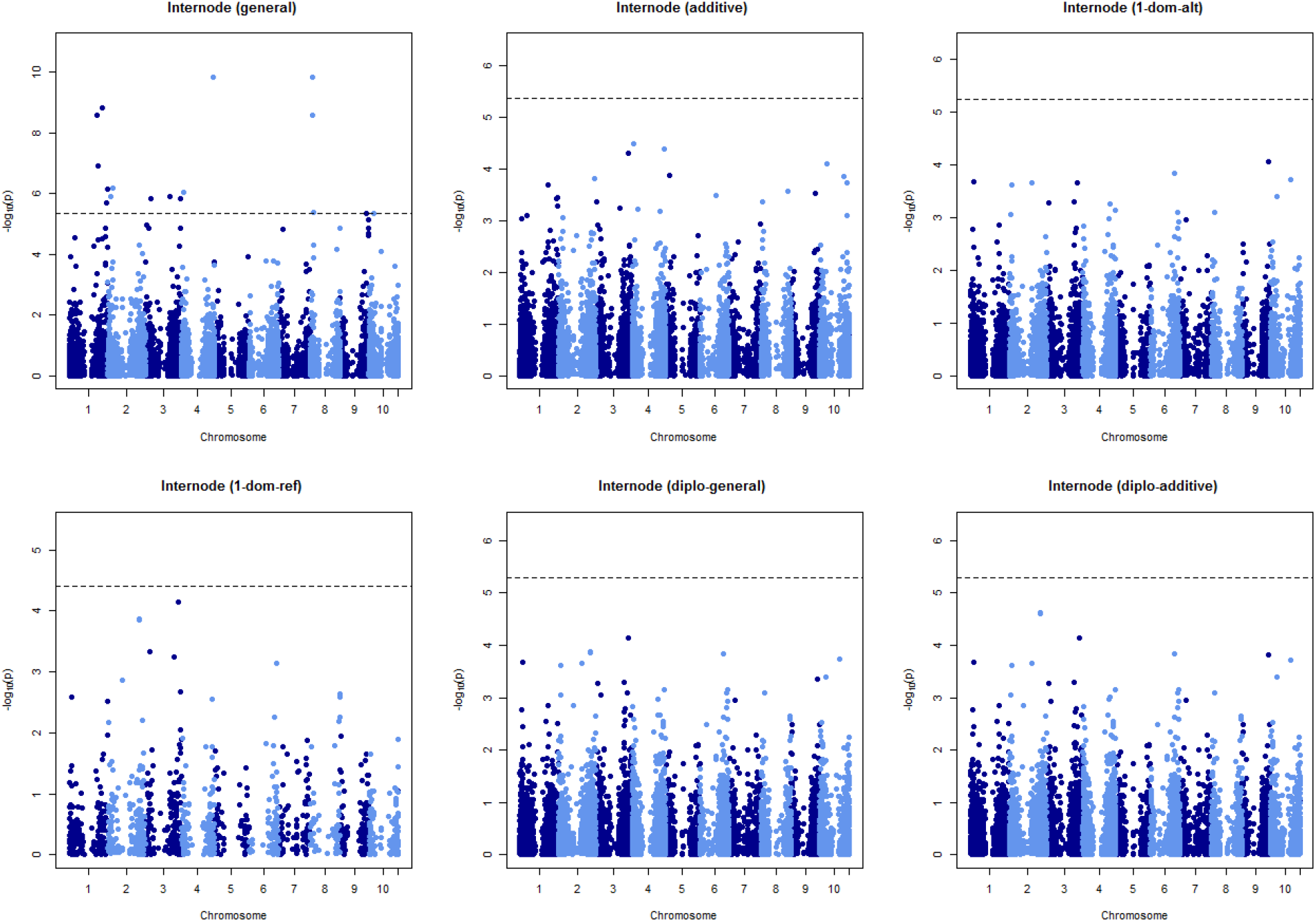
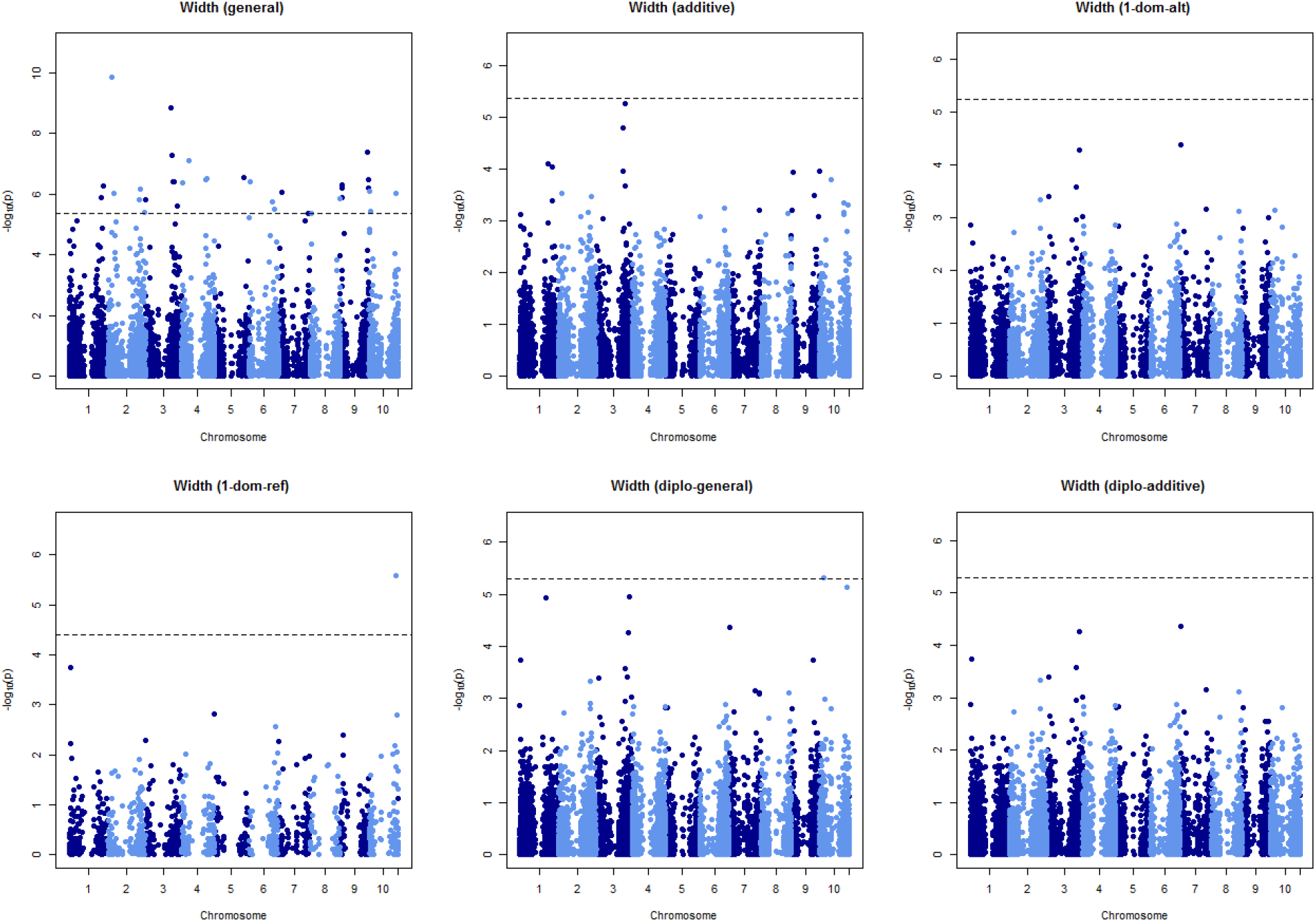
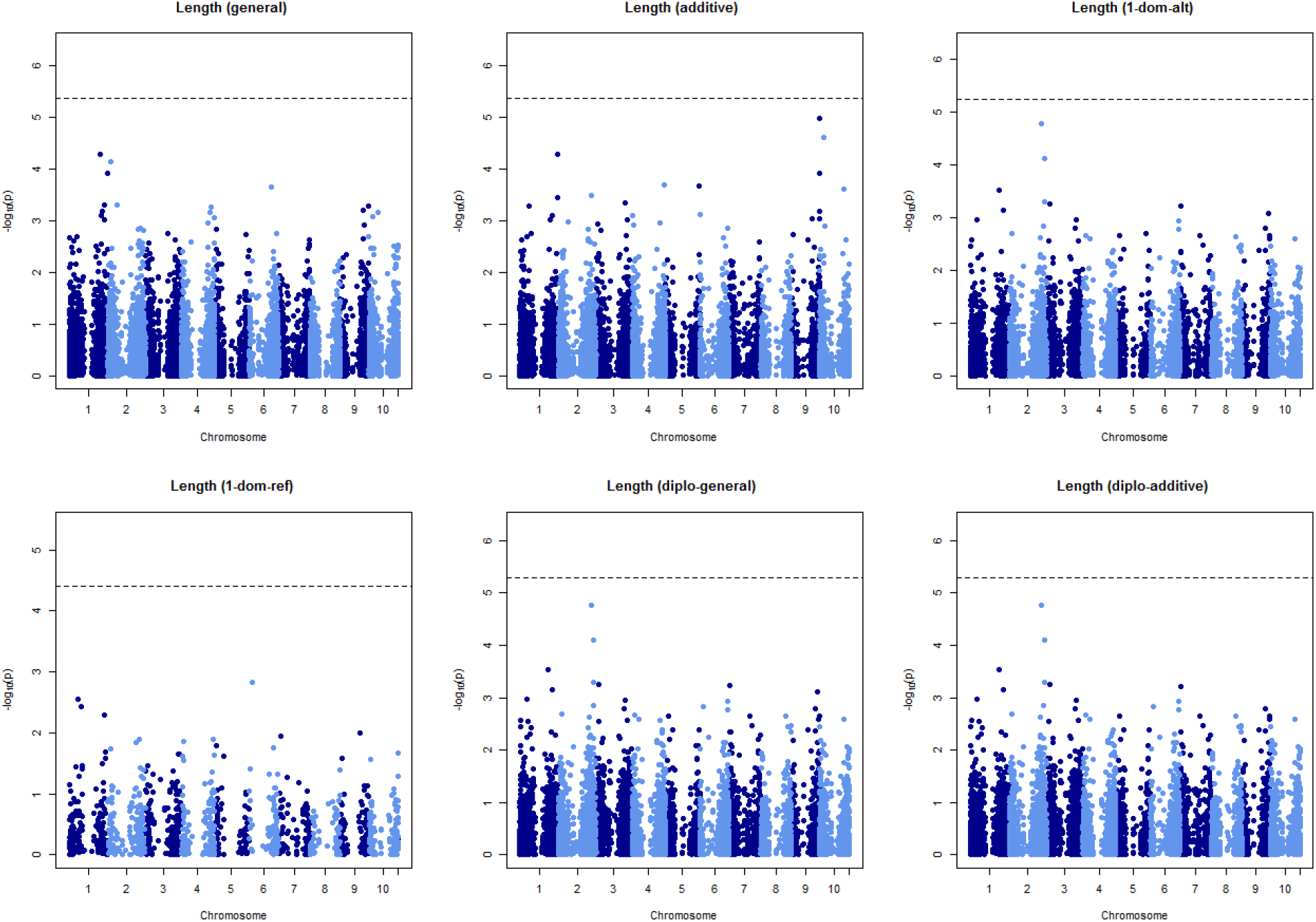
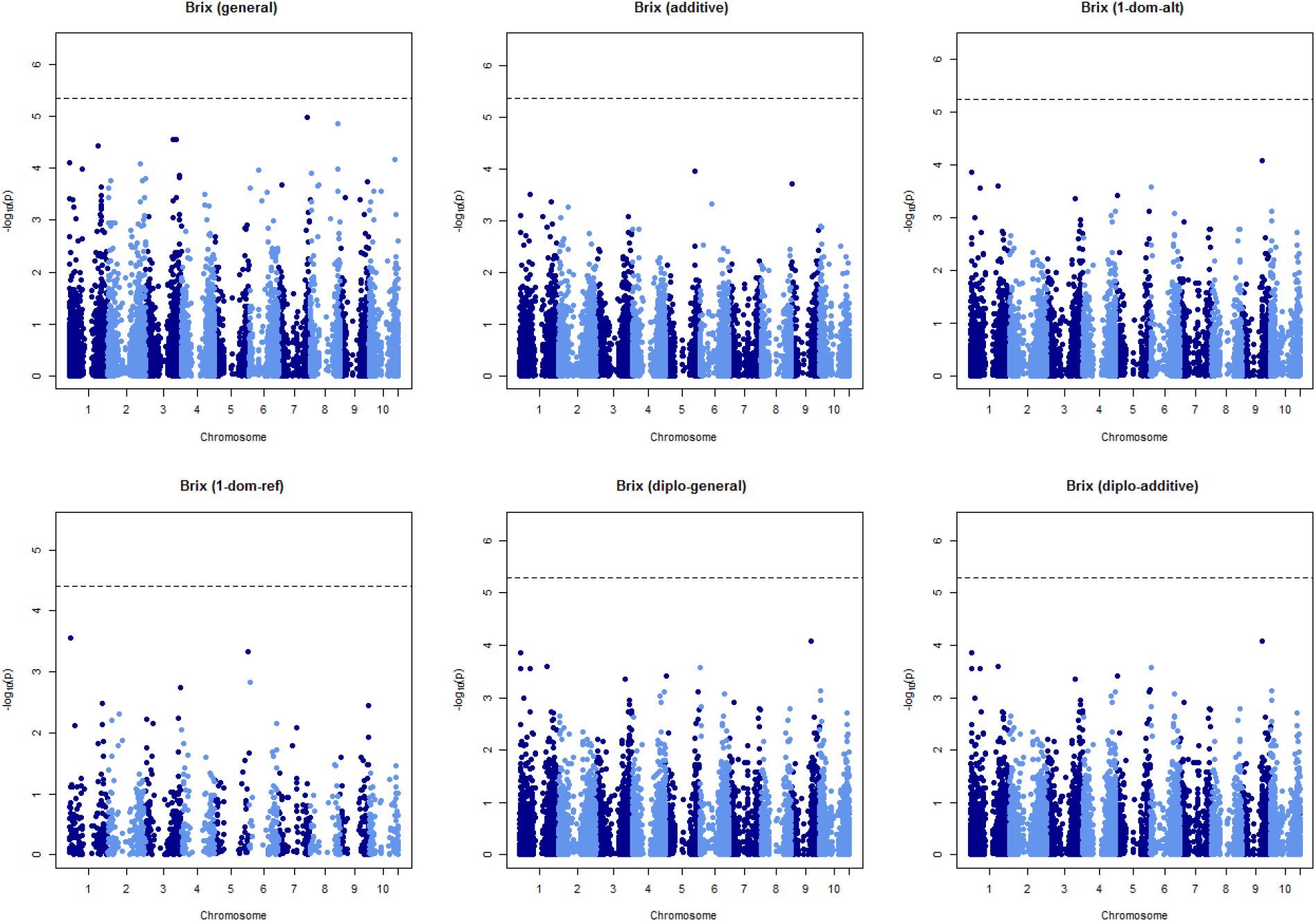
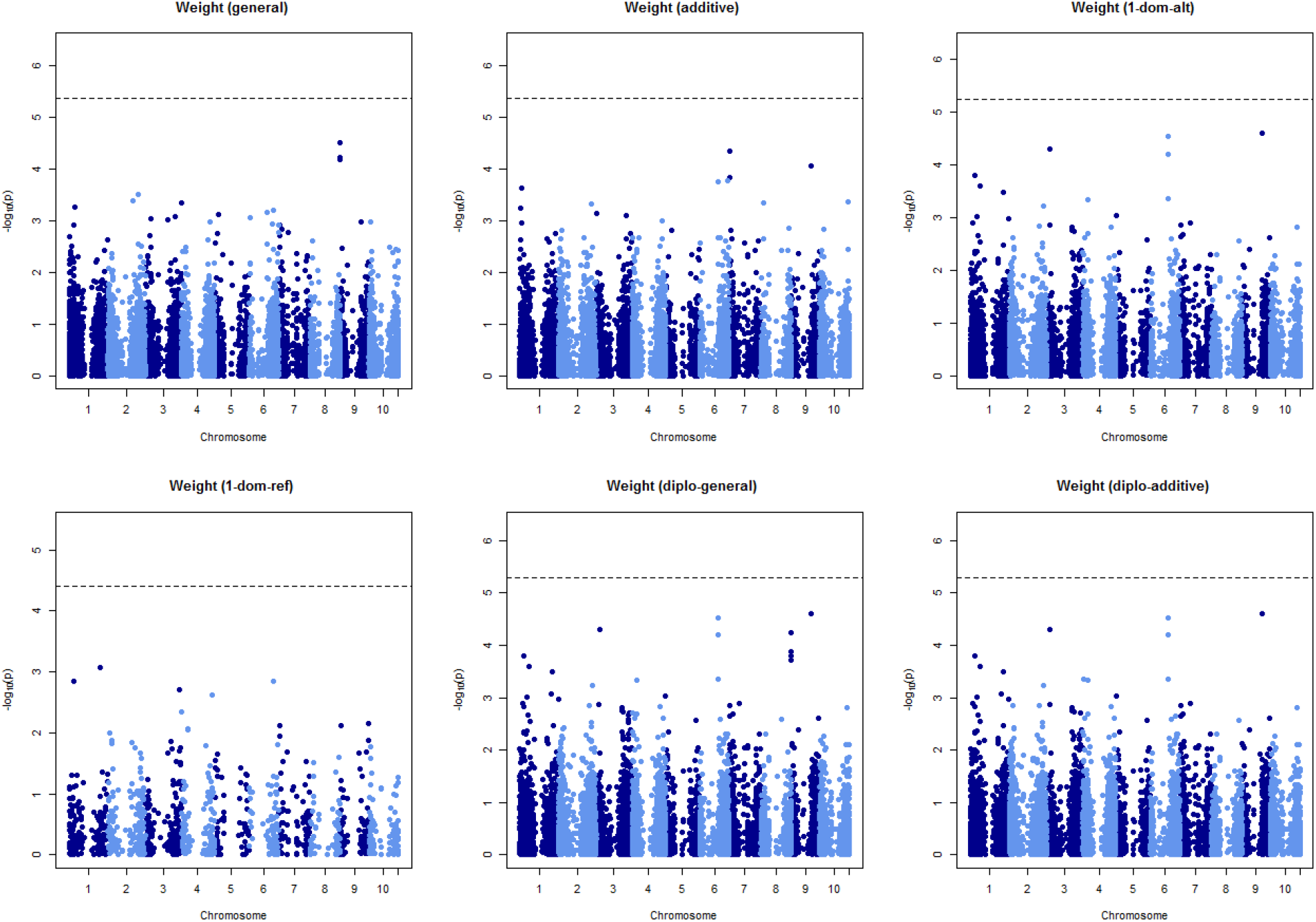
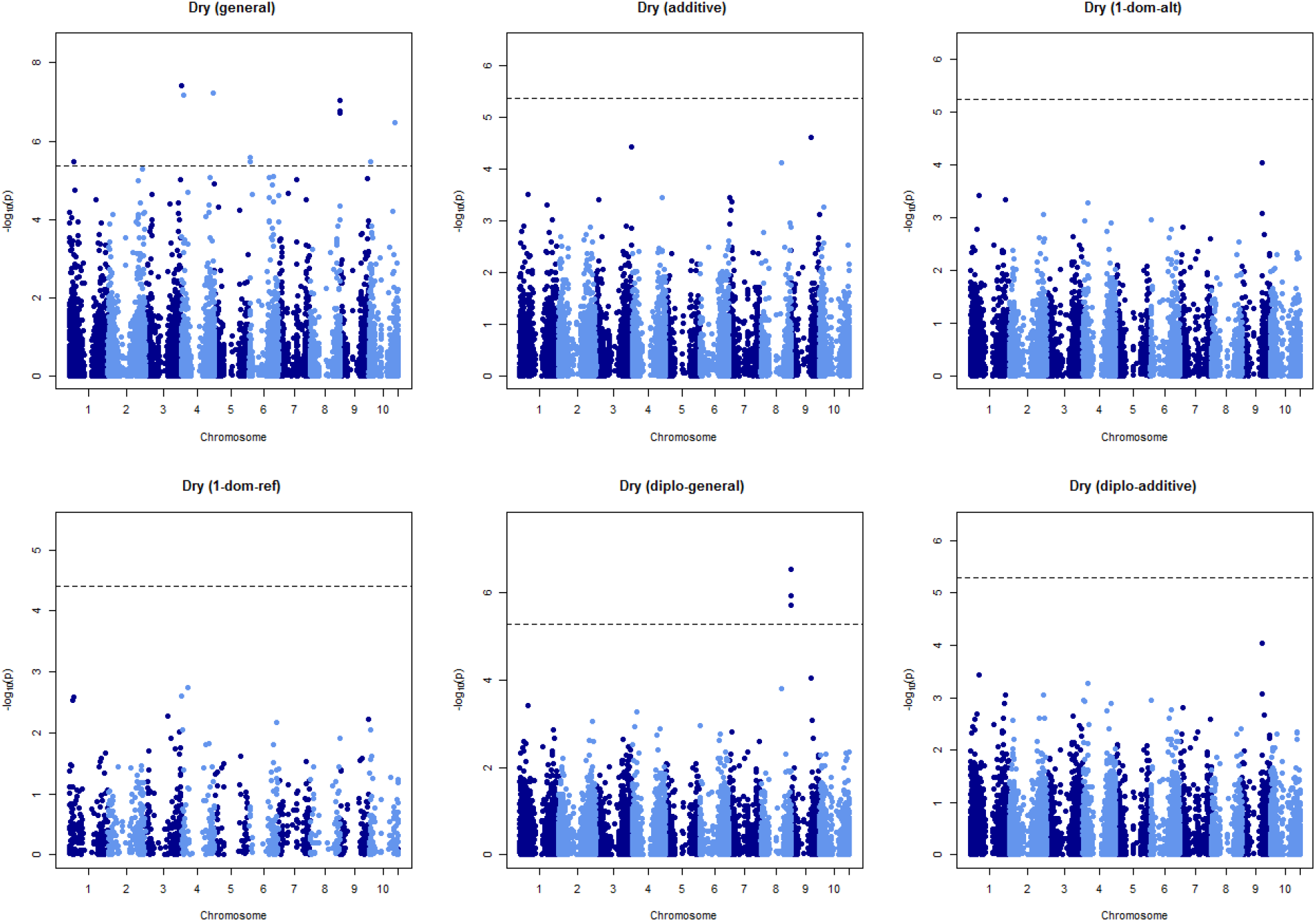
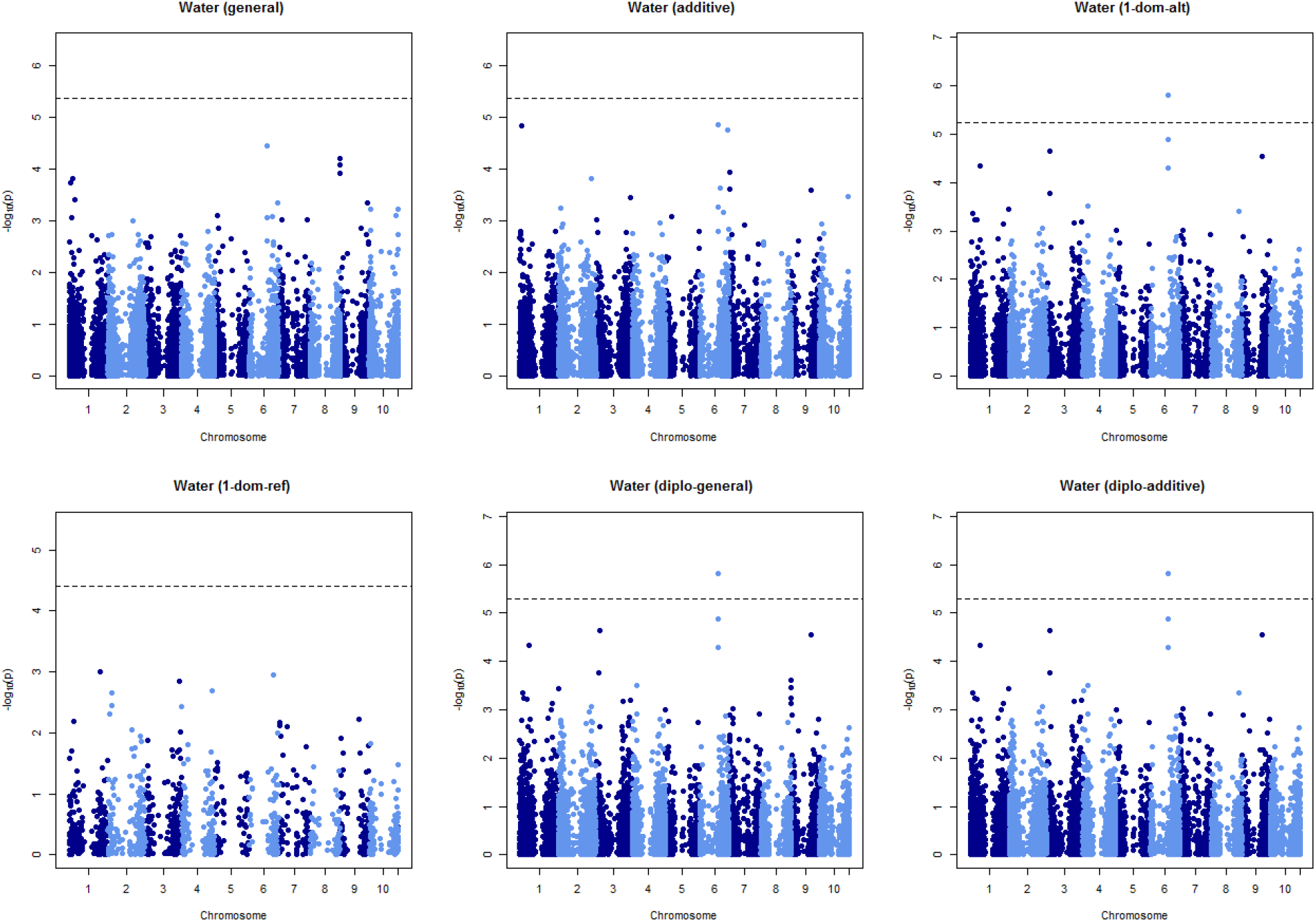
Significant marker-trait associations identified in *Saccharum* set of the sugarcane diversity panel with 273 accessions. Stalk = Stalk number; Diameter = Stalk diameter; Width = Leaf width; Length = Leaf length; Weight = Total weight; Dry = Dry weight; Water = Water content; Internode = Internode length.

**Figure S4.**
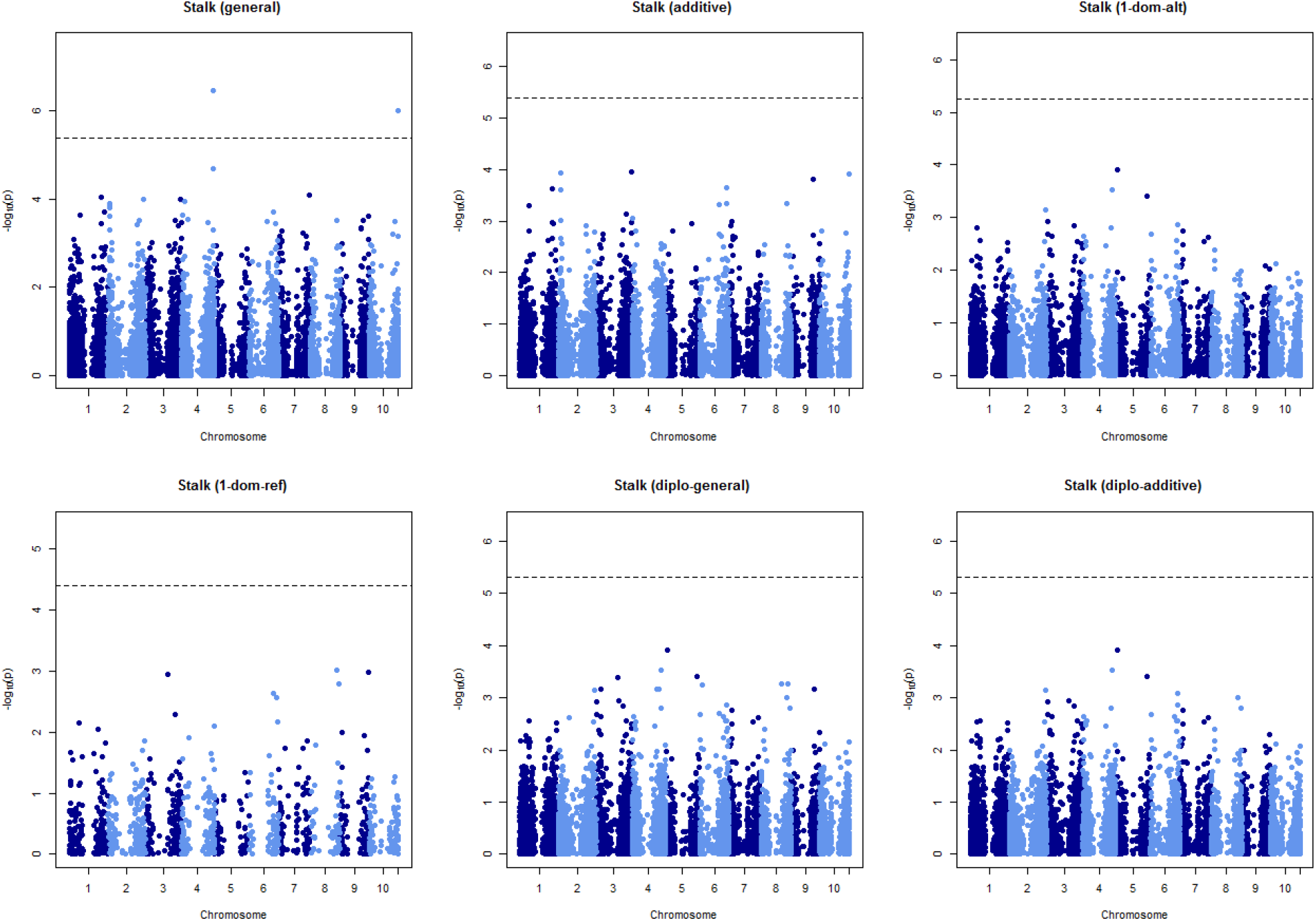
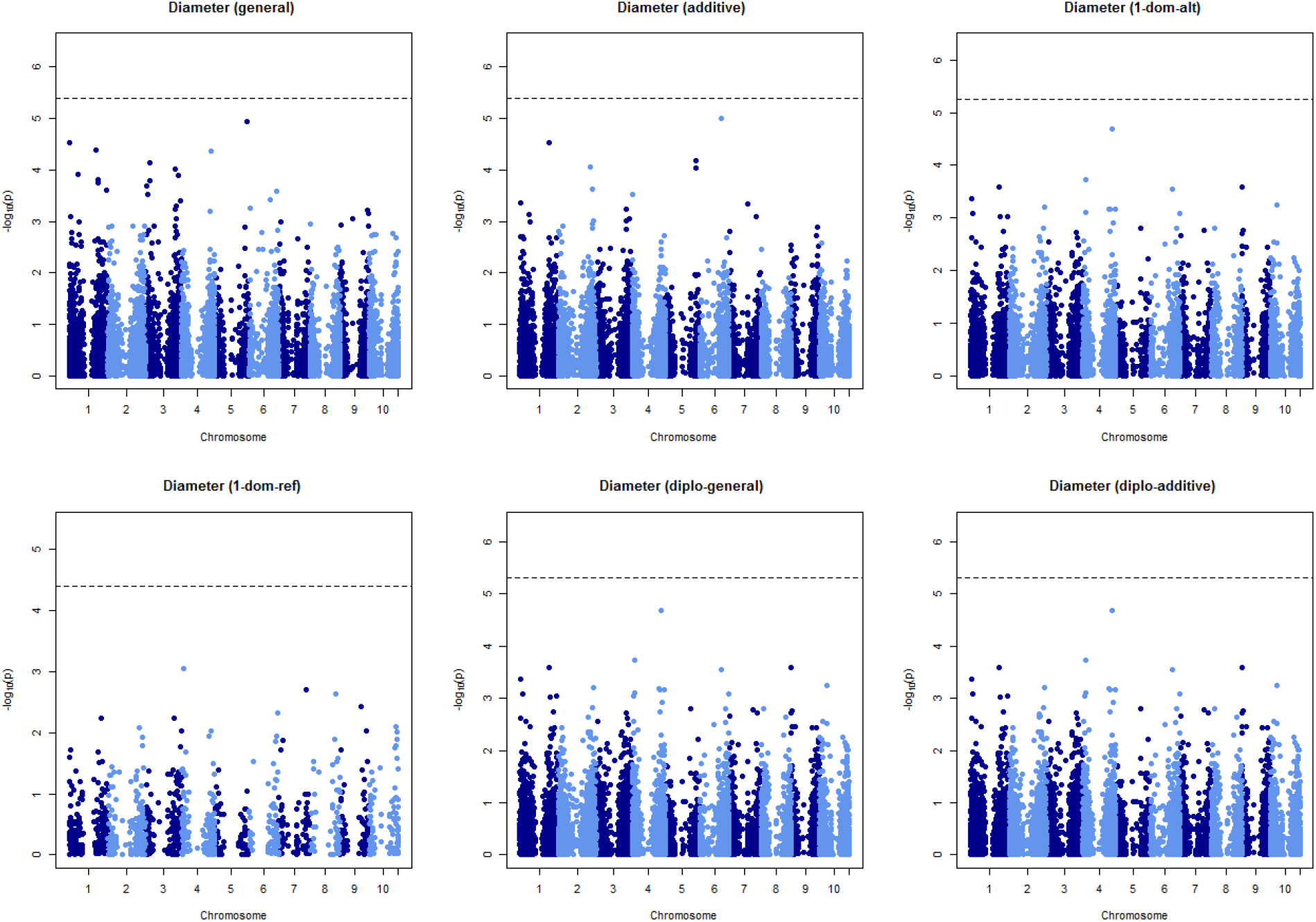
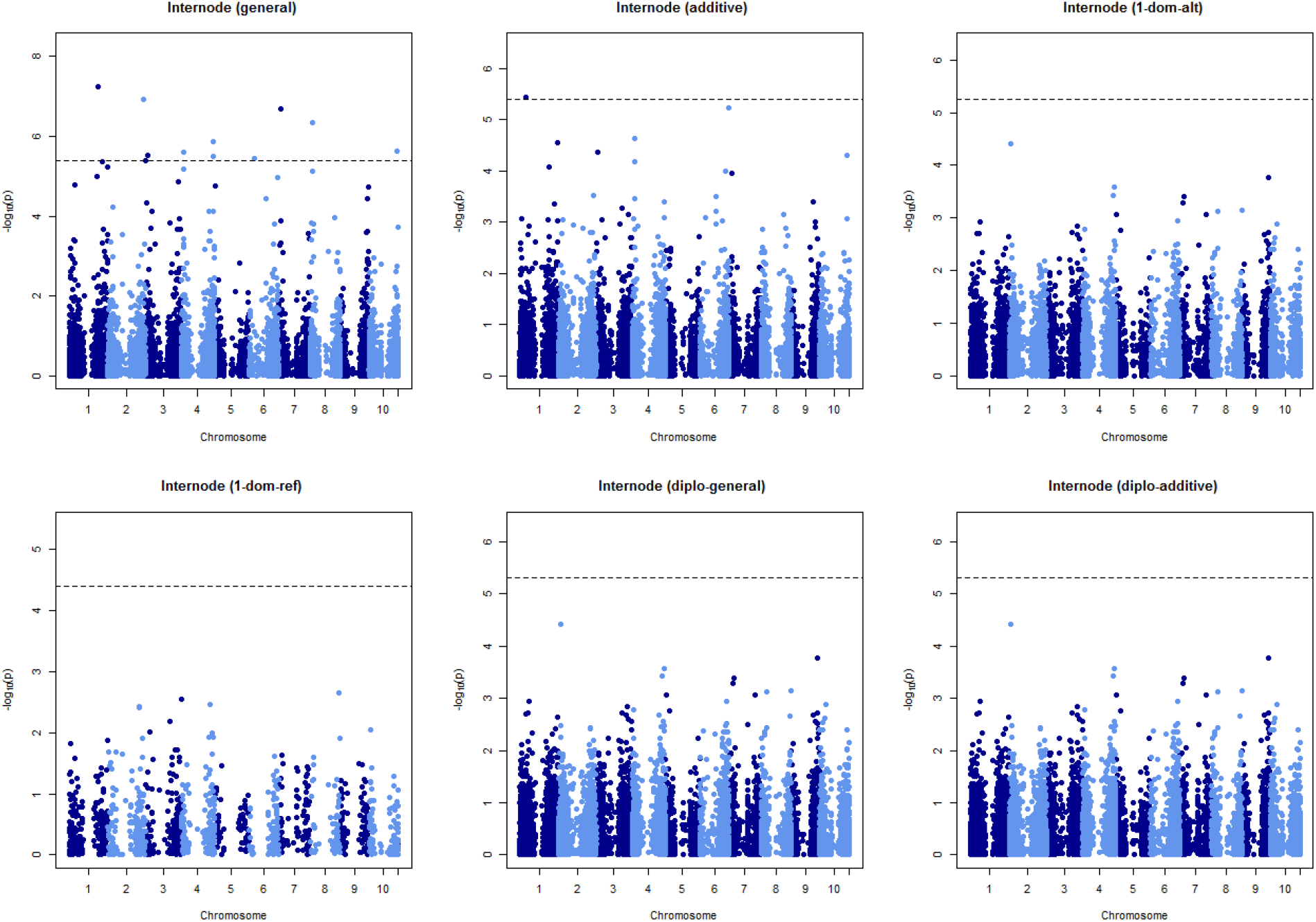
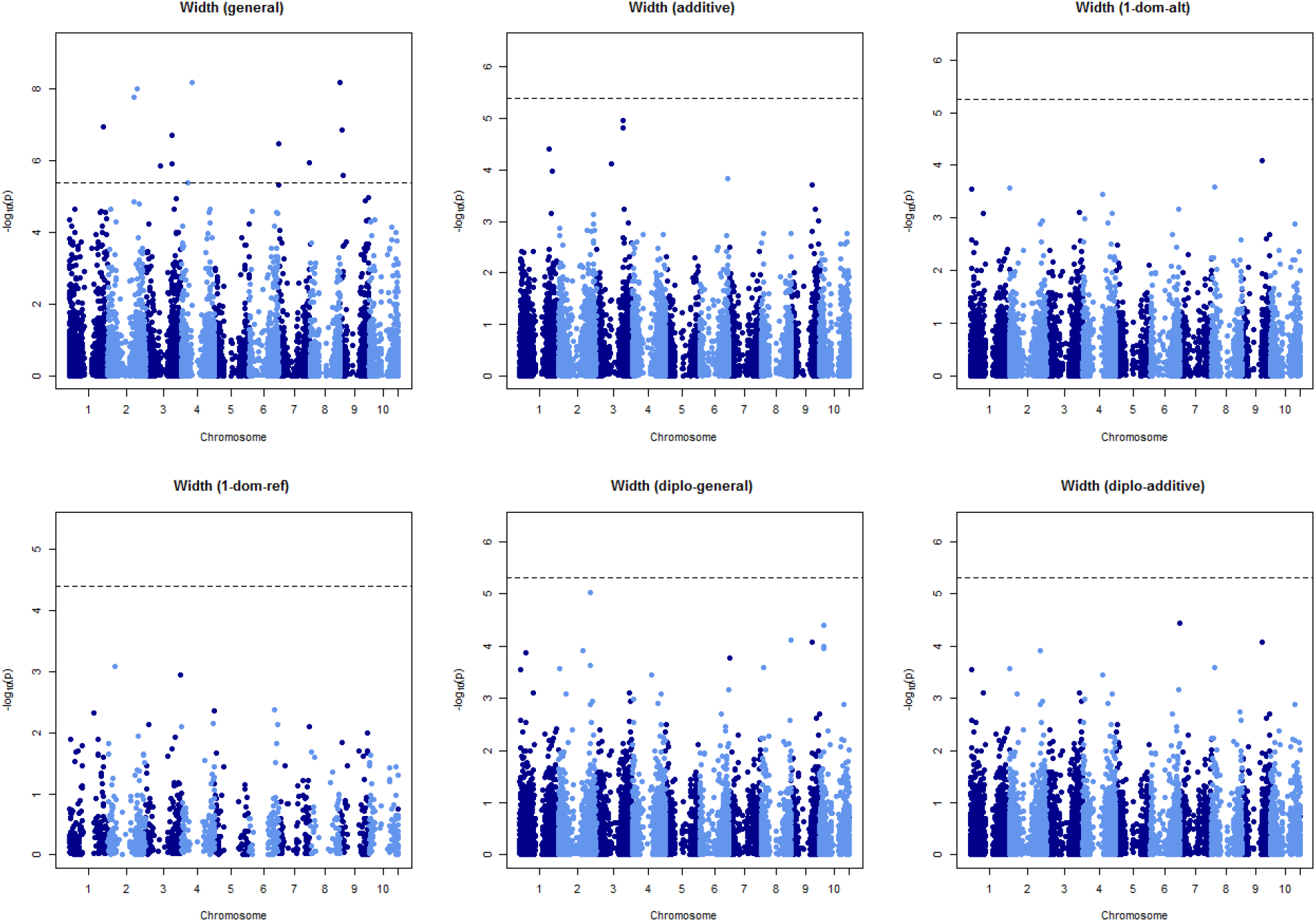
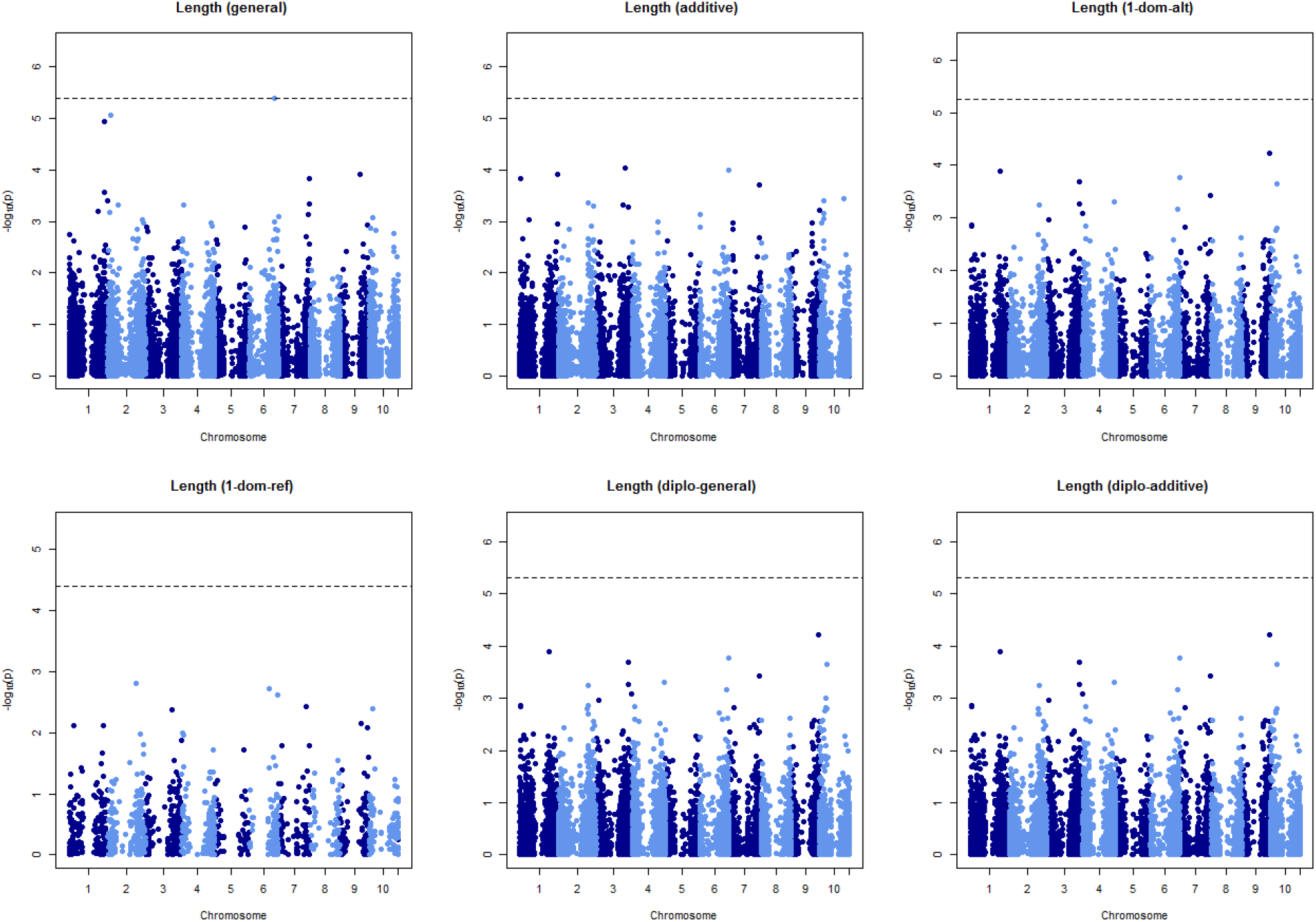
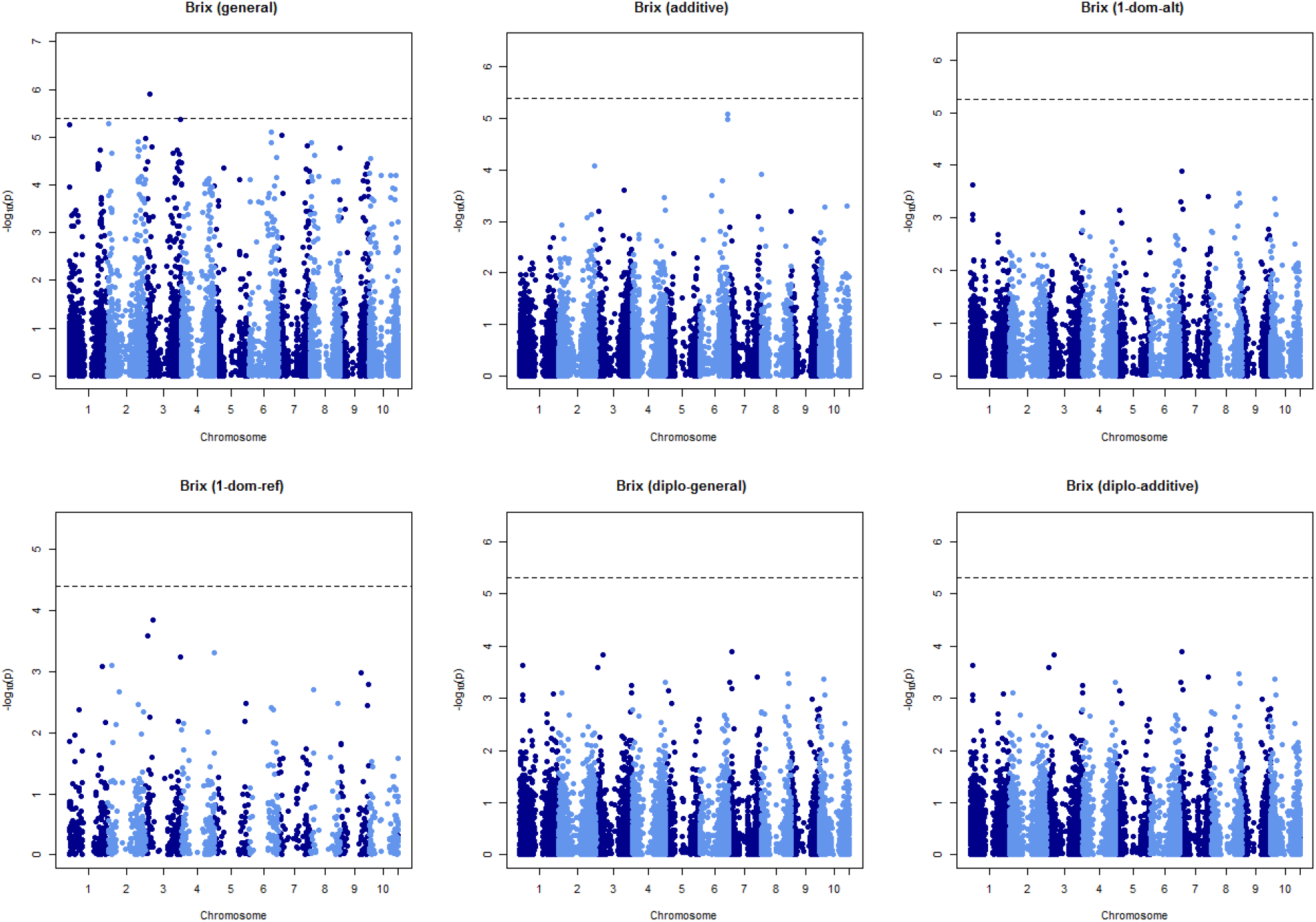
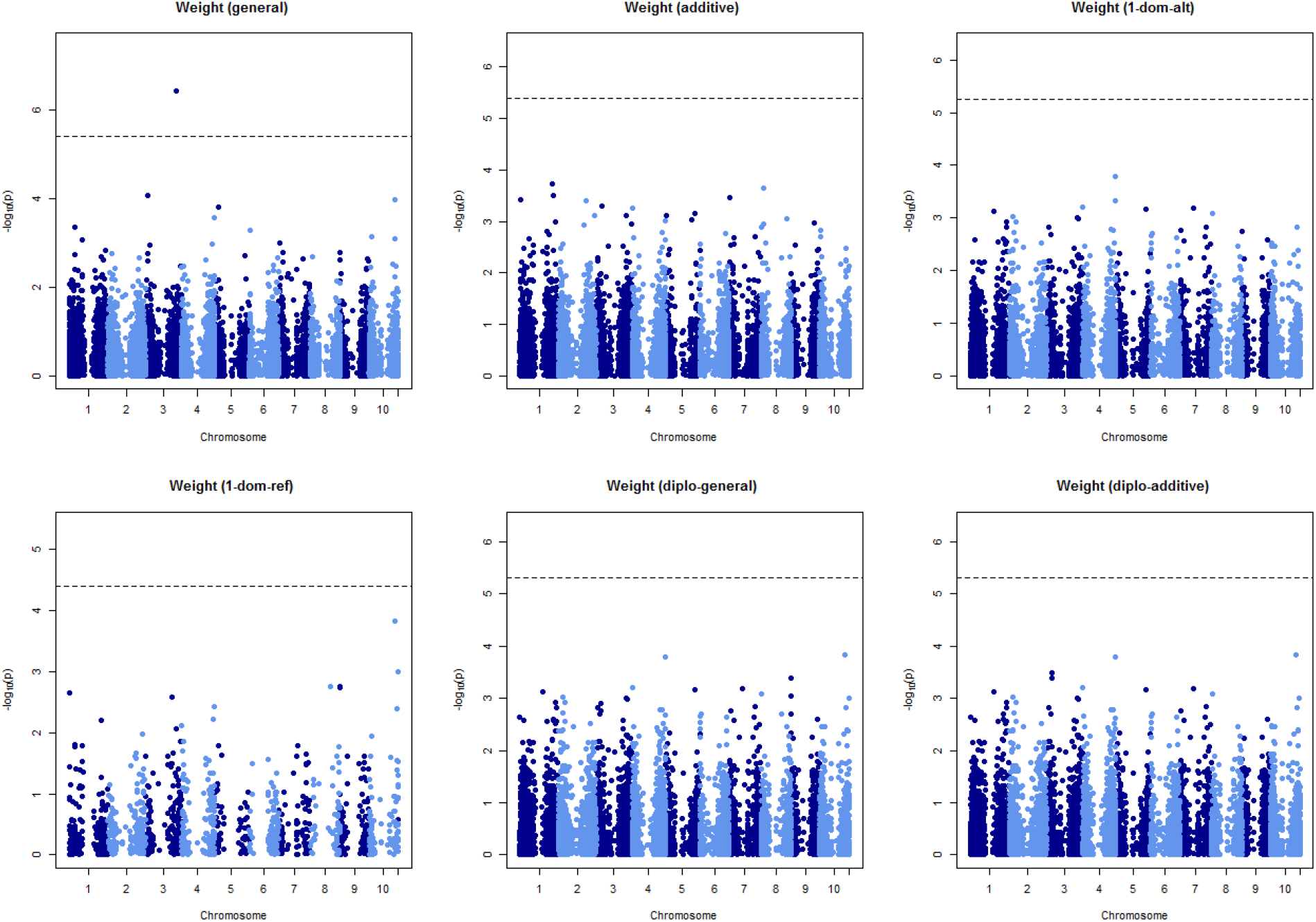
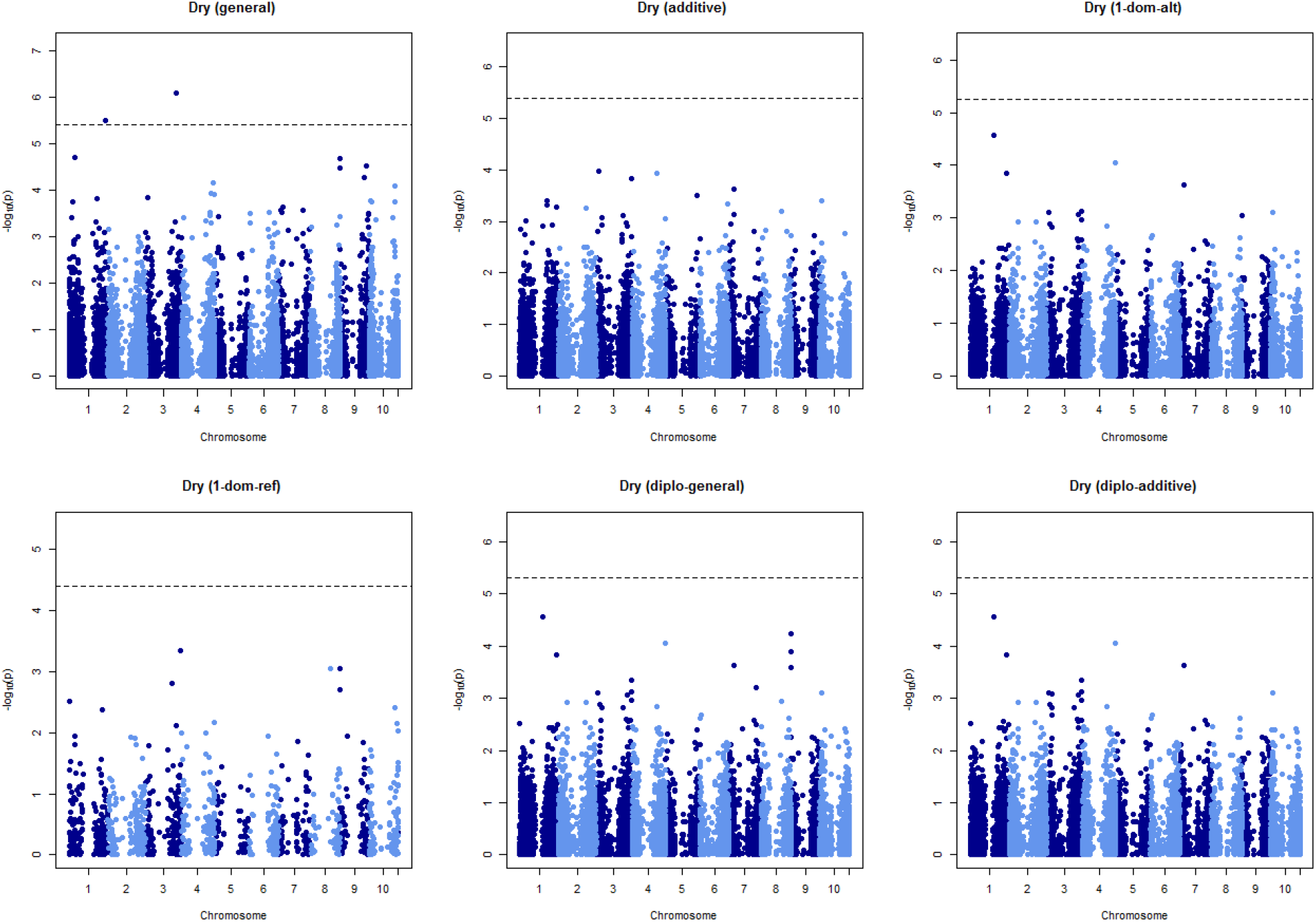
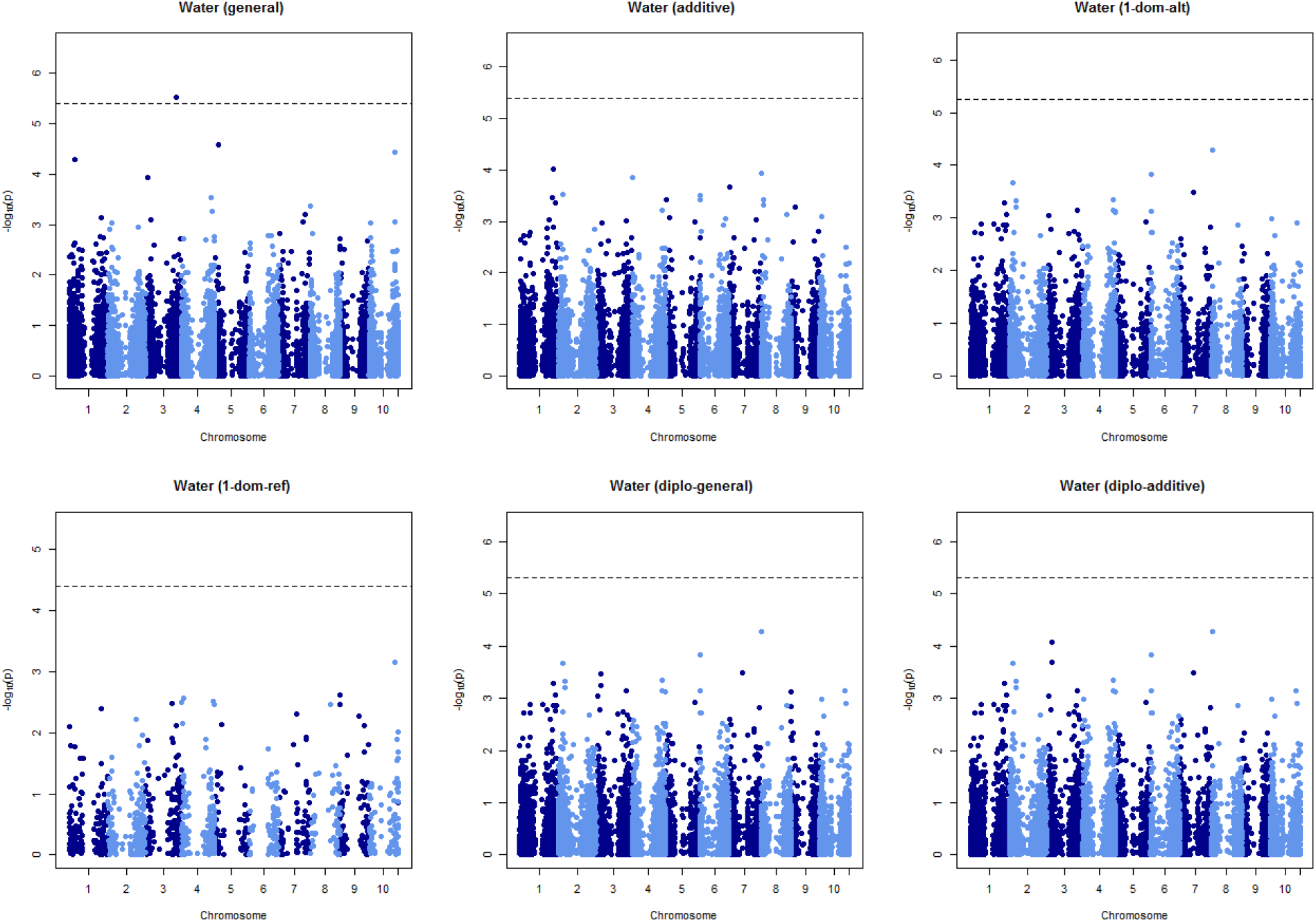
Significant marker-trait associations identified in *Saccharum spontaneum* set of the sugarcane diversity panel with 100 accessions. Stalk = Stalk number; Diameter = Stalk diameter; Width = Leaf width; Length = Leaf length; Weight = Total weight; Dry = Dry weight; Water = Water content; Internode = Internode length.

**Figure S5.**
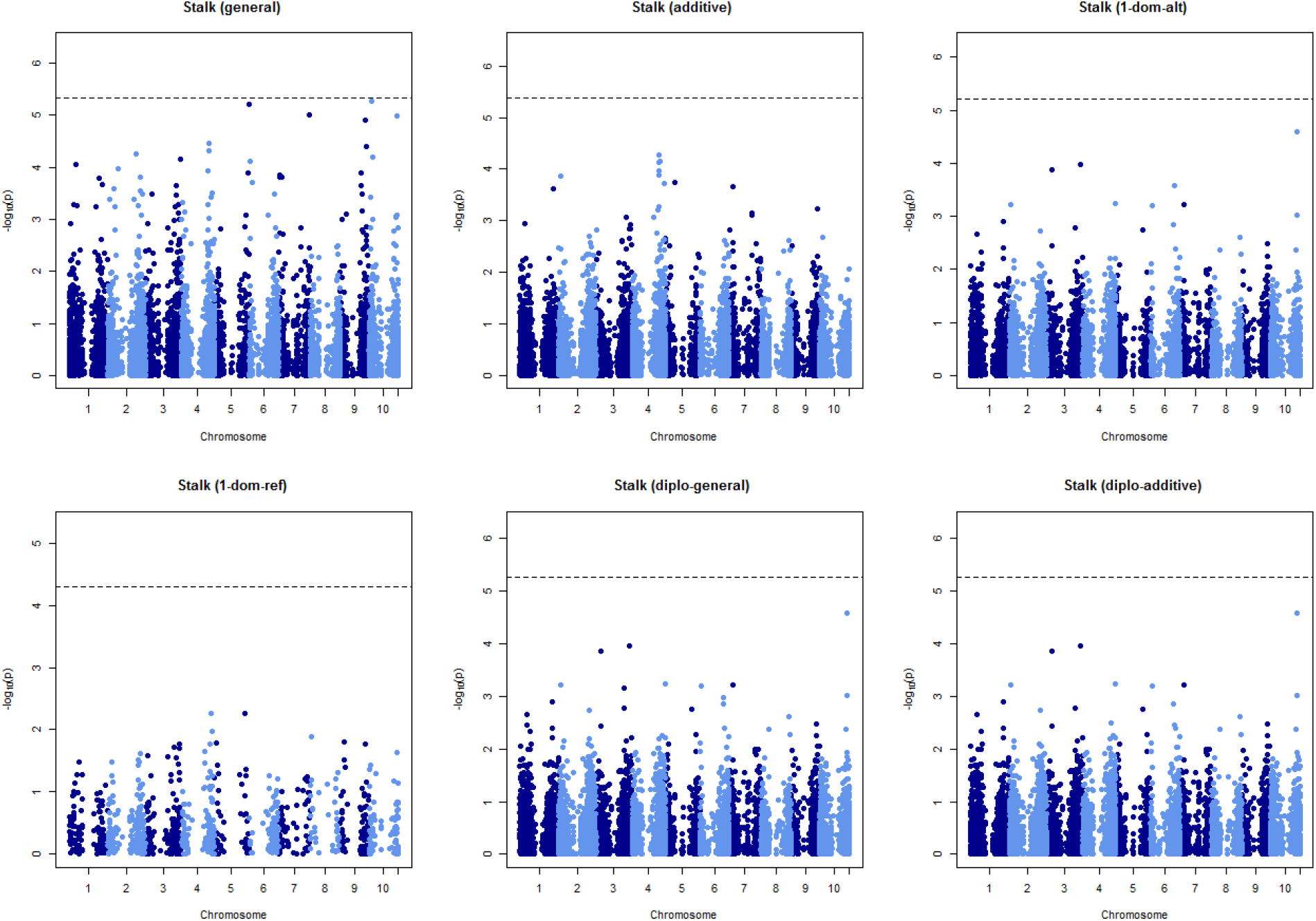
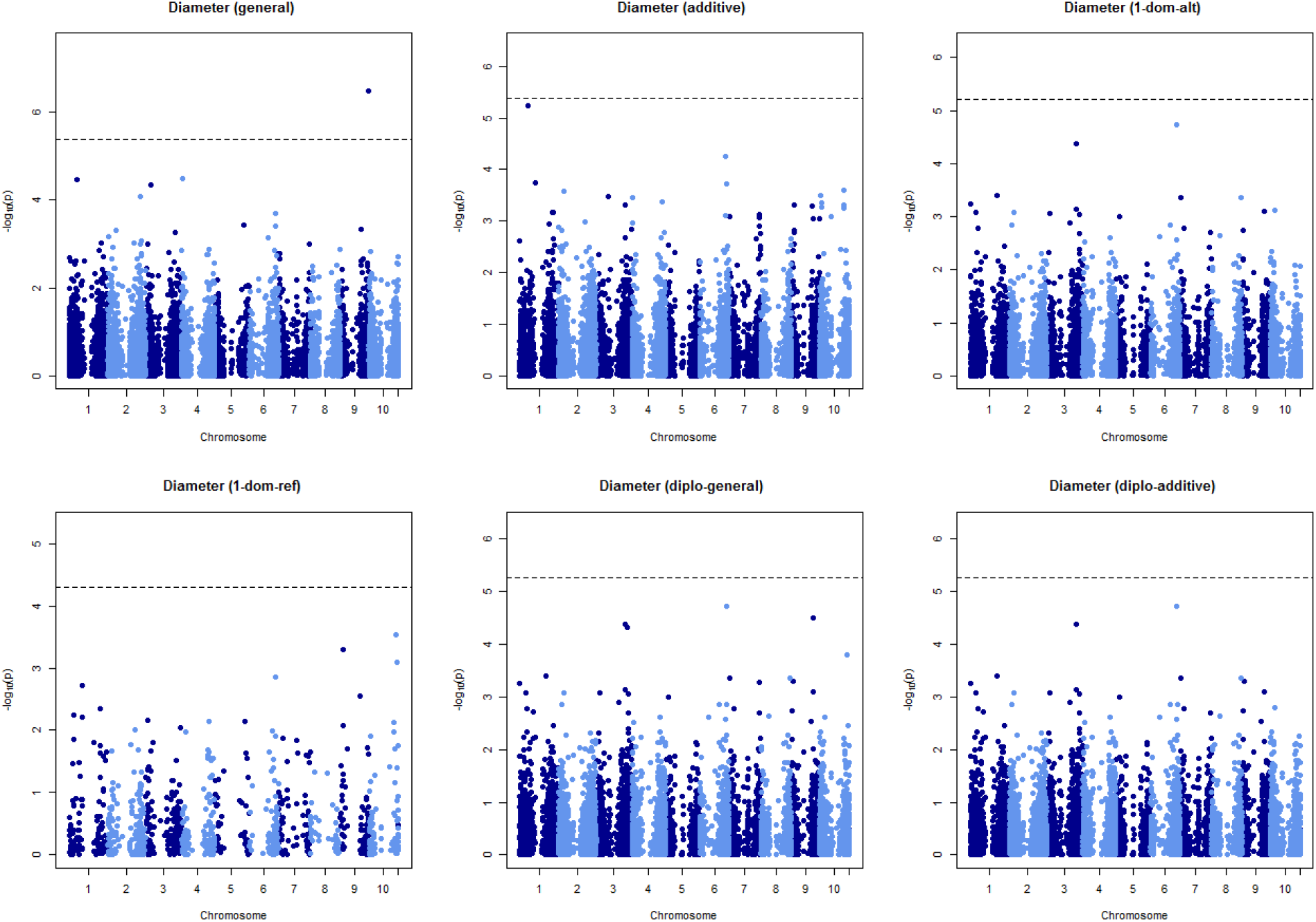
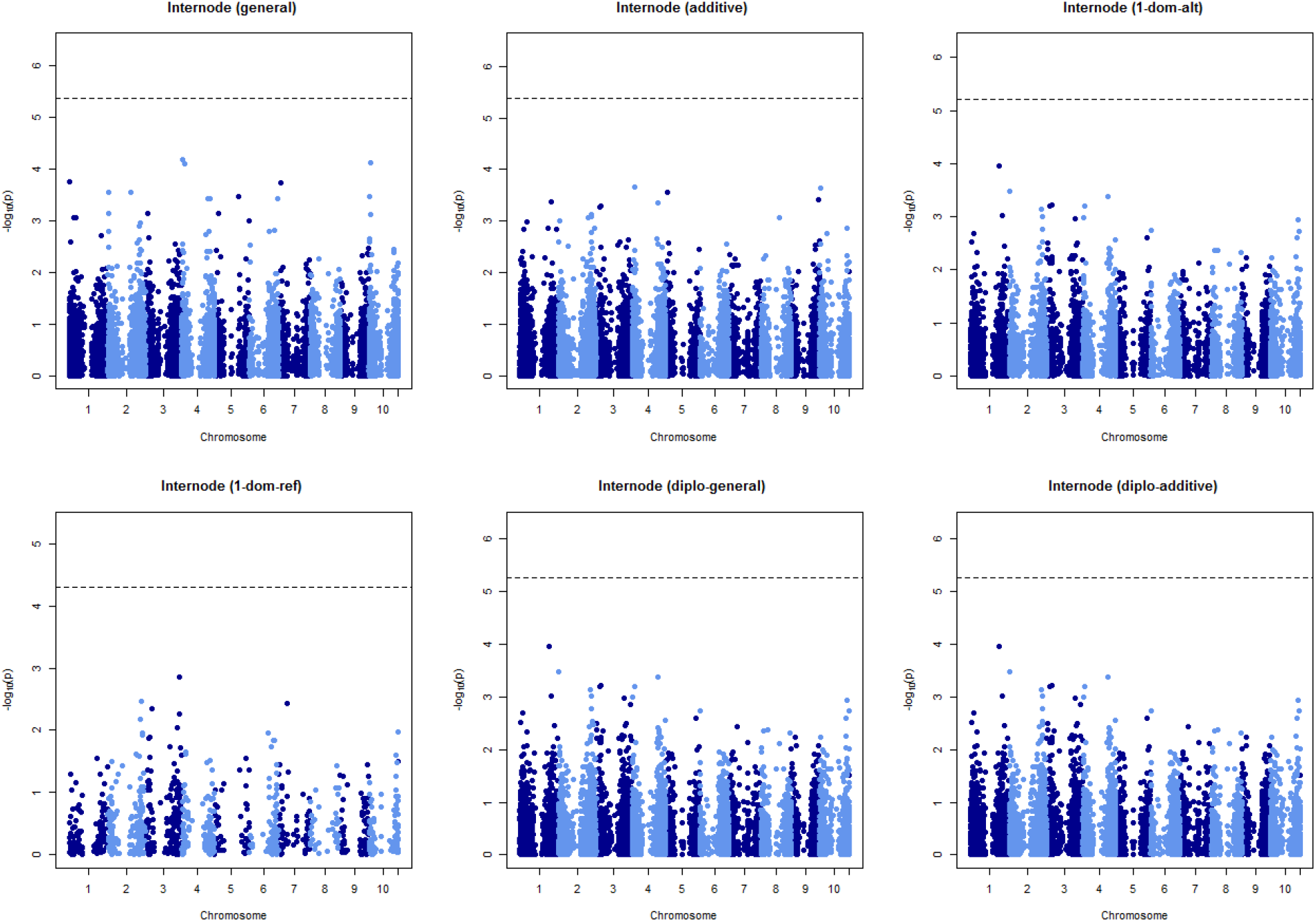
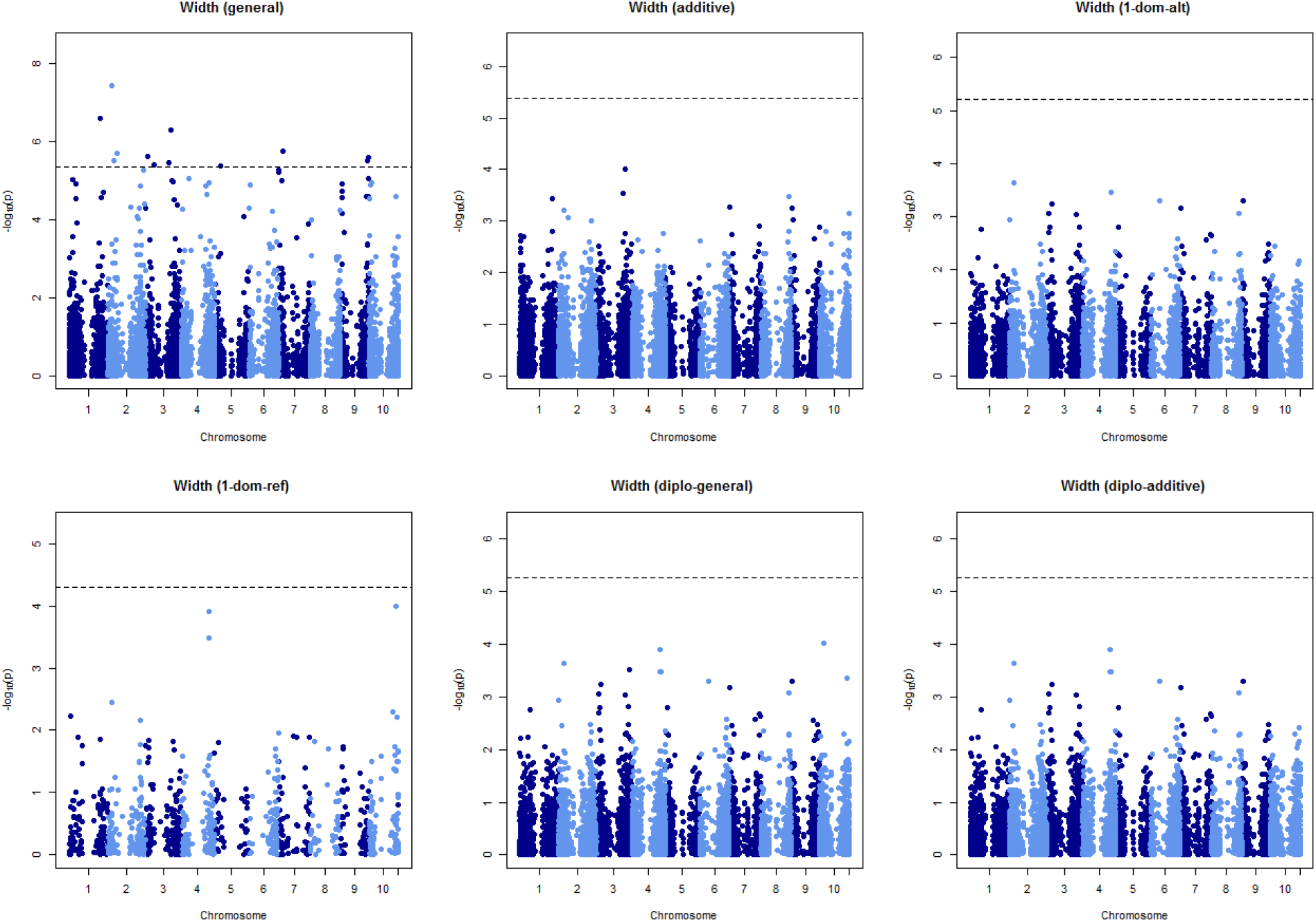
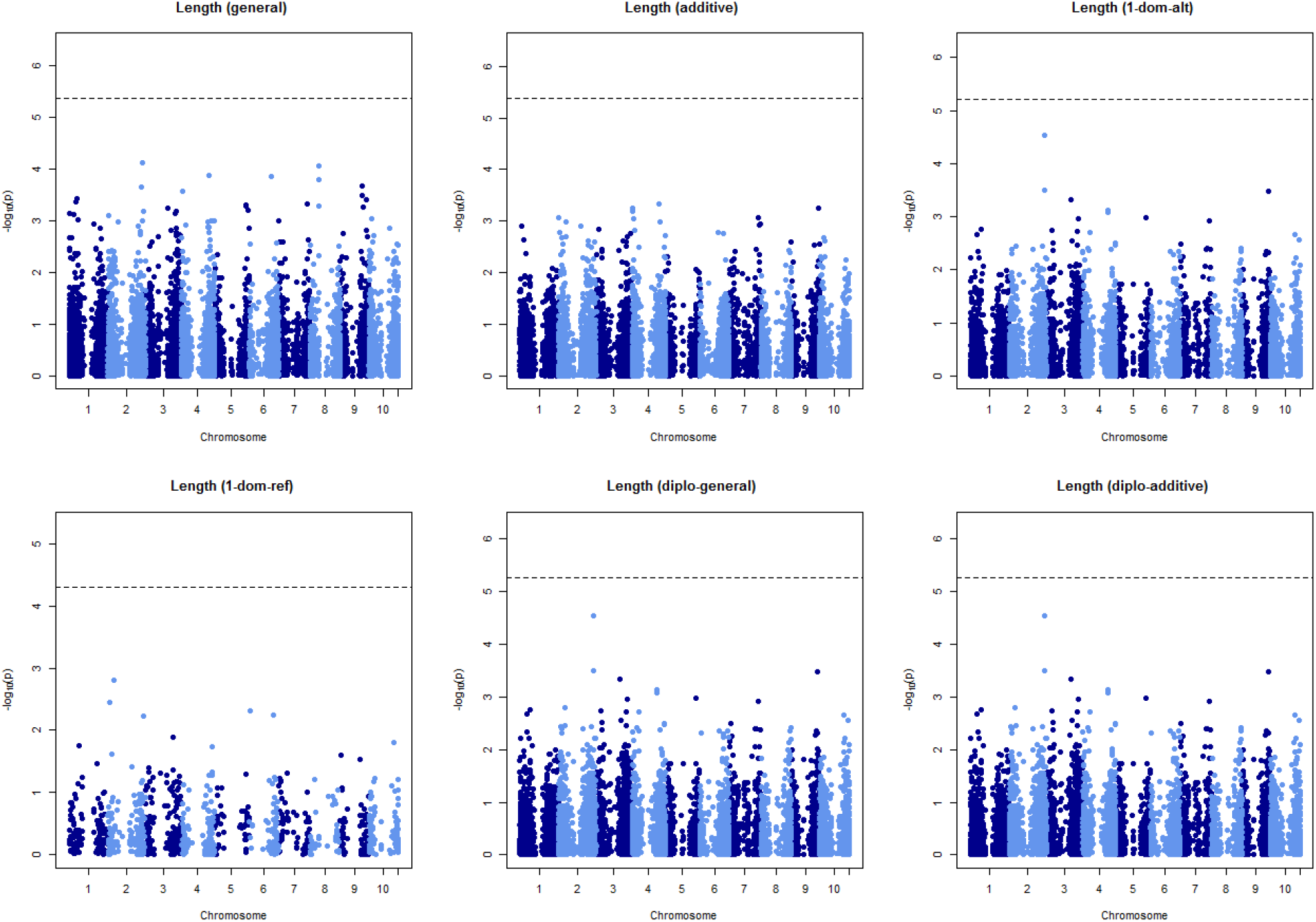
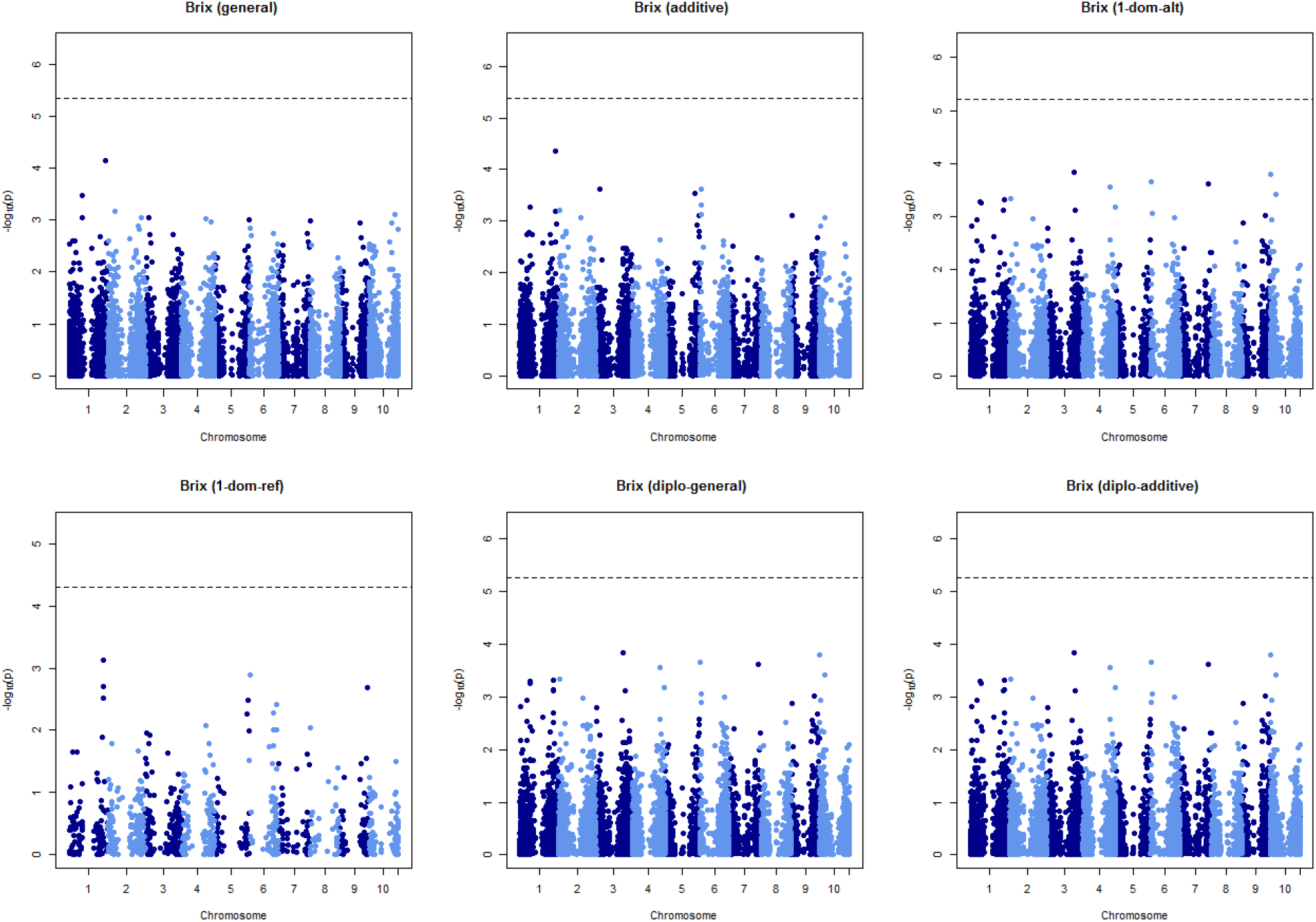
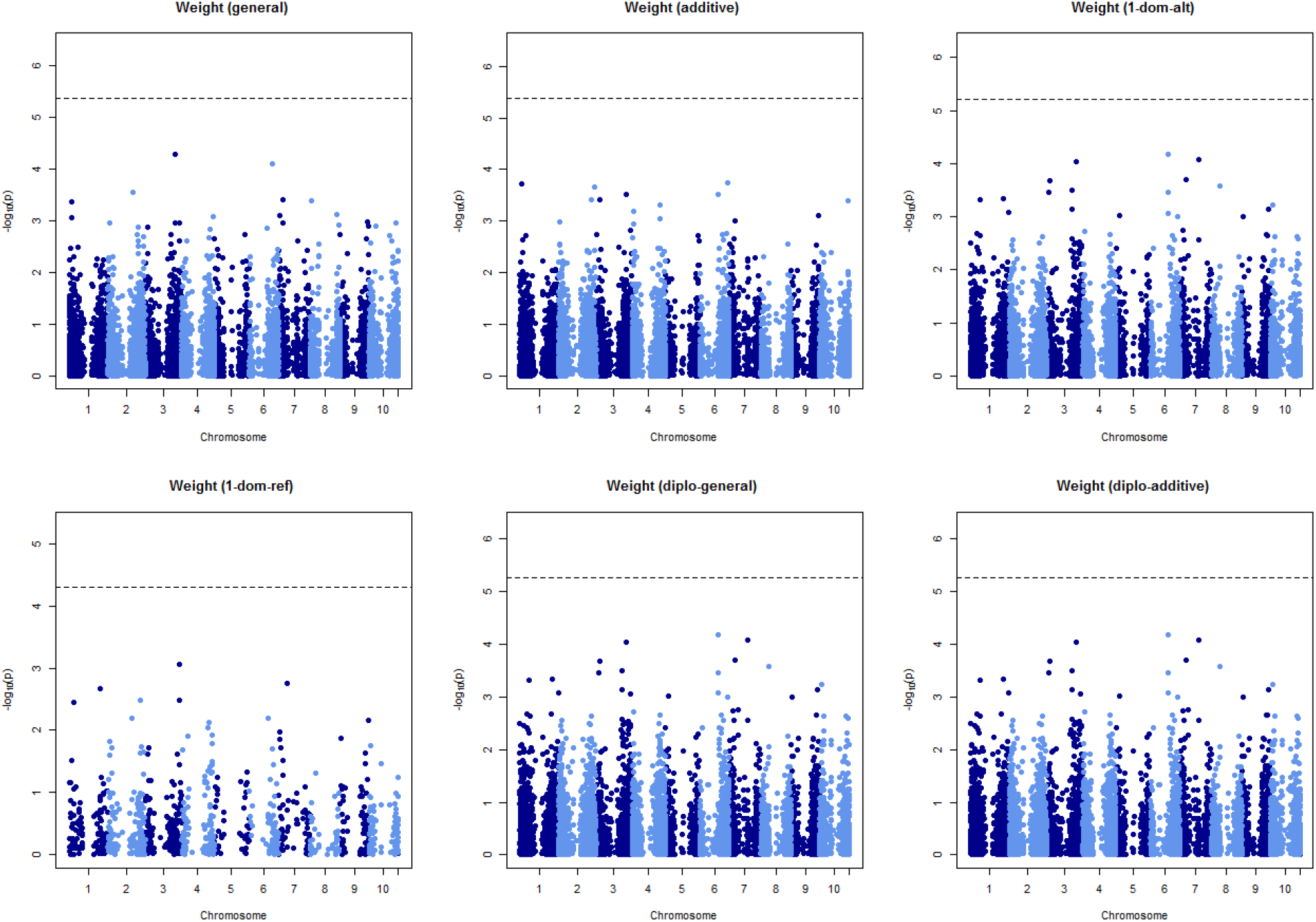
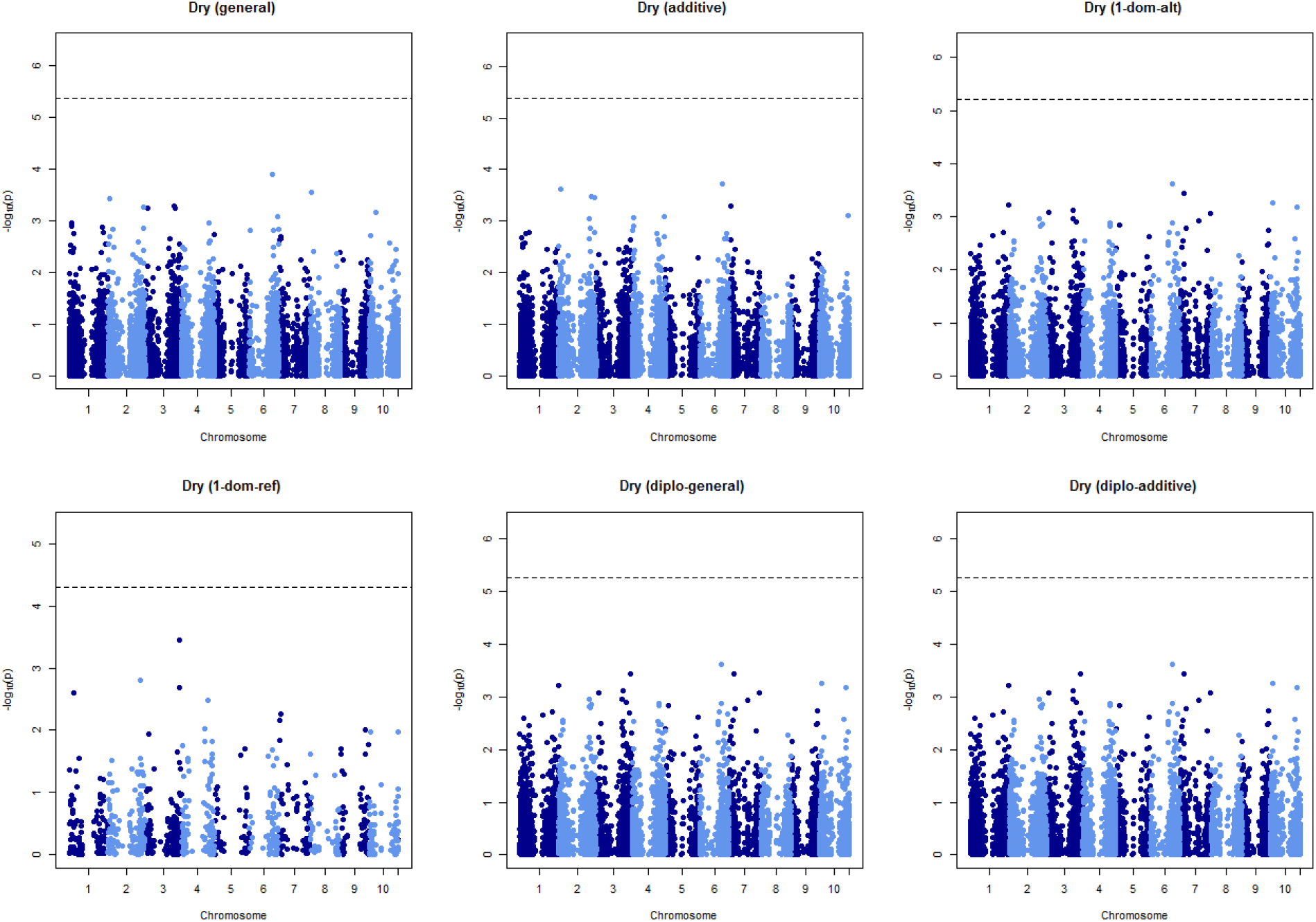
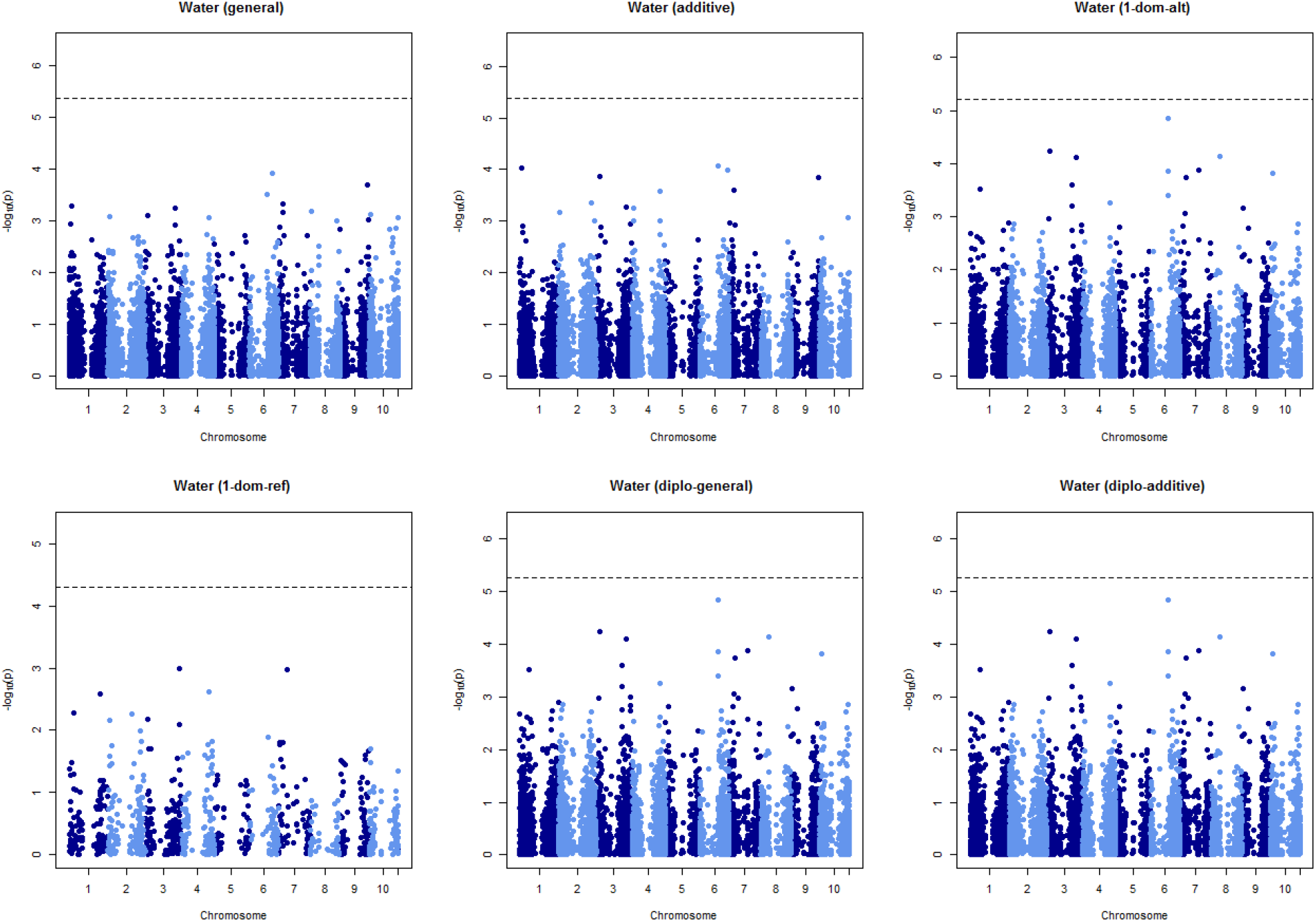
Significant marker-trait associations identified in sugarcane hybrid set of the sugarcane diversity panel with 173 accessions. Stalk = Stalk number; Diameter = Stalk diameter; Width = Leaf width; Length = Leaf length; Weight = Total weight; Dry = Dry weight; Water = Water content; Internode = Internode length.

**Figure S6.**
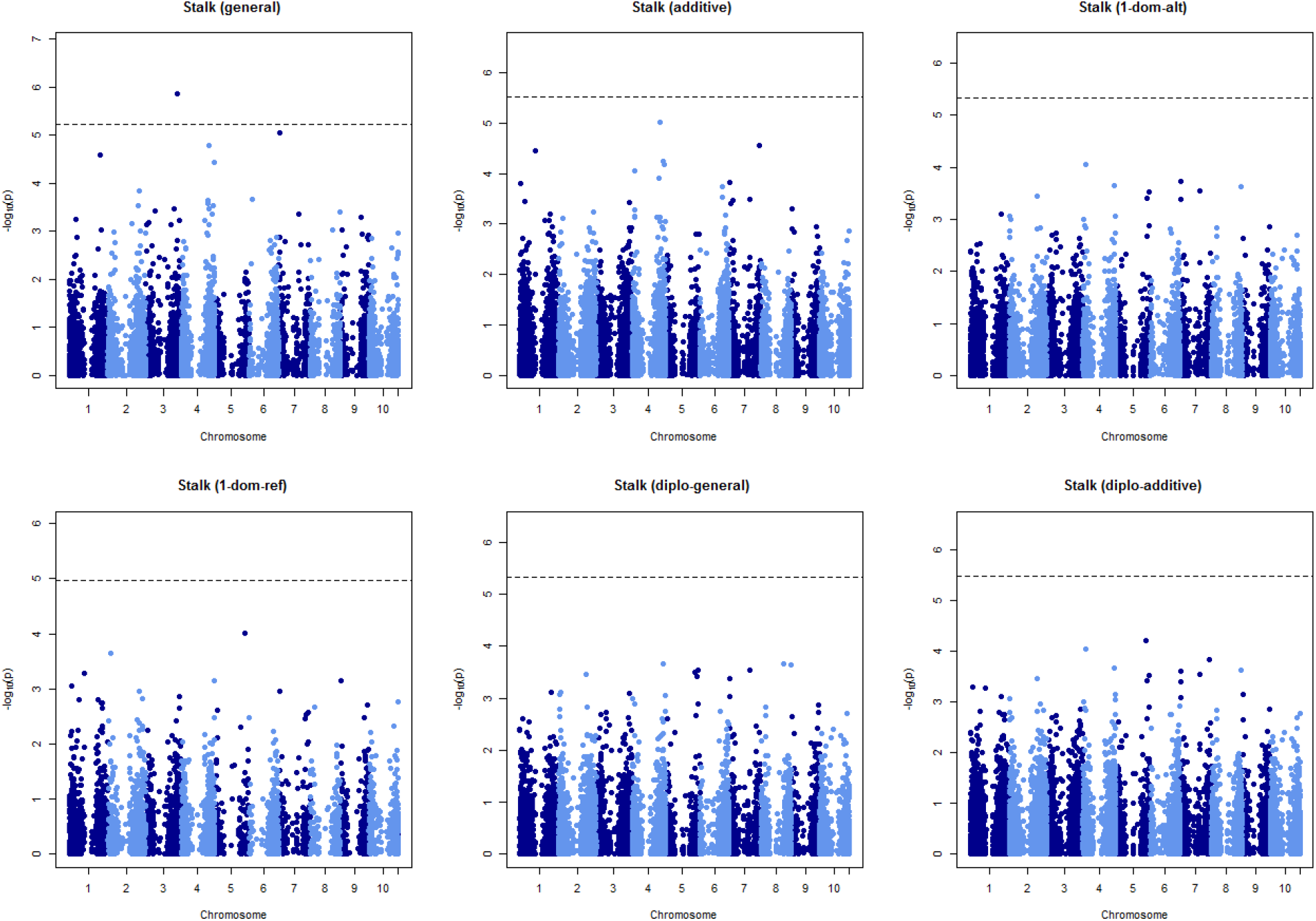
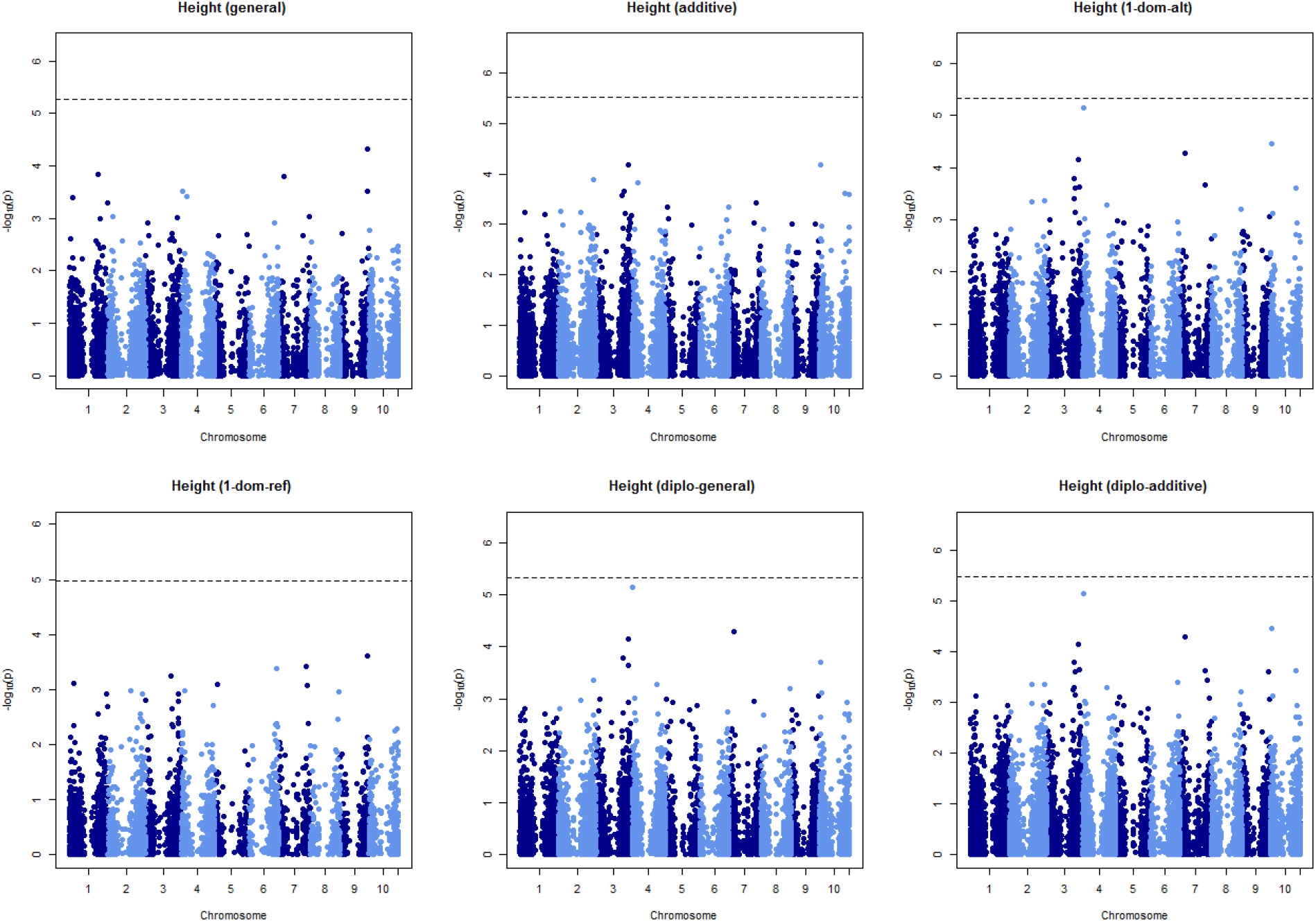
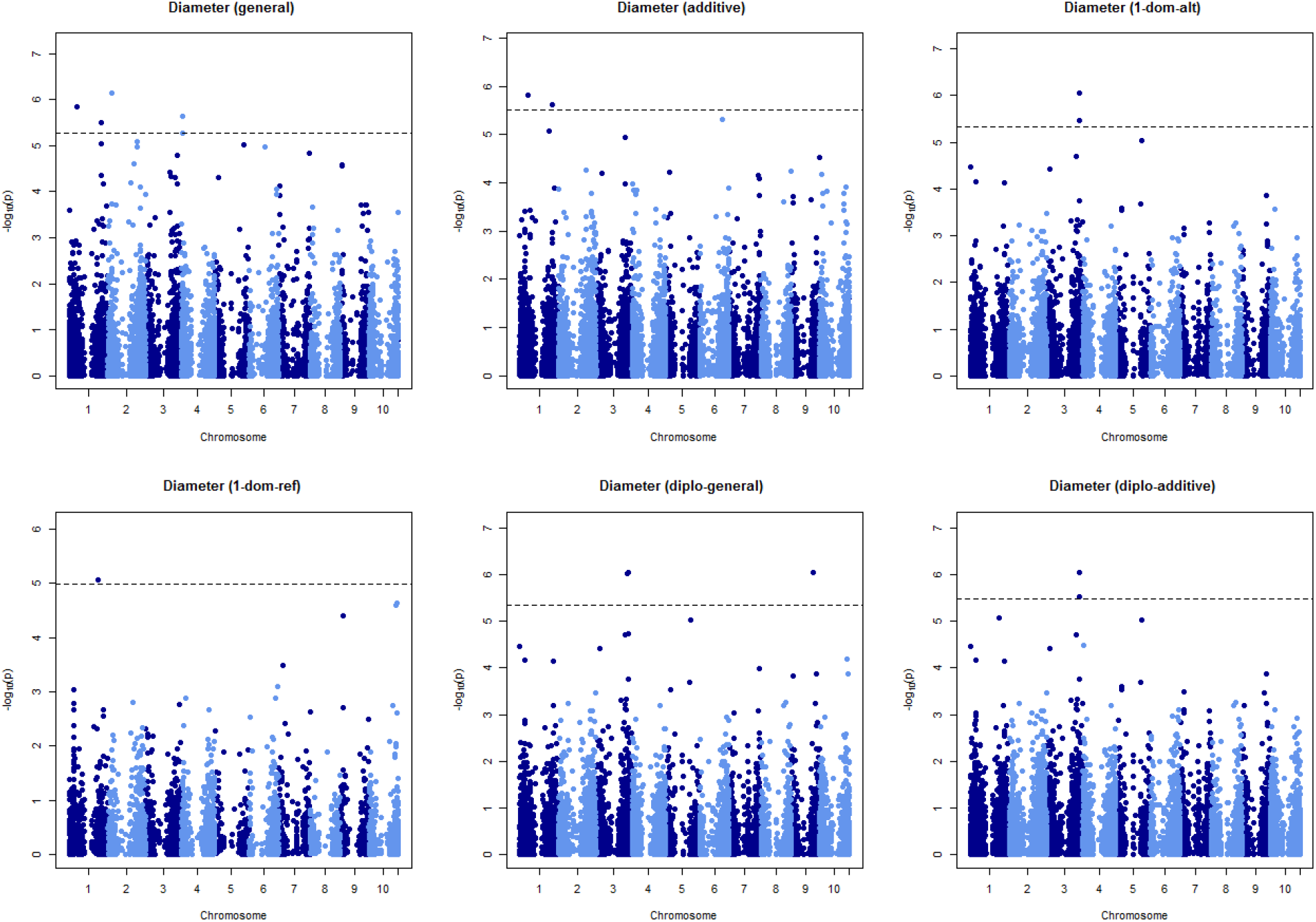
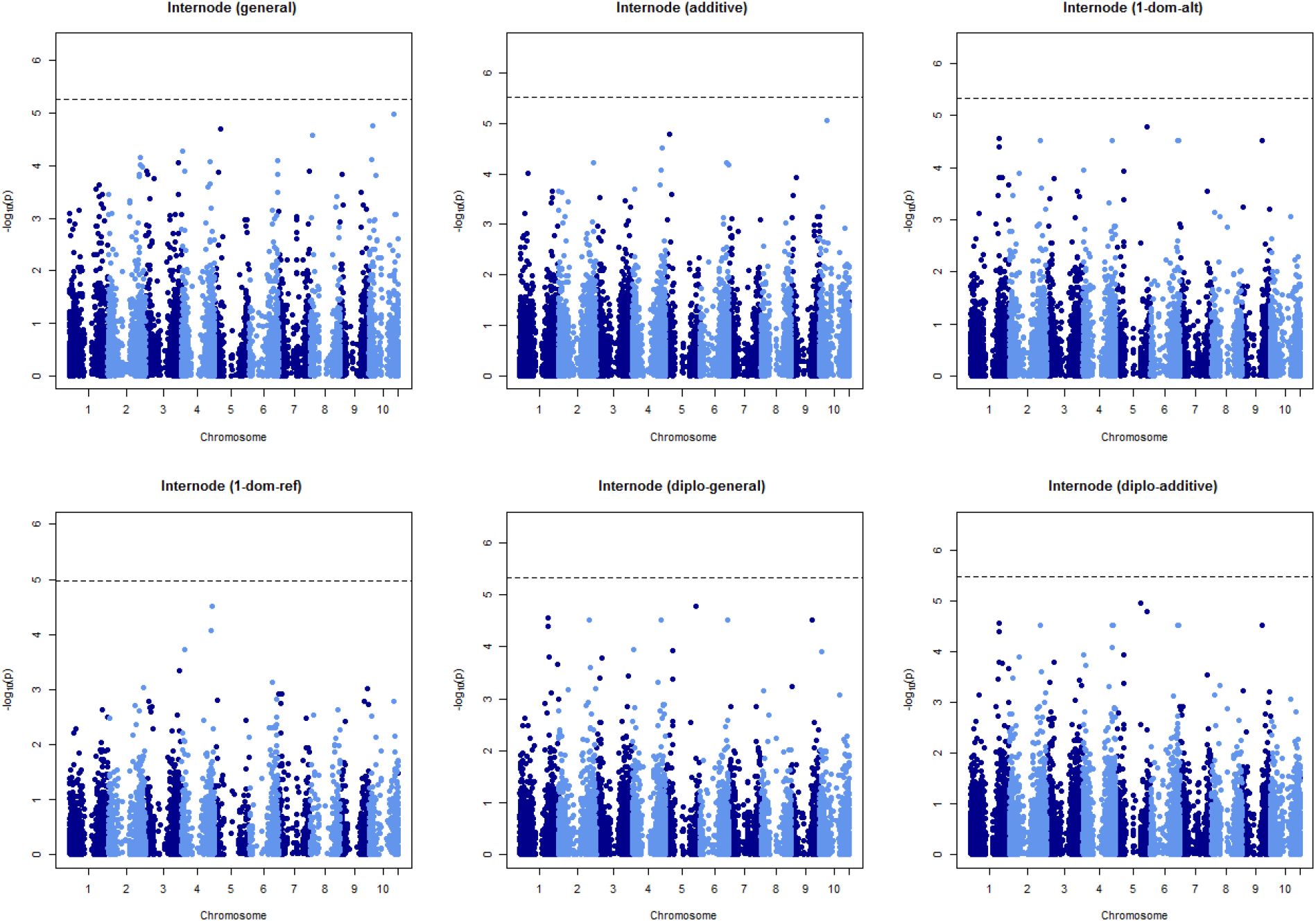
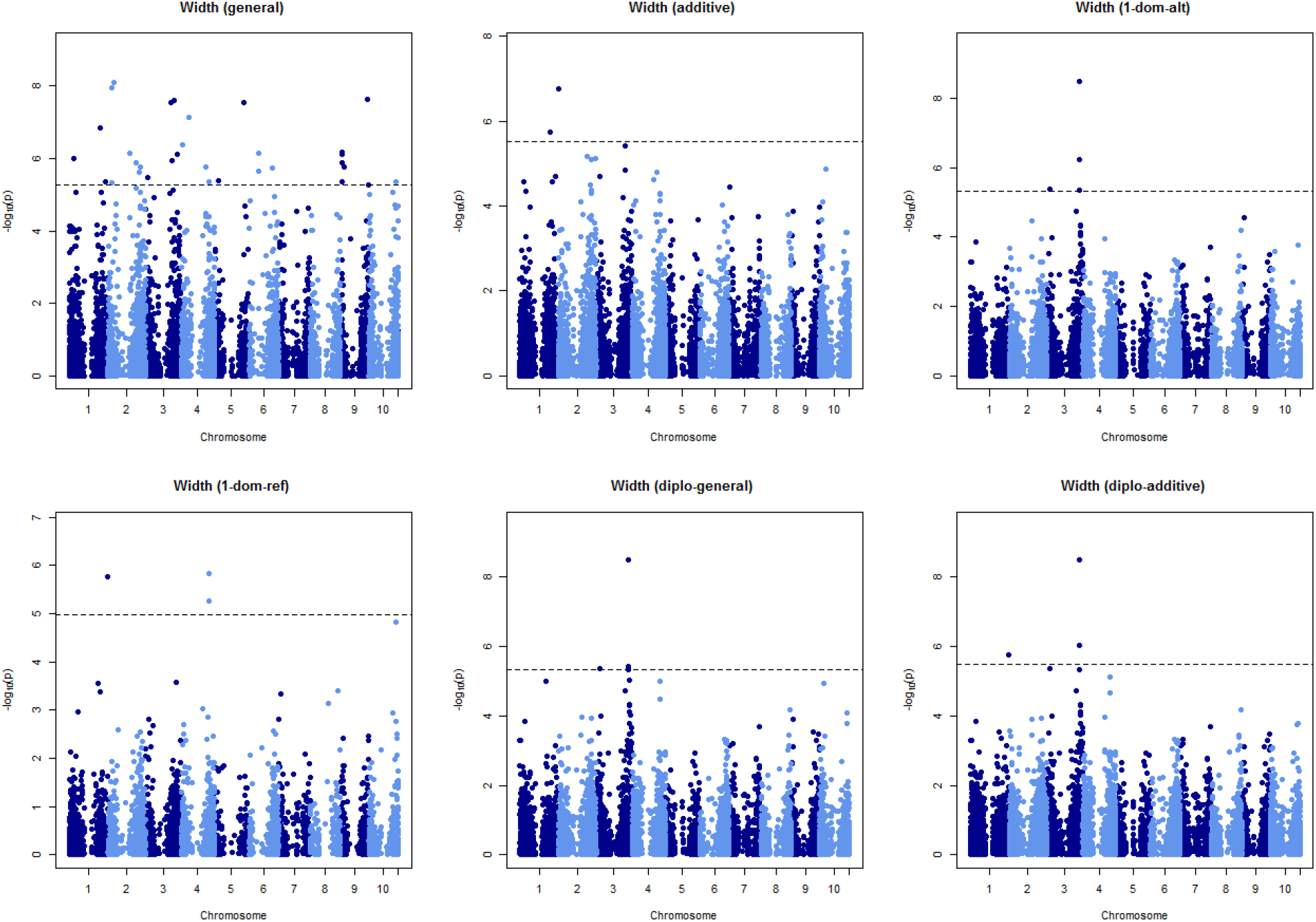
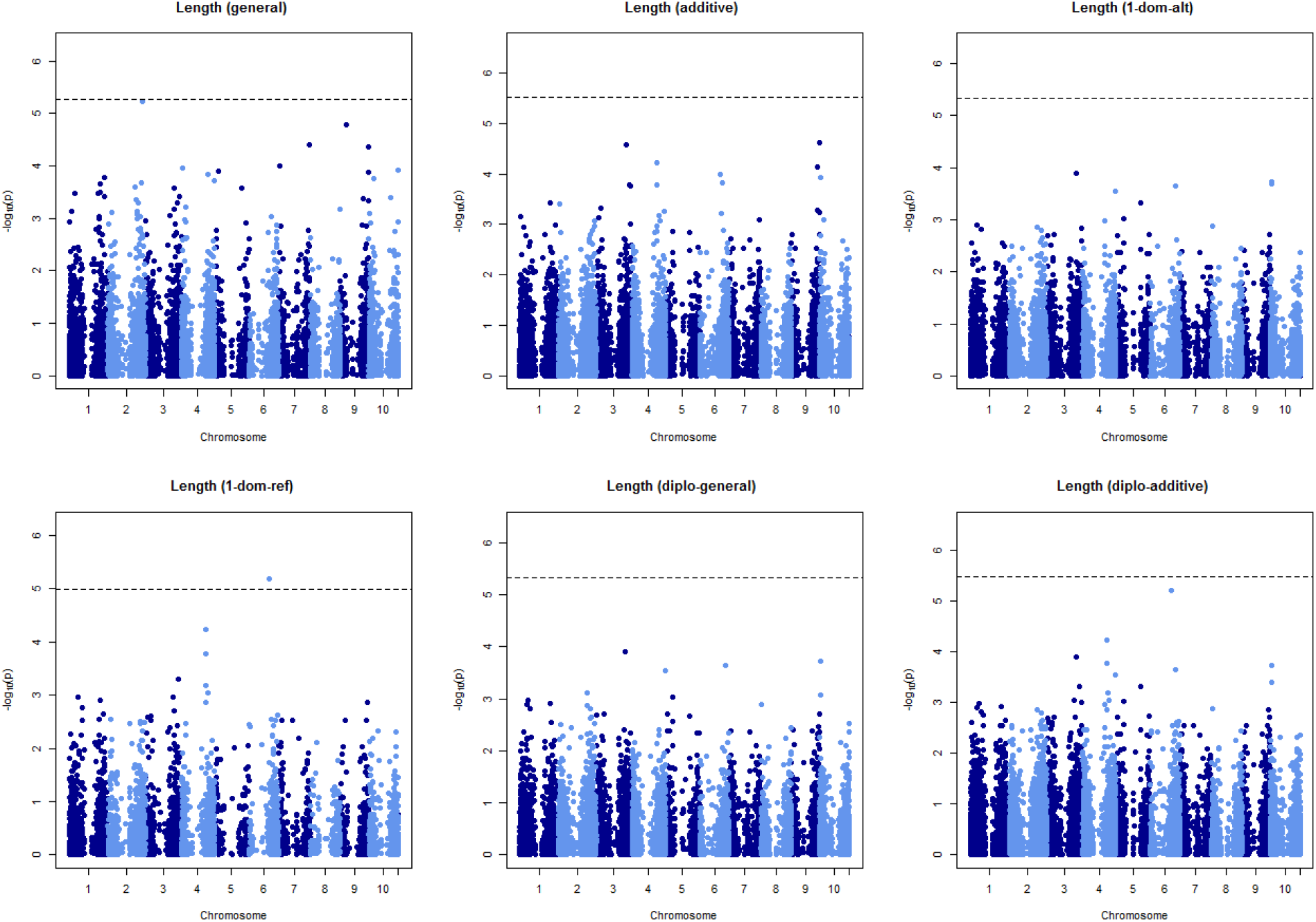
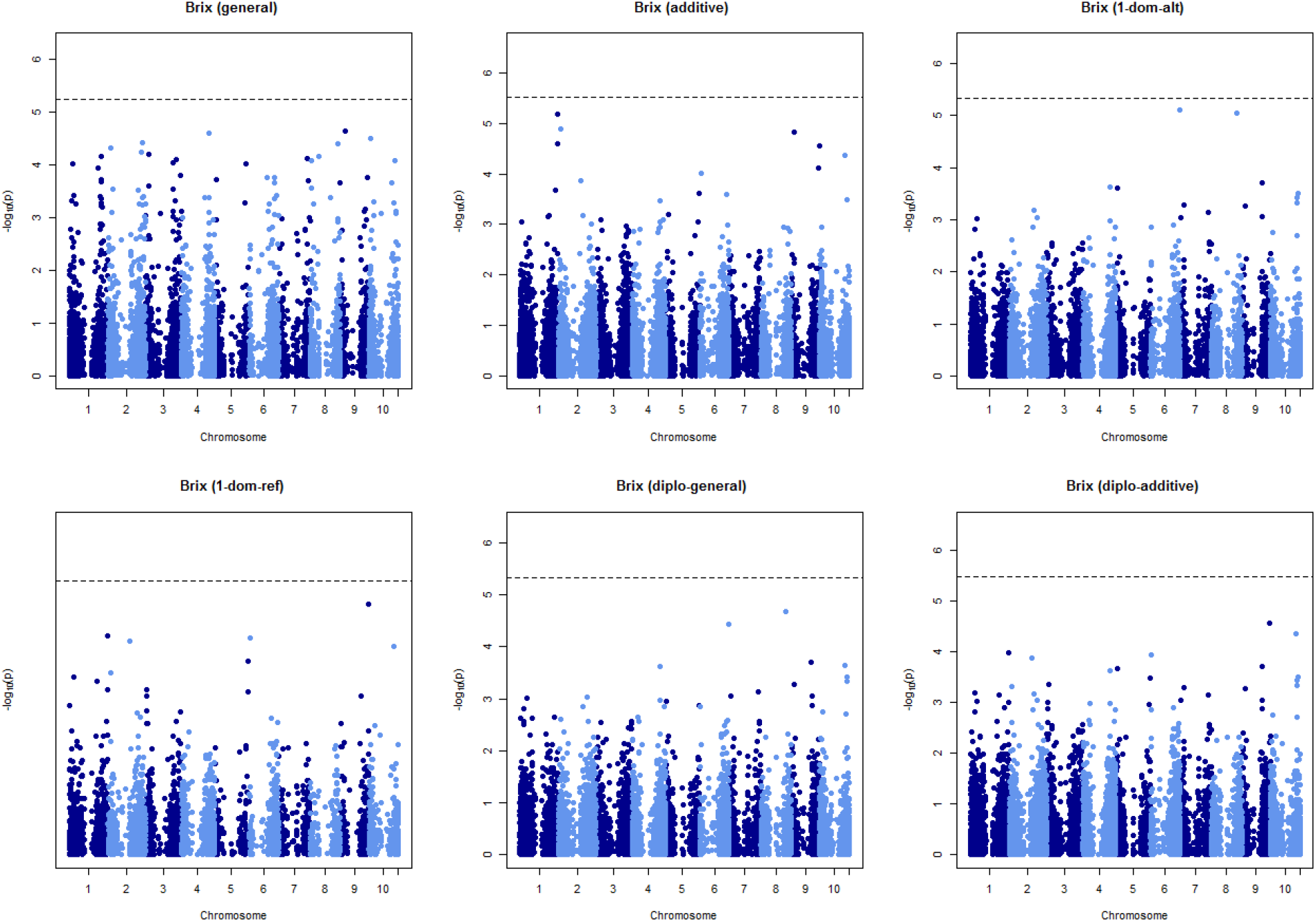
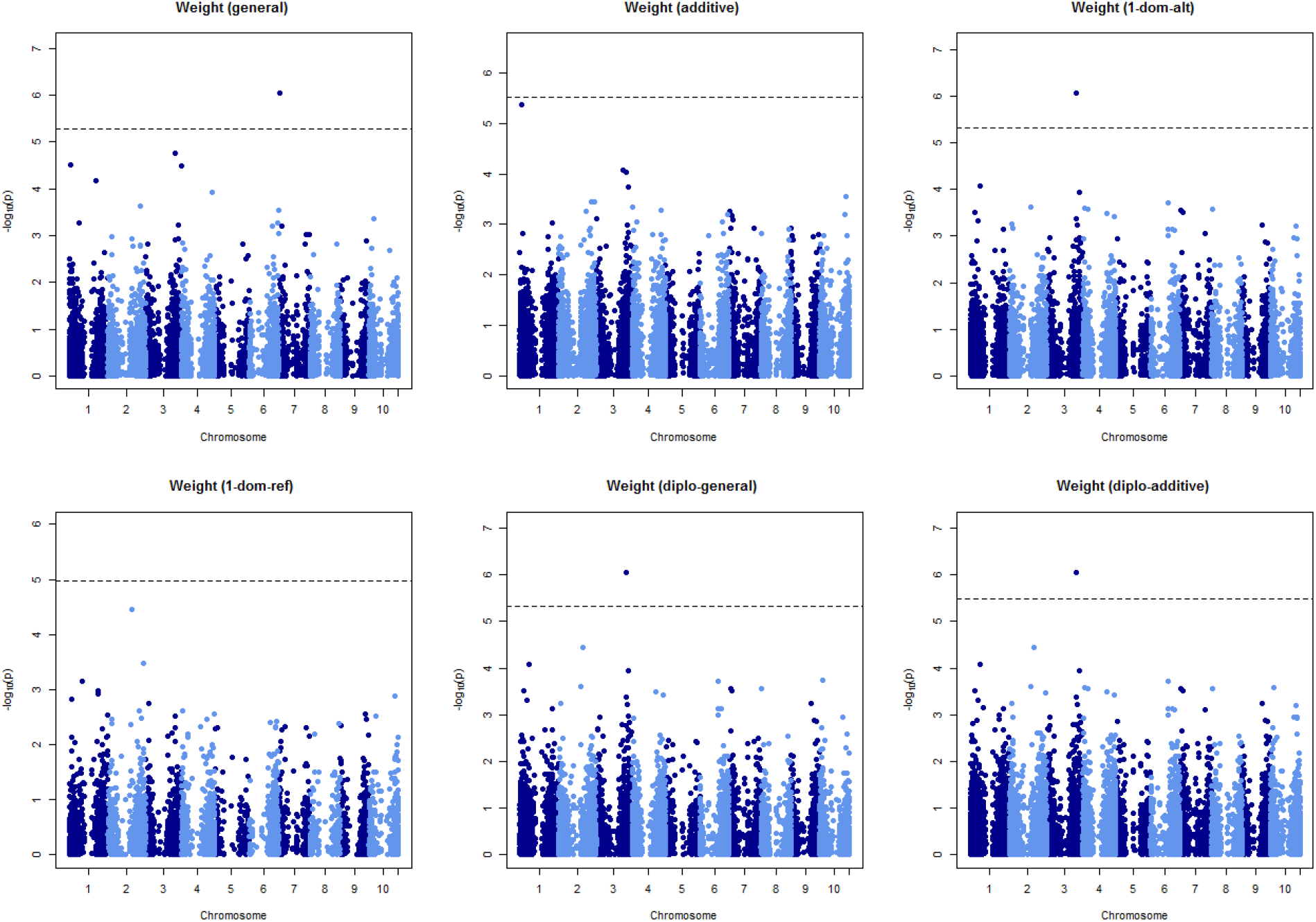
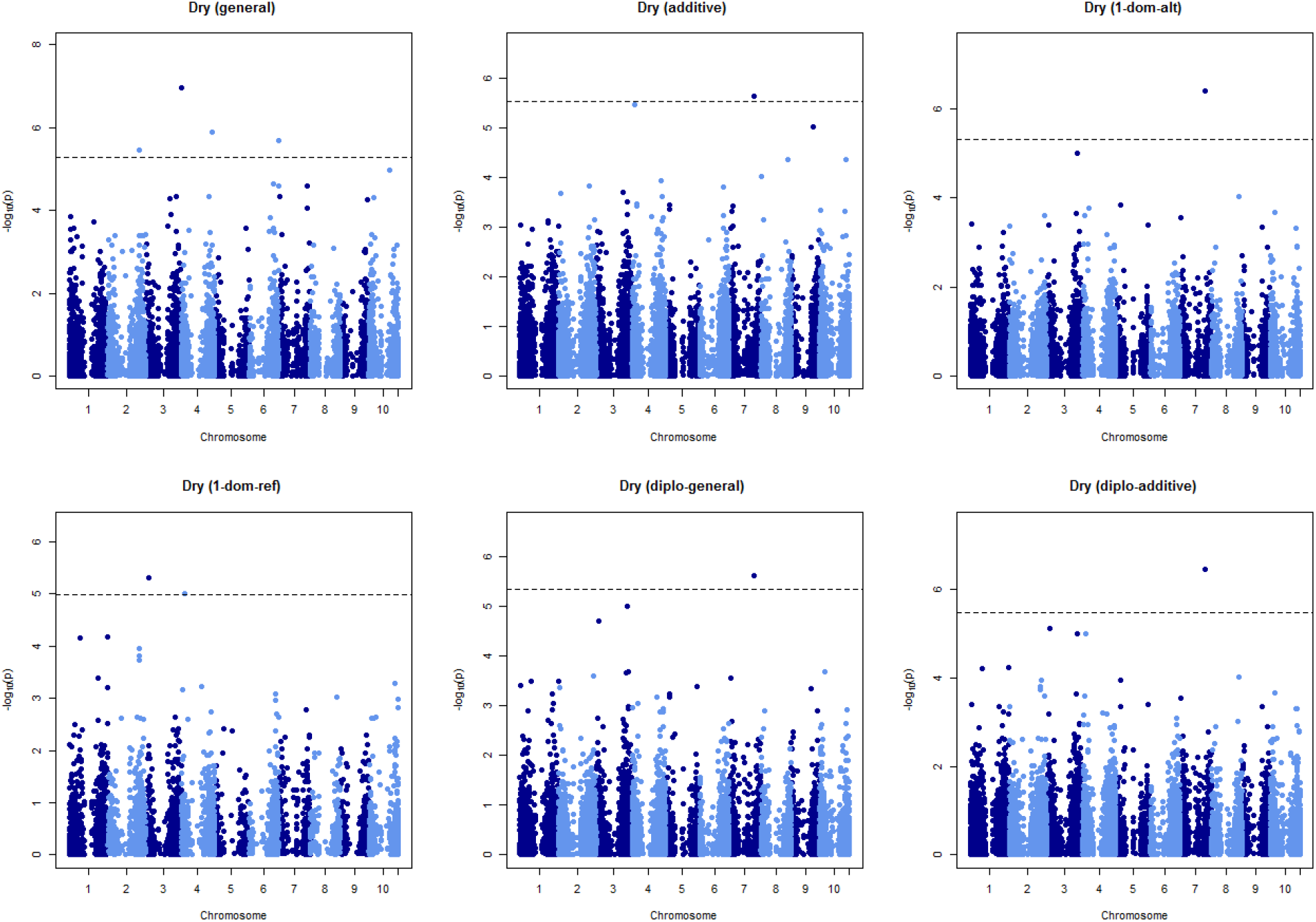
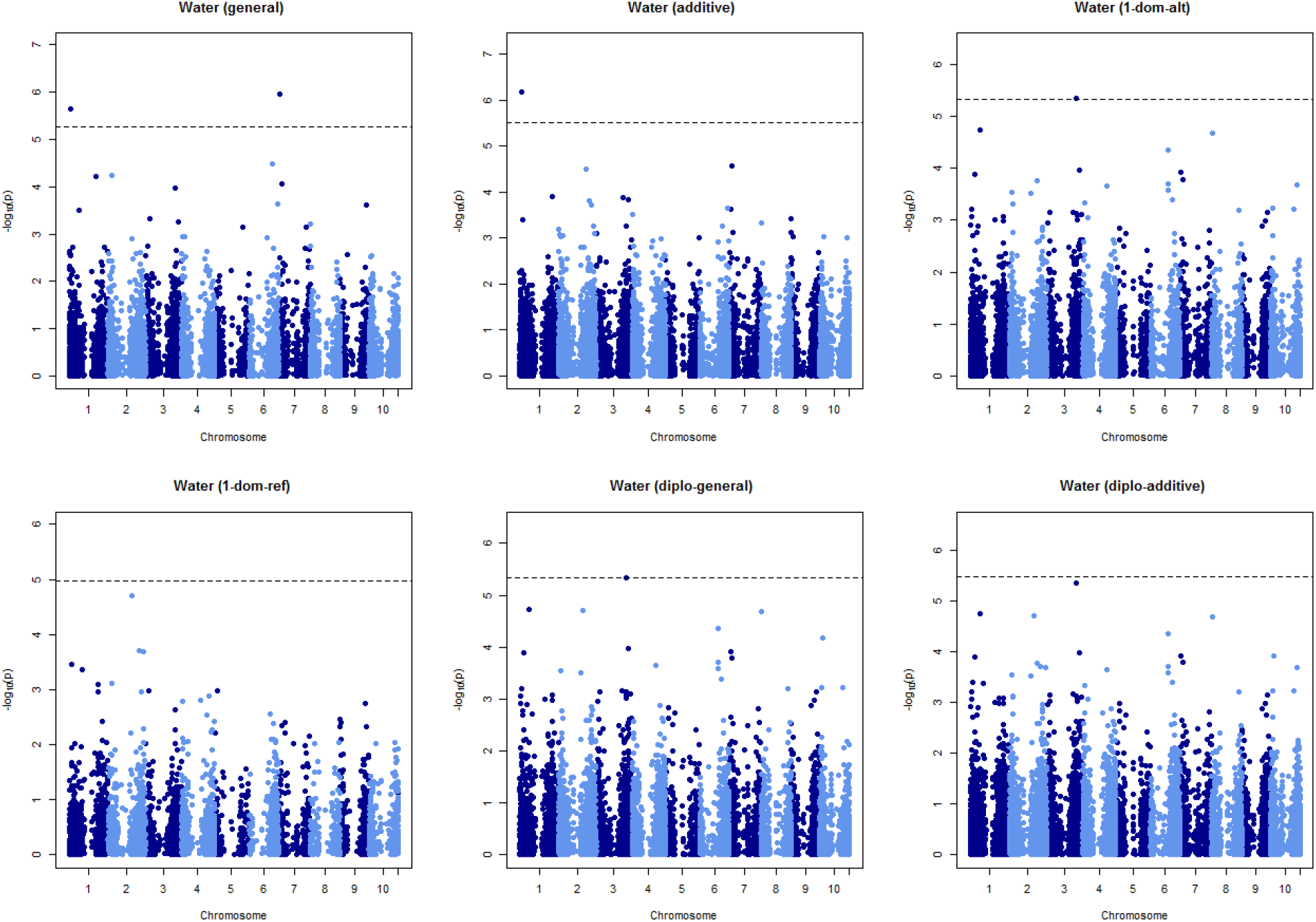
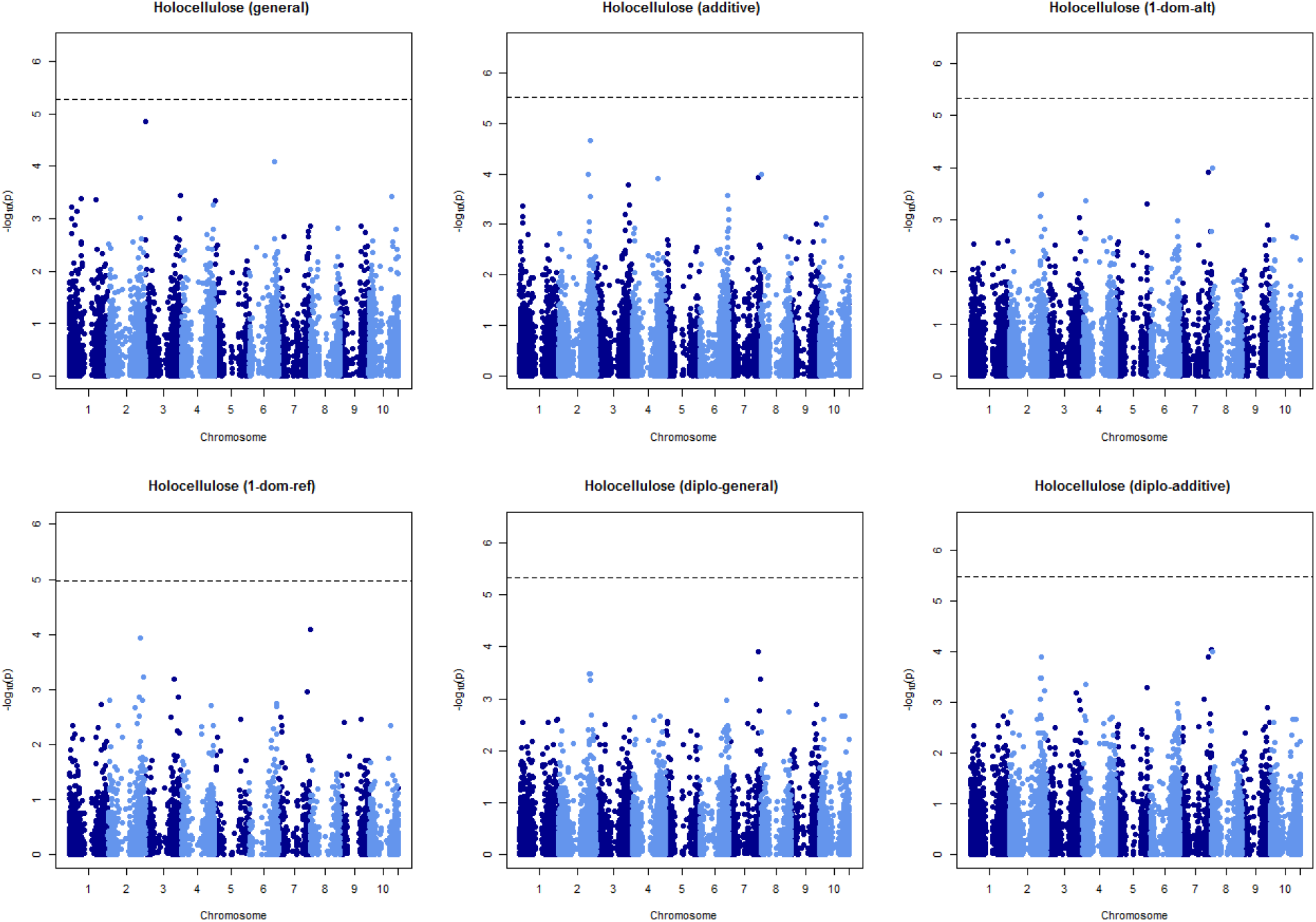
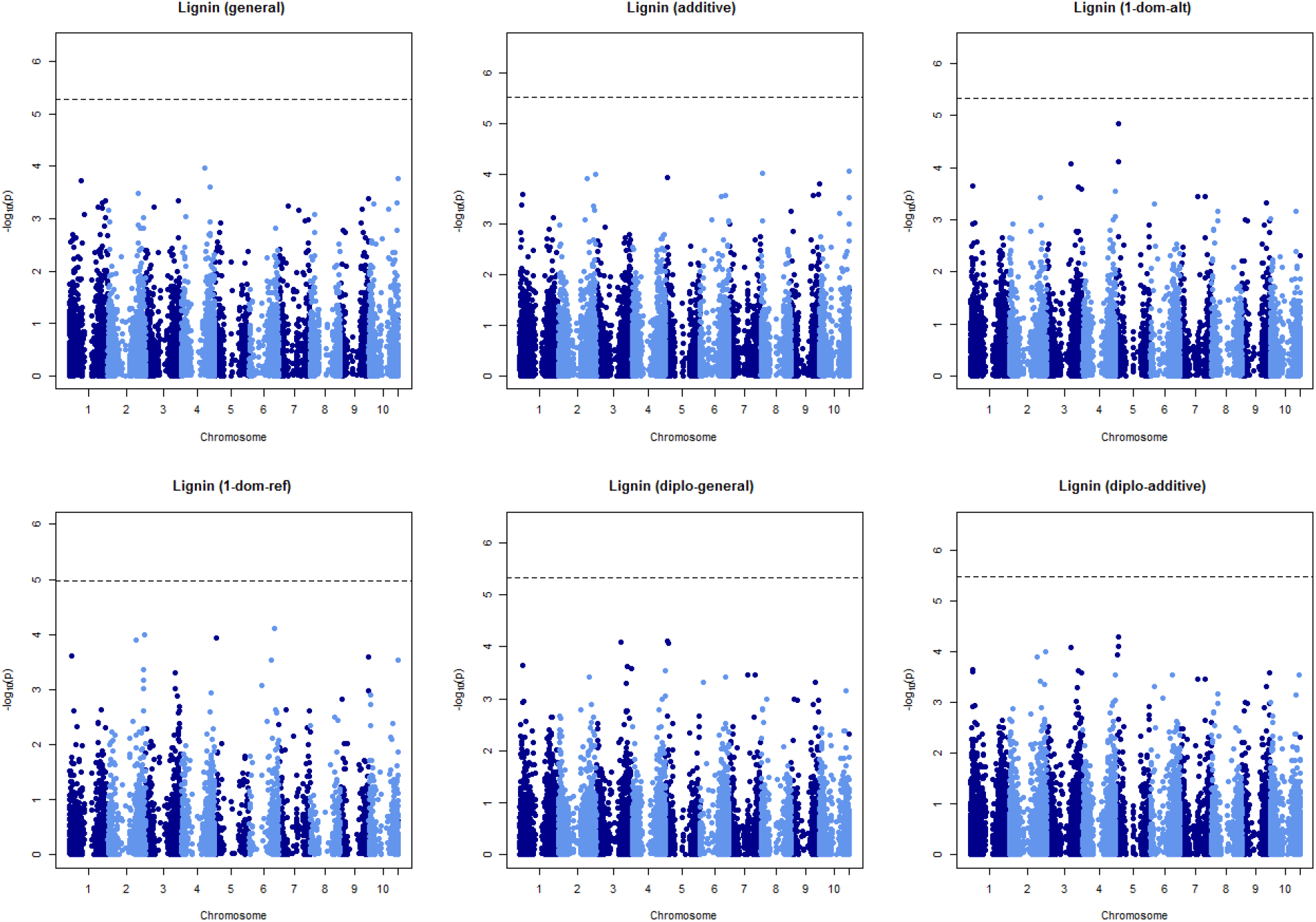
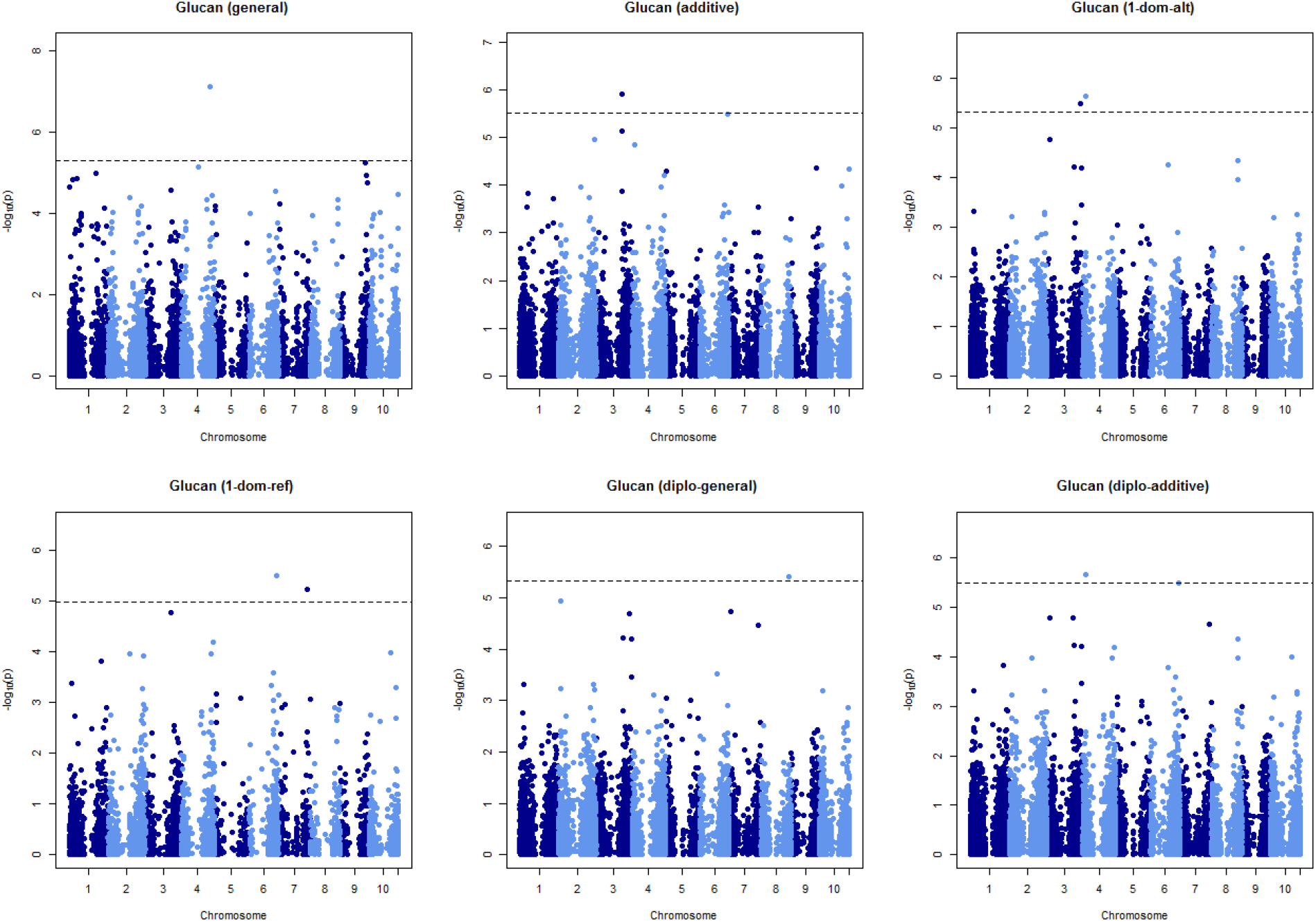
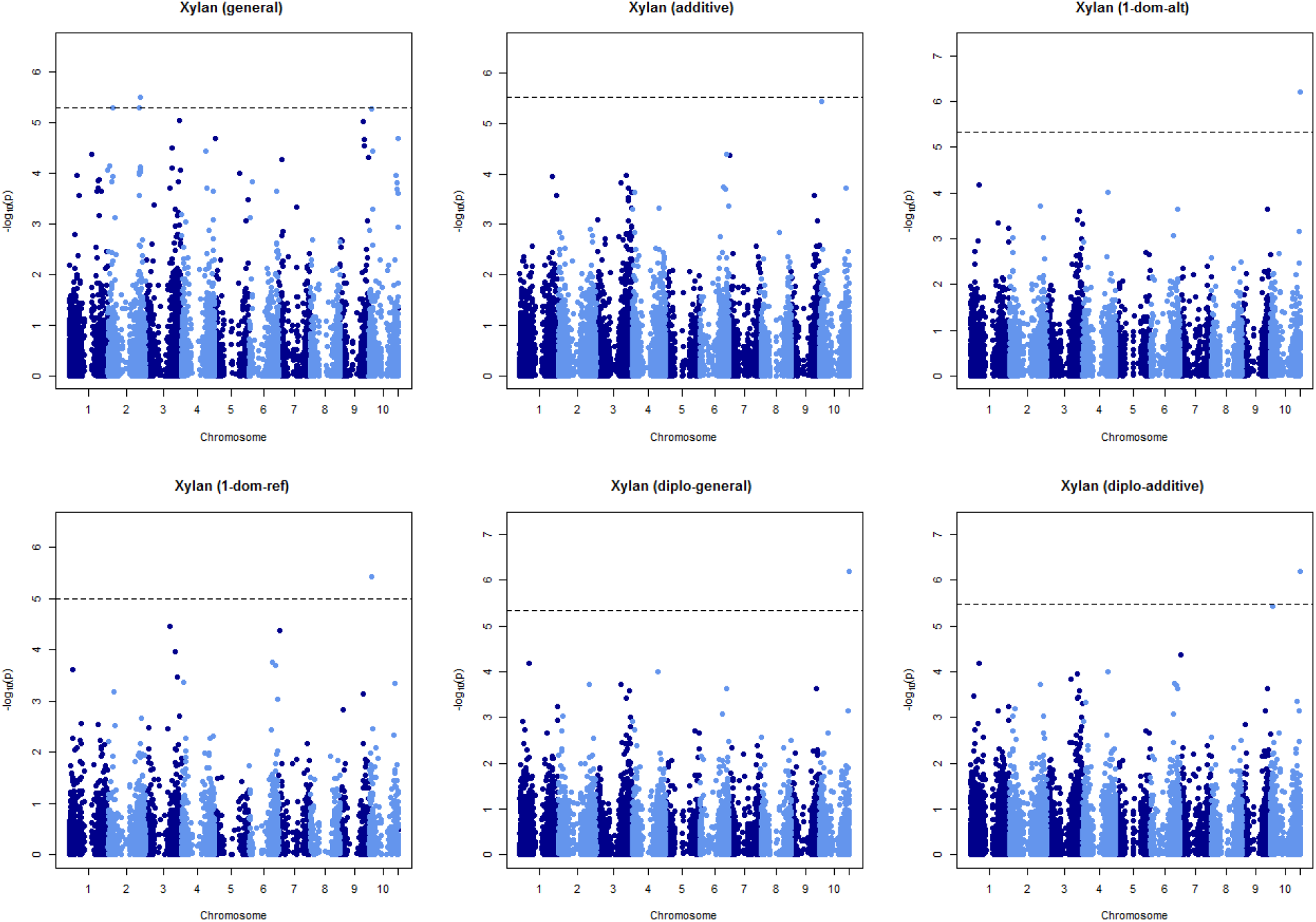
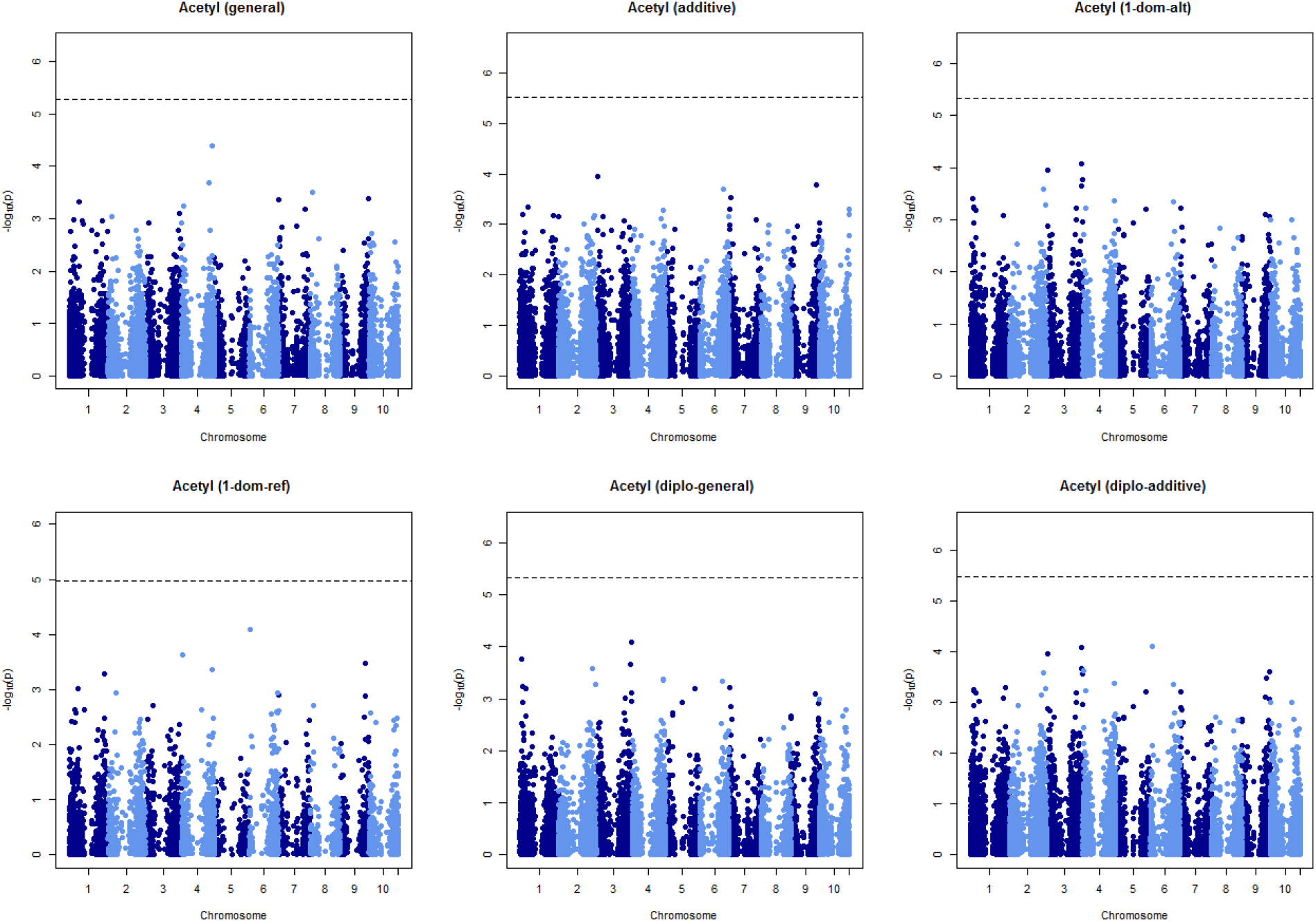
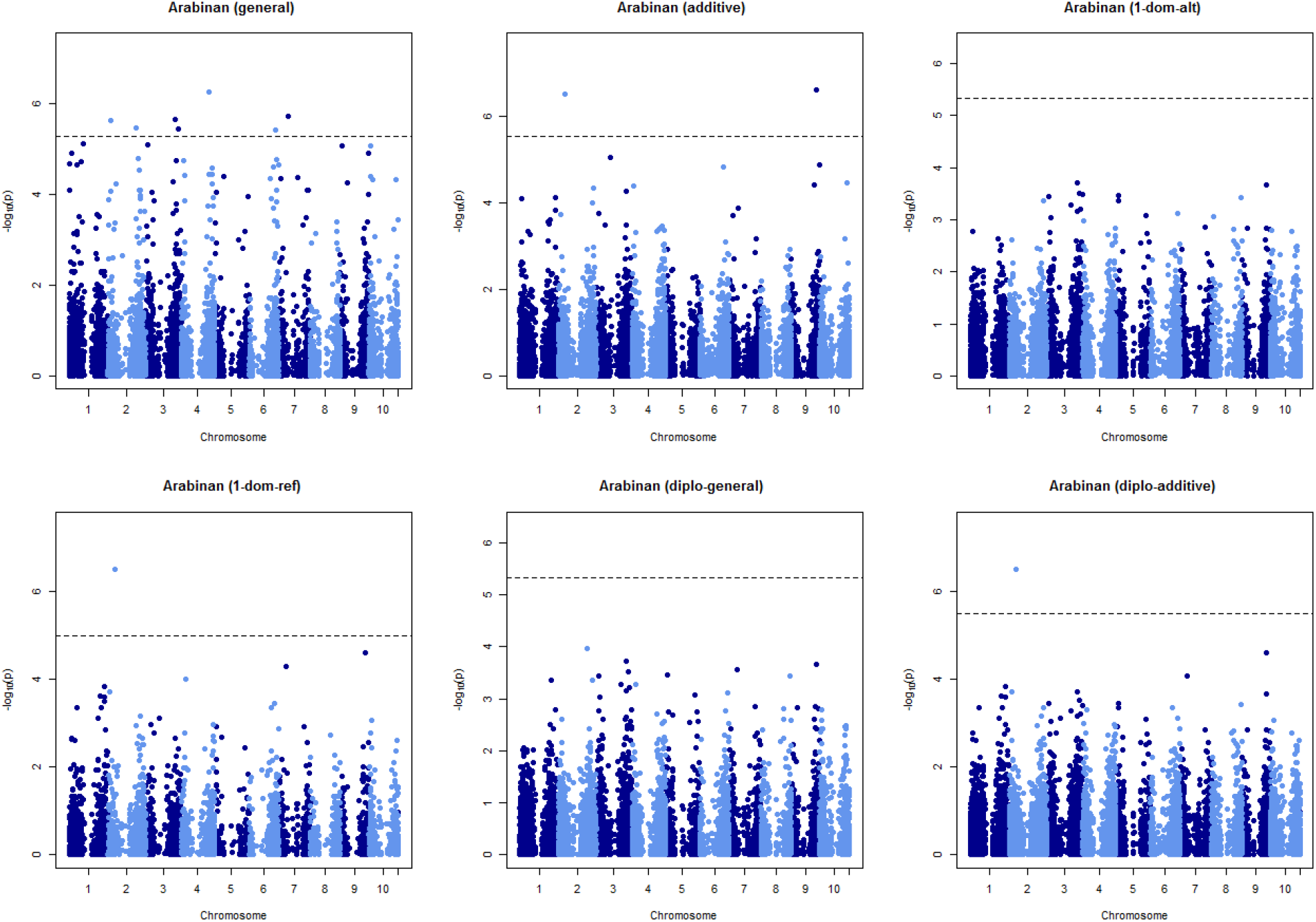
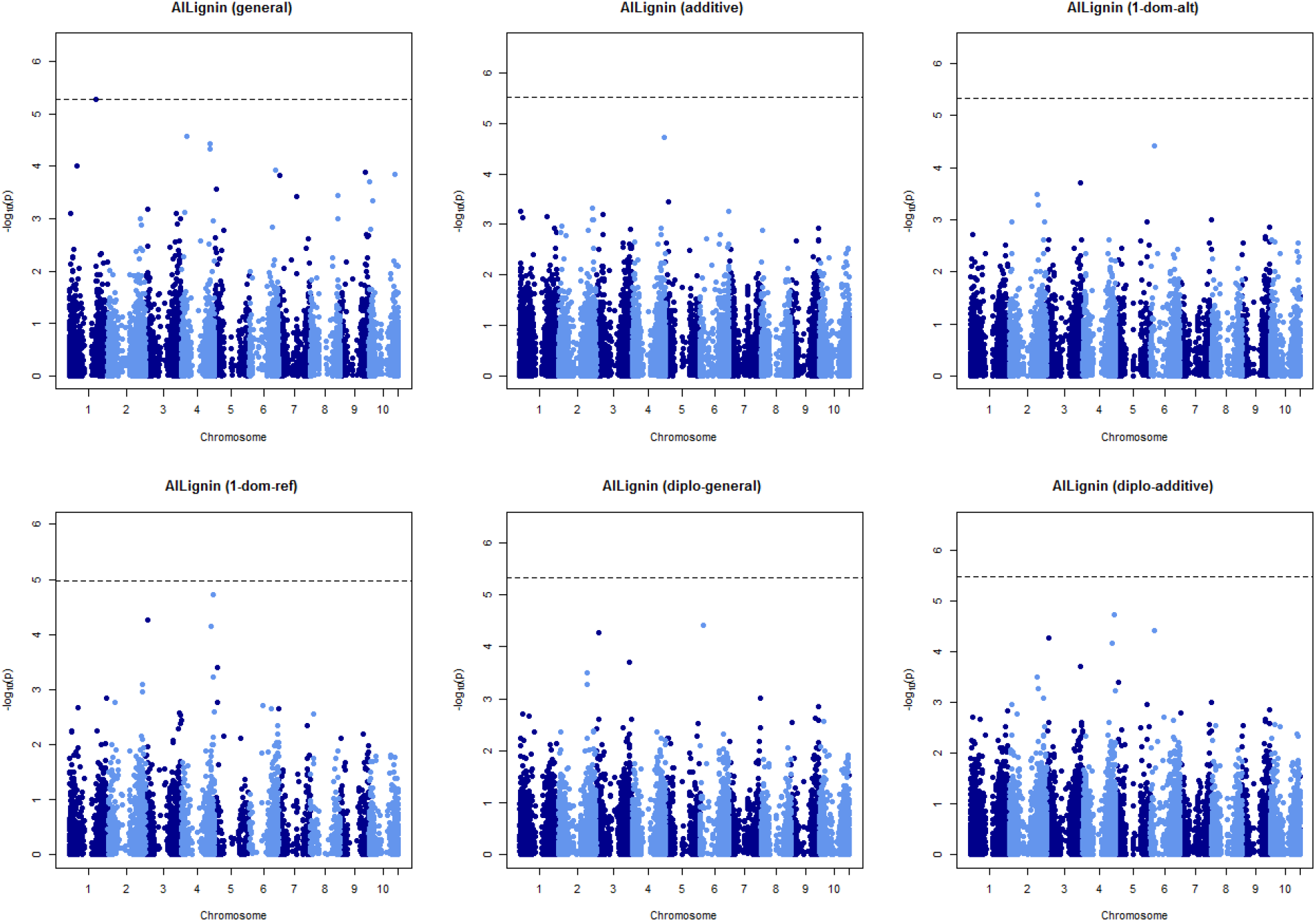
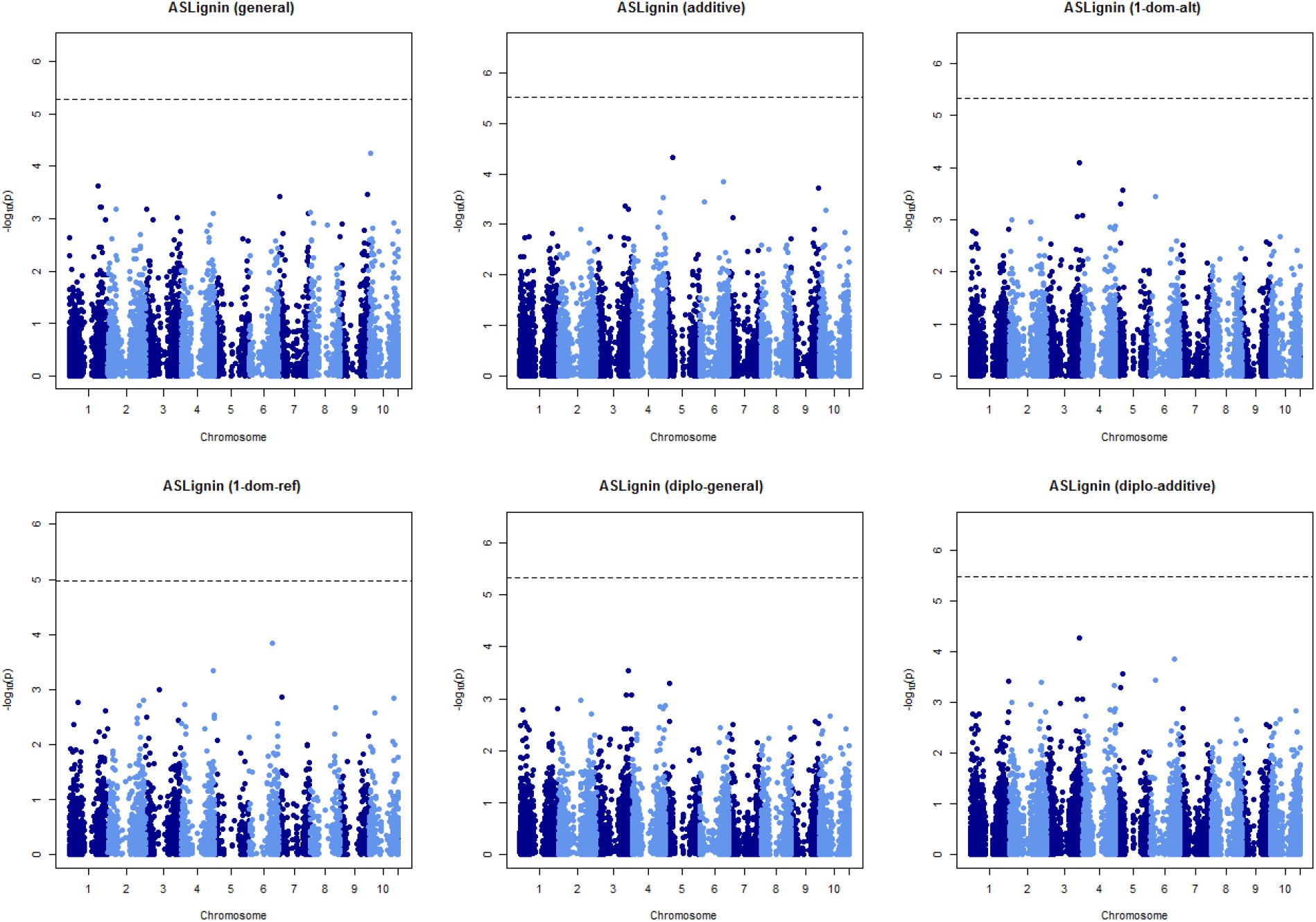
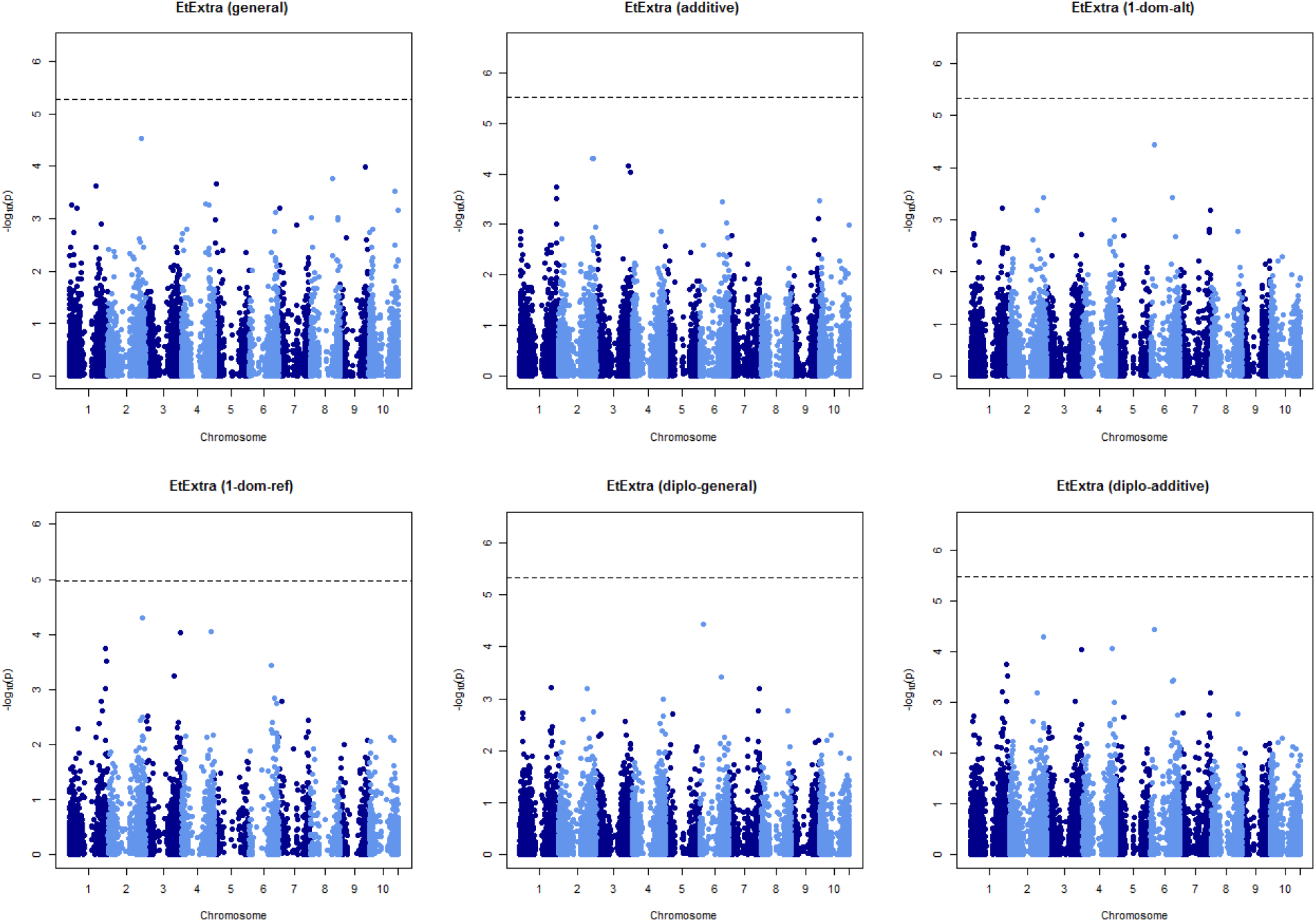
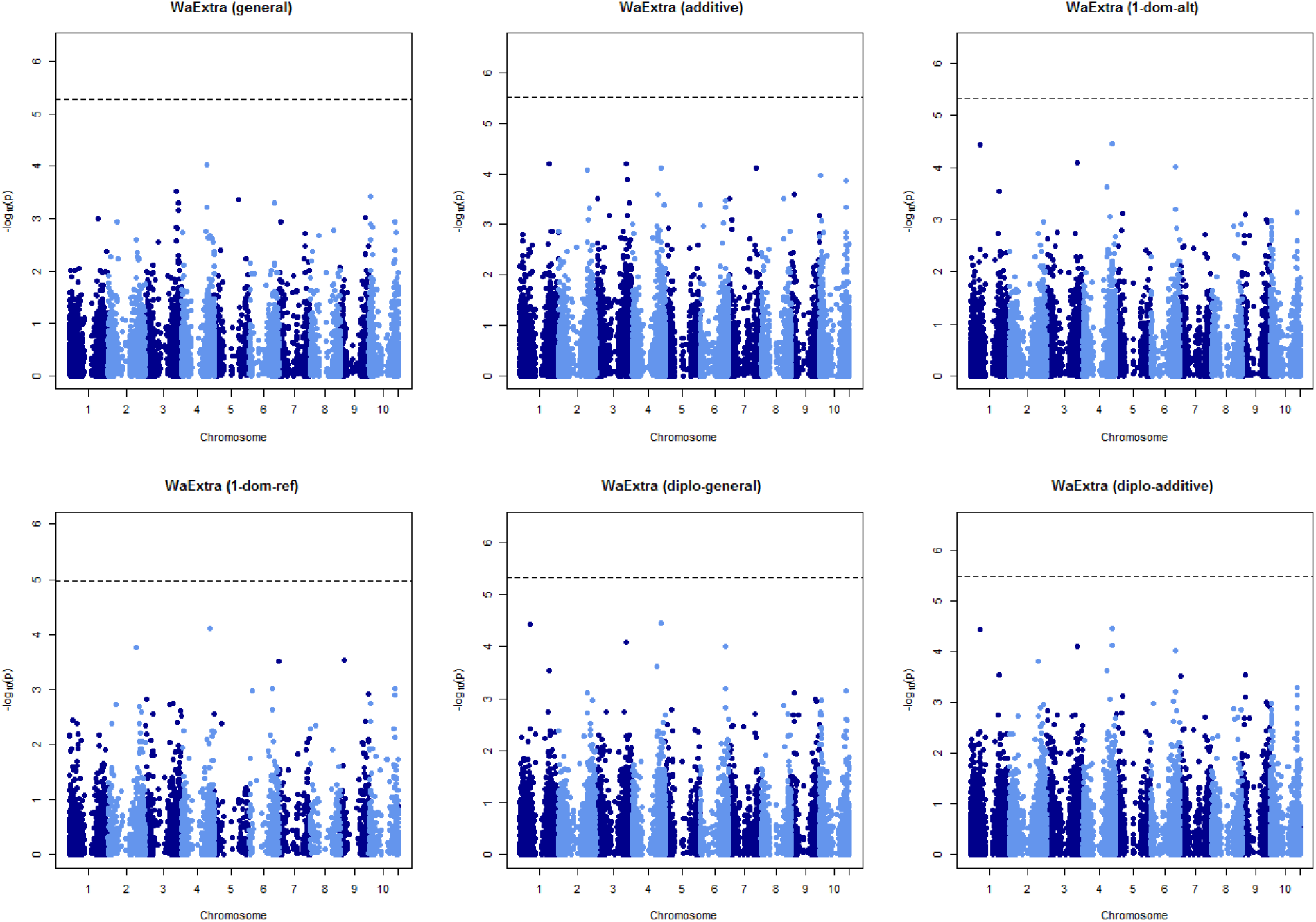
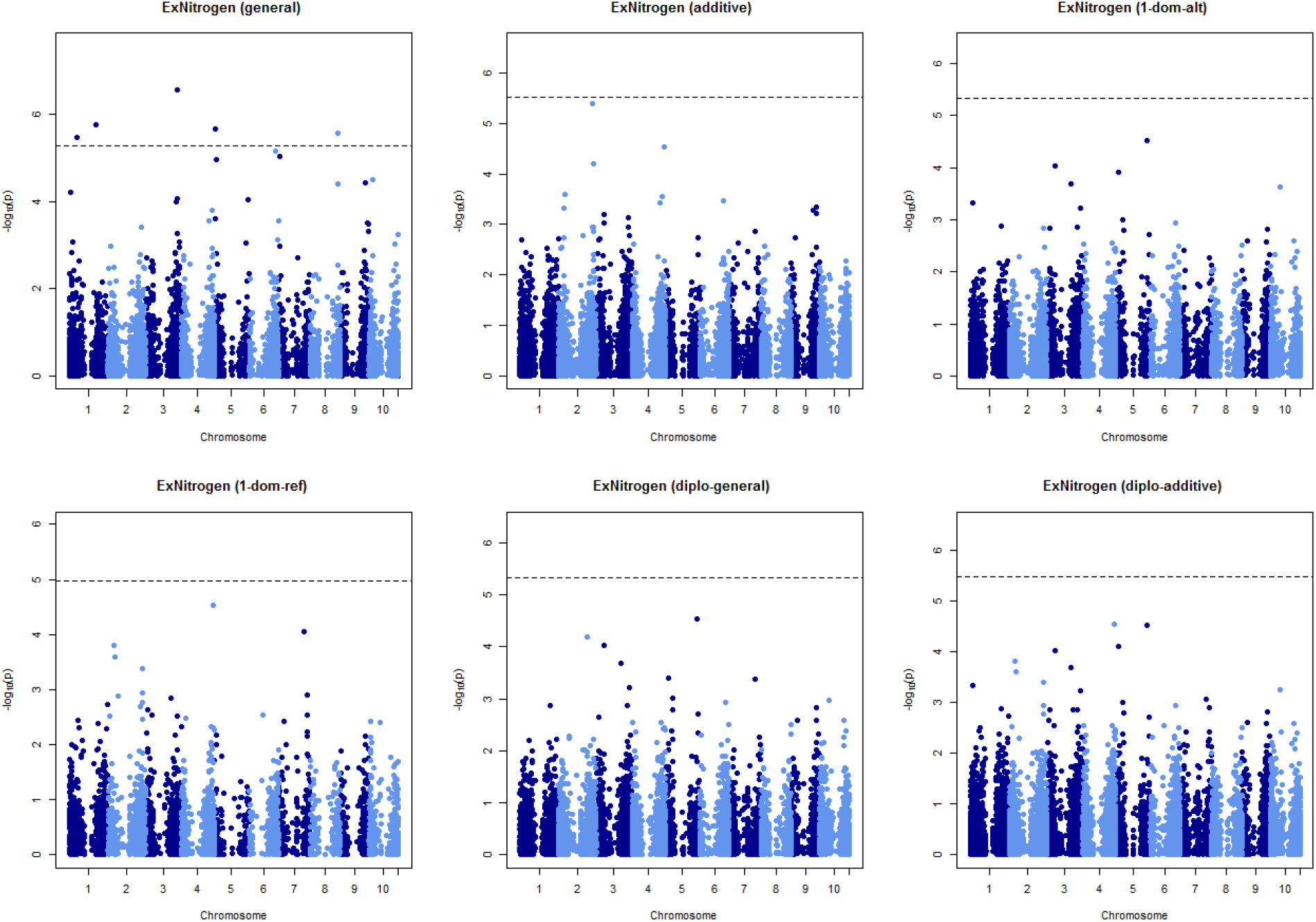
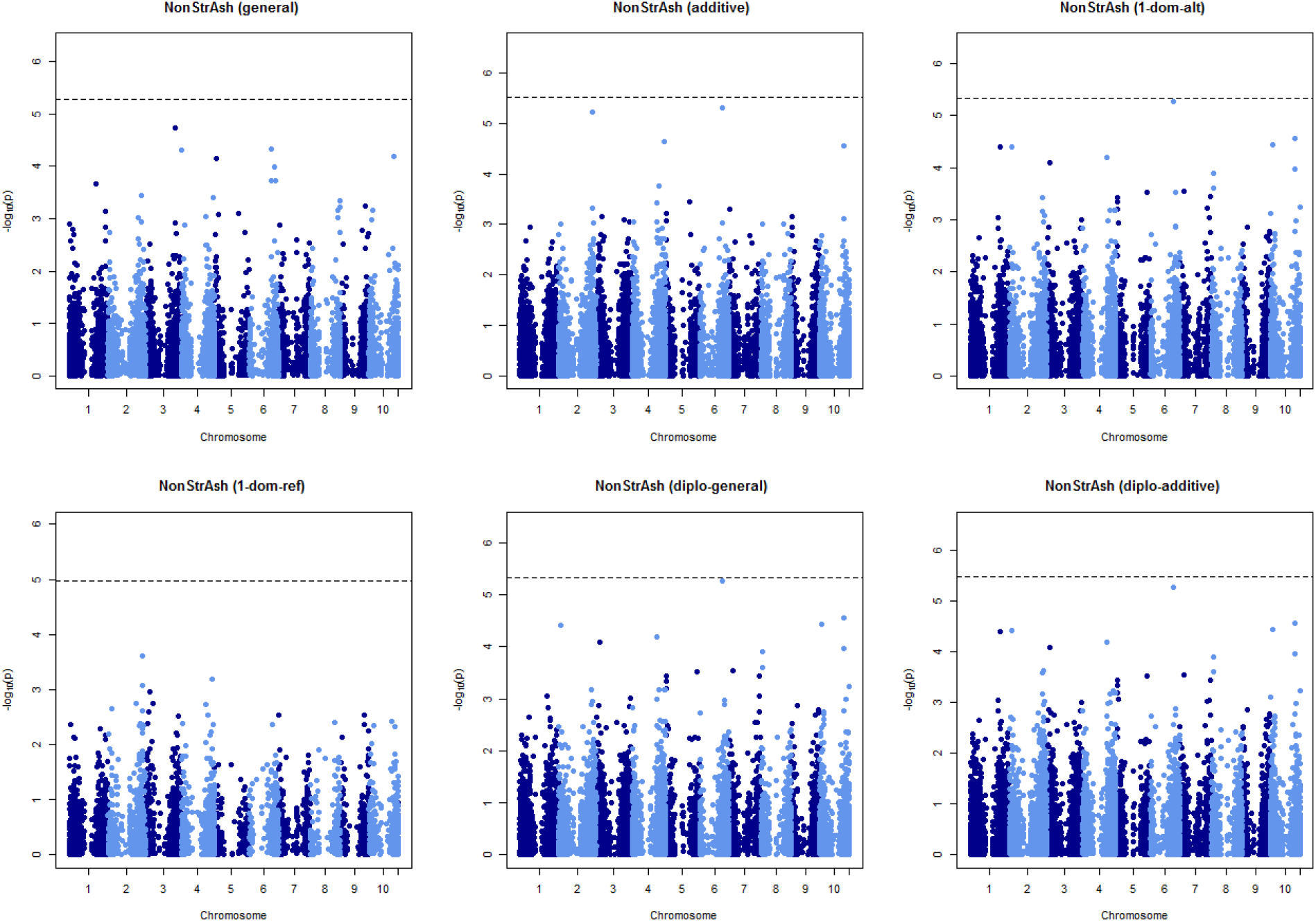
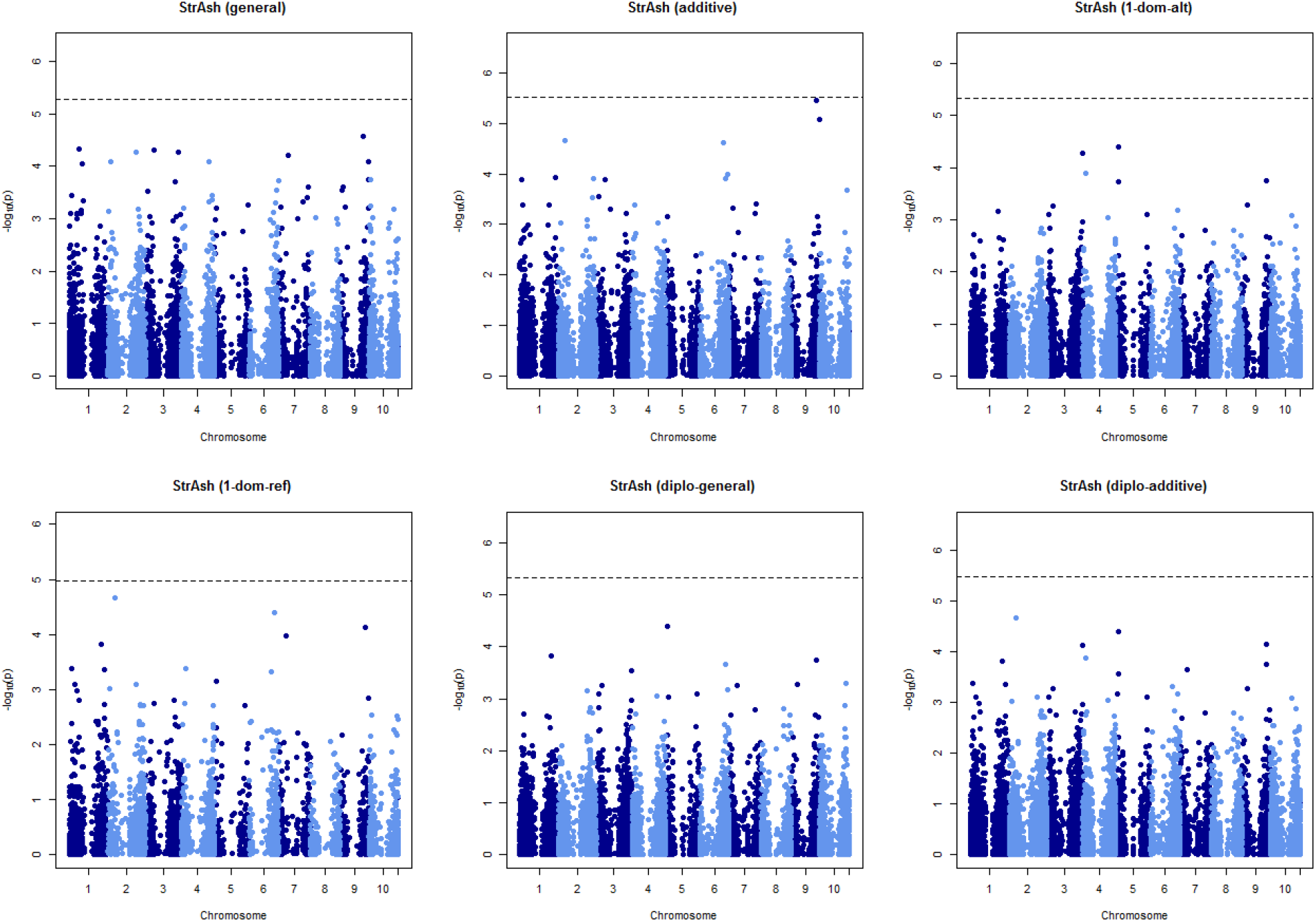
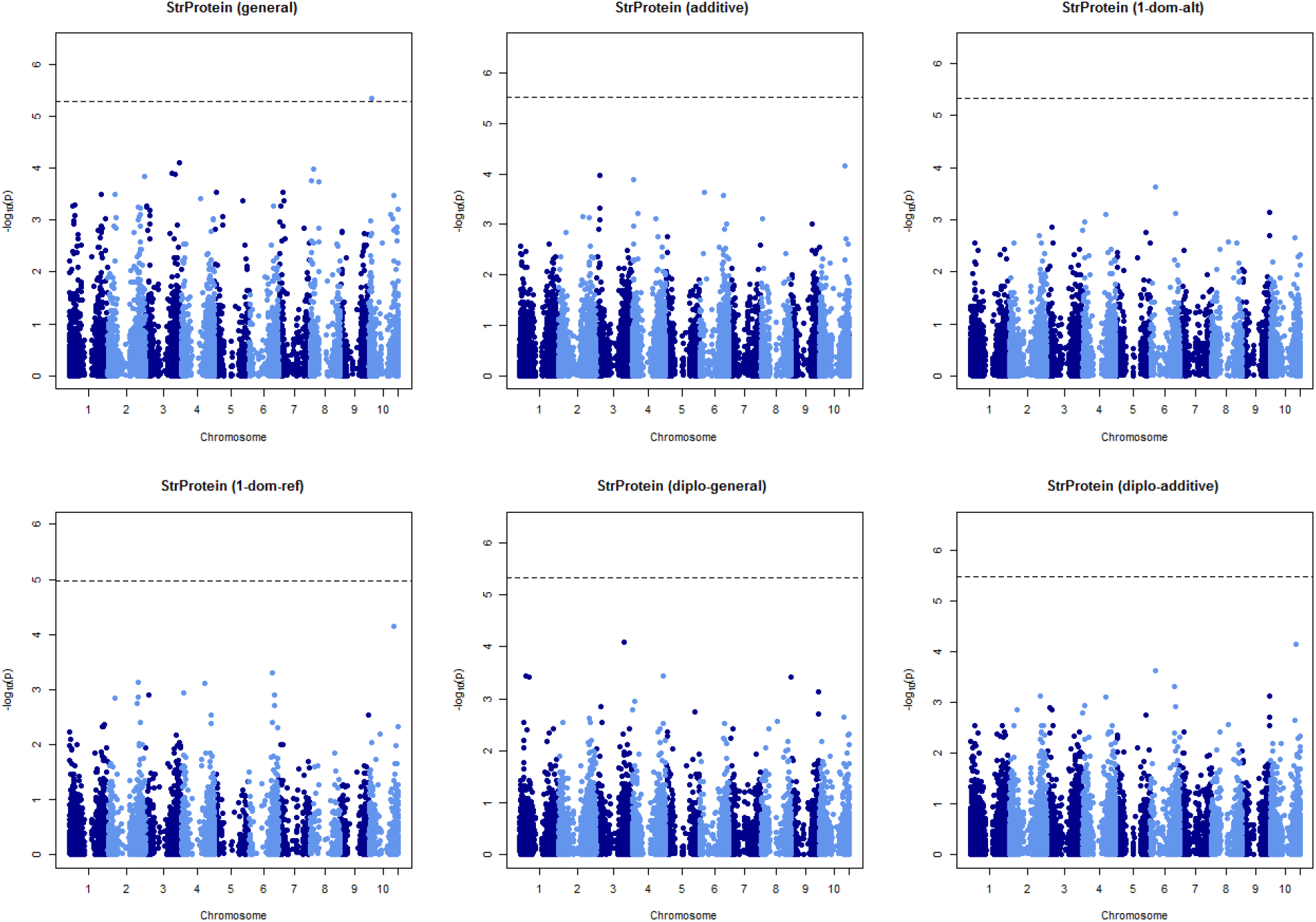
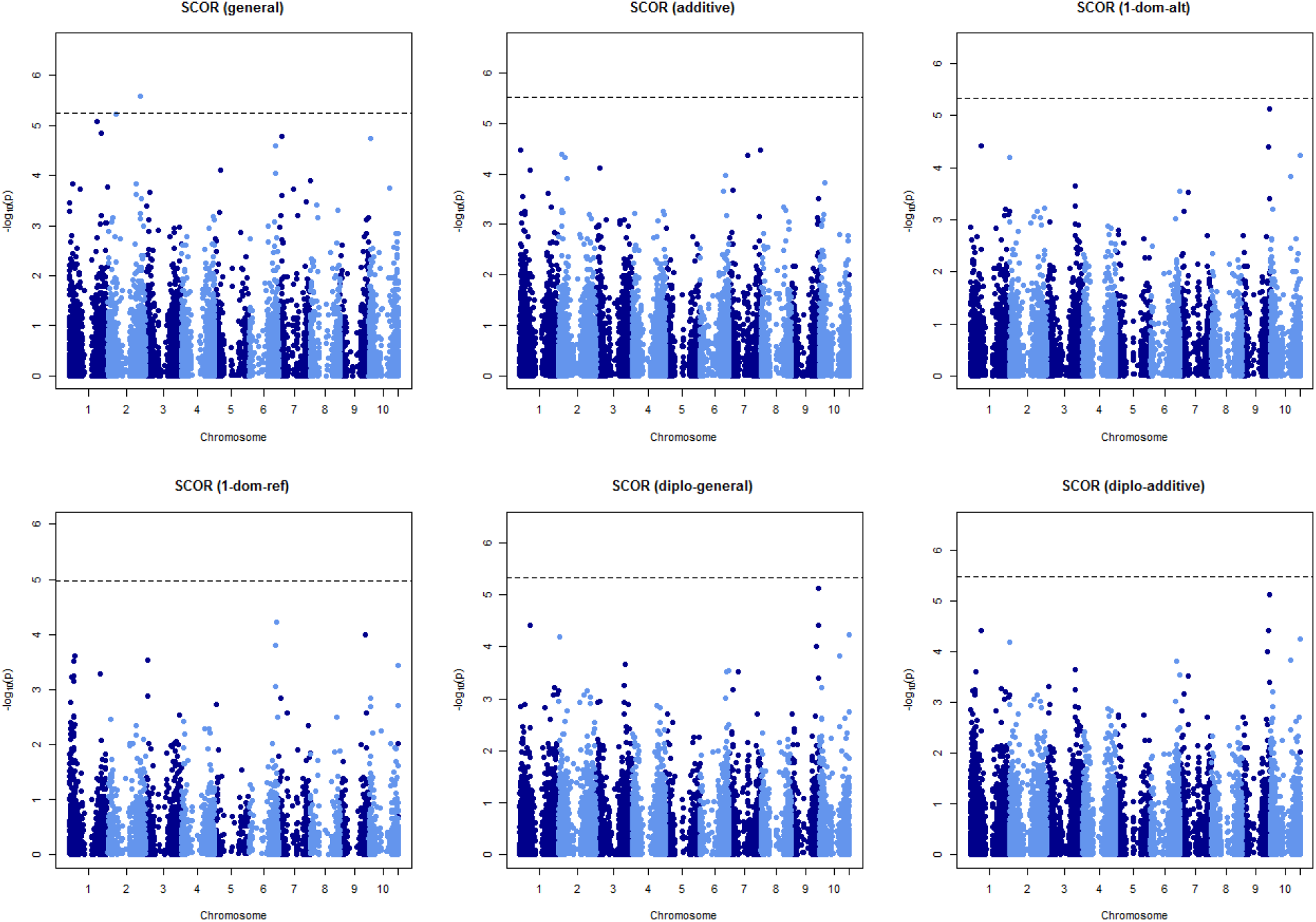
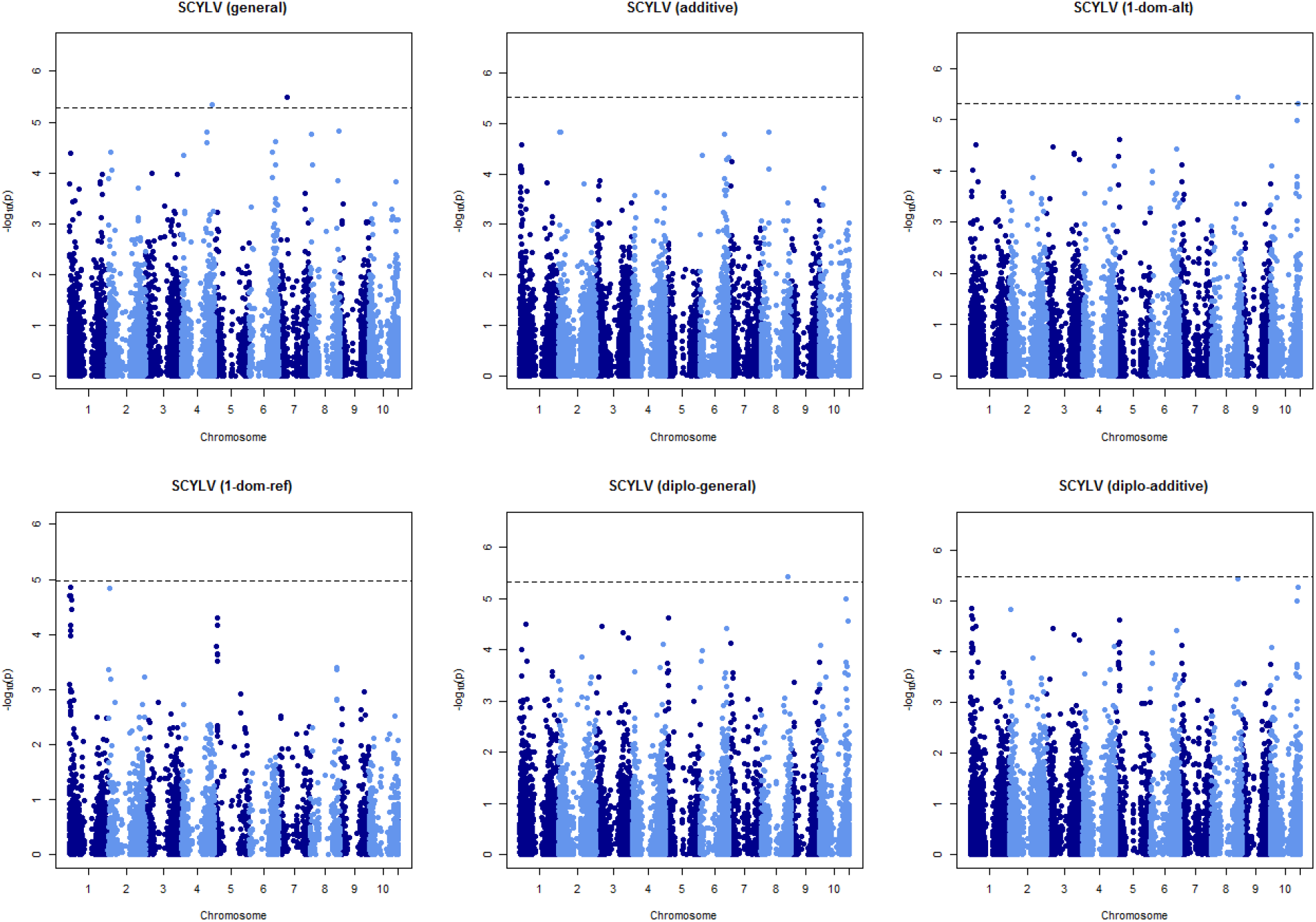
Significant marker-trait associations identified in the sugarcane diversity panel with 162 accessions with ploidy = 8. Stalk = Stalk number; Diameter = Stalk diameter; Width = Leaf width; Length = Leaf length; Weight = Total weight; Dry = Dry weight; Water = Water content; Internode = Internode length.

**Table S1.**
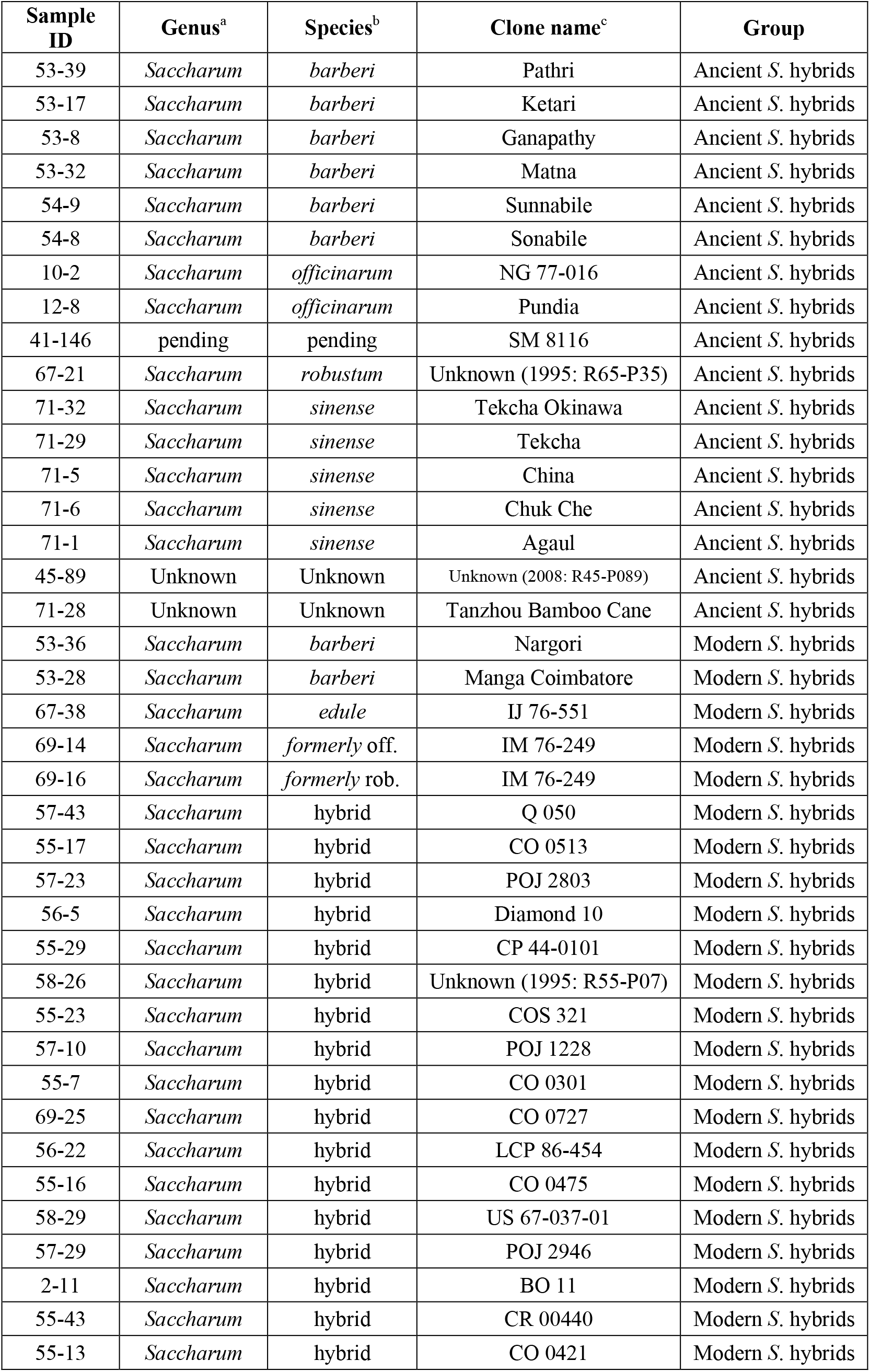
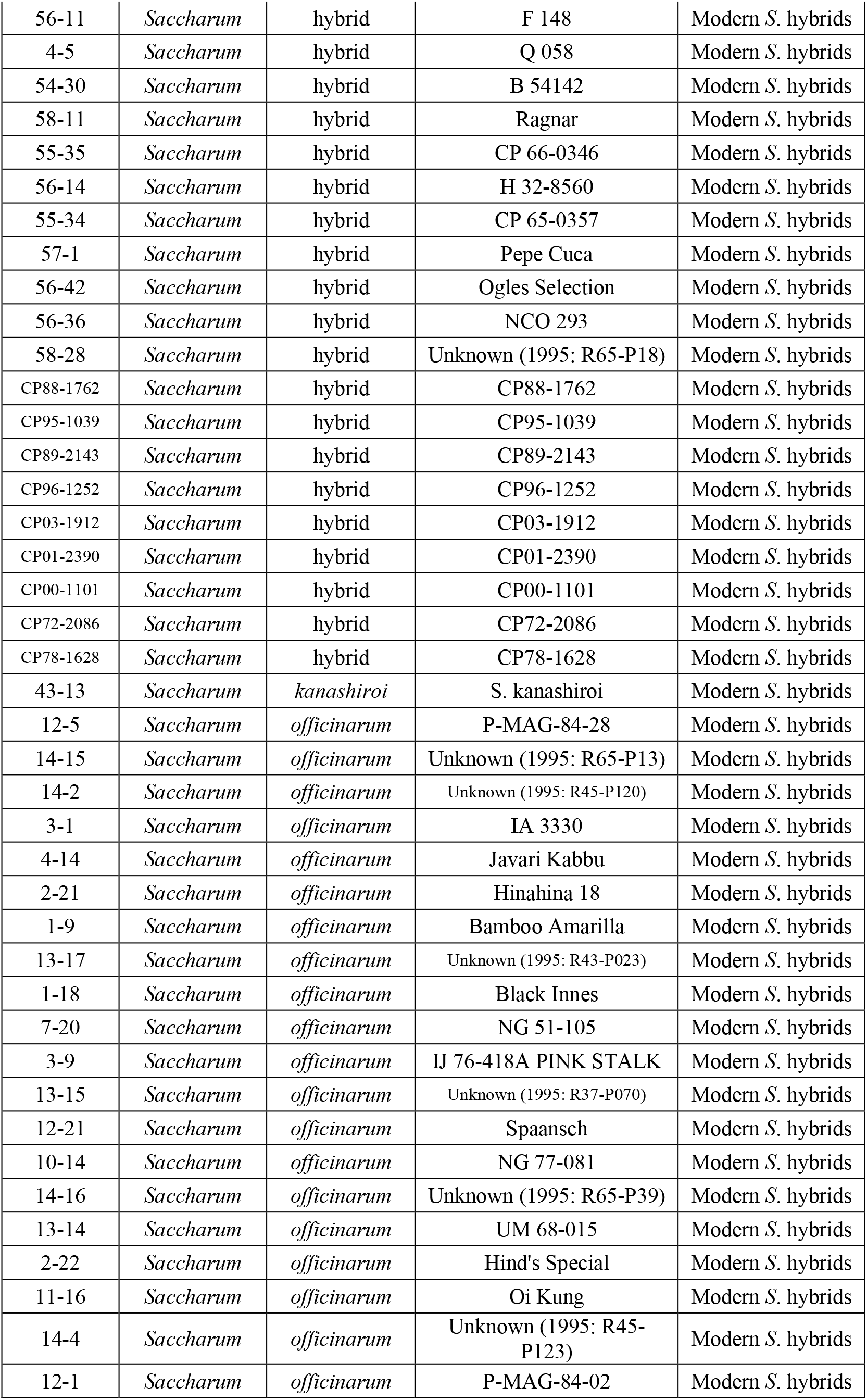
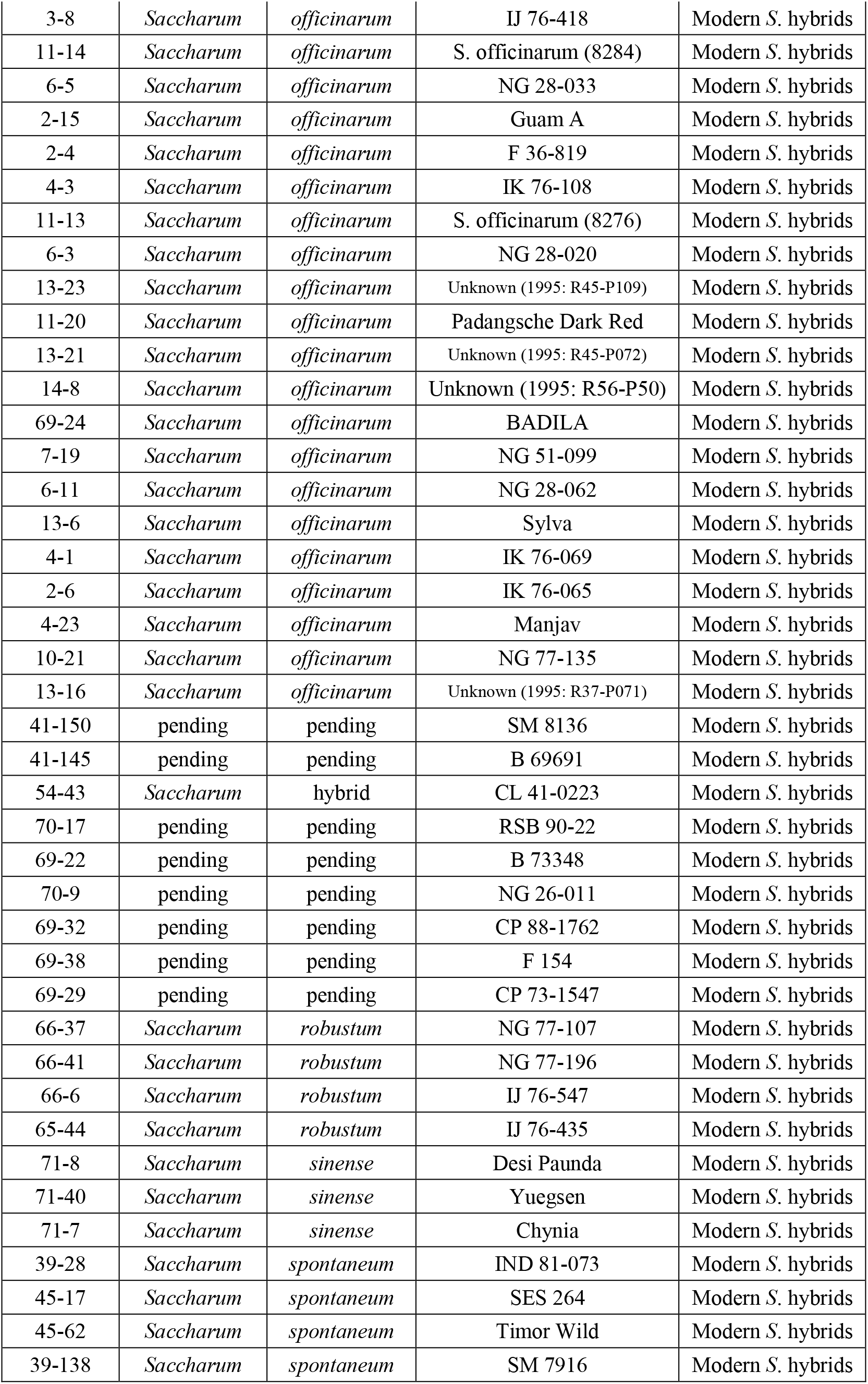
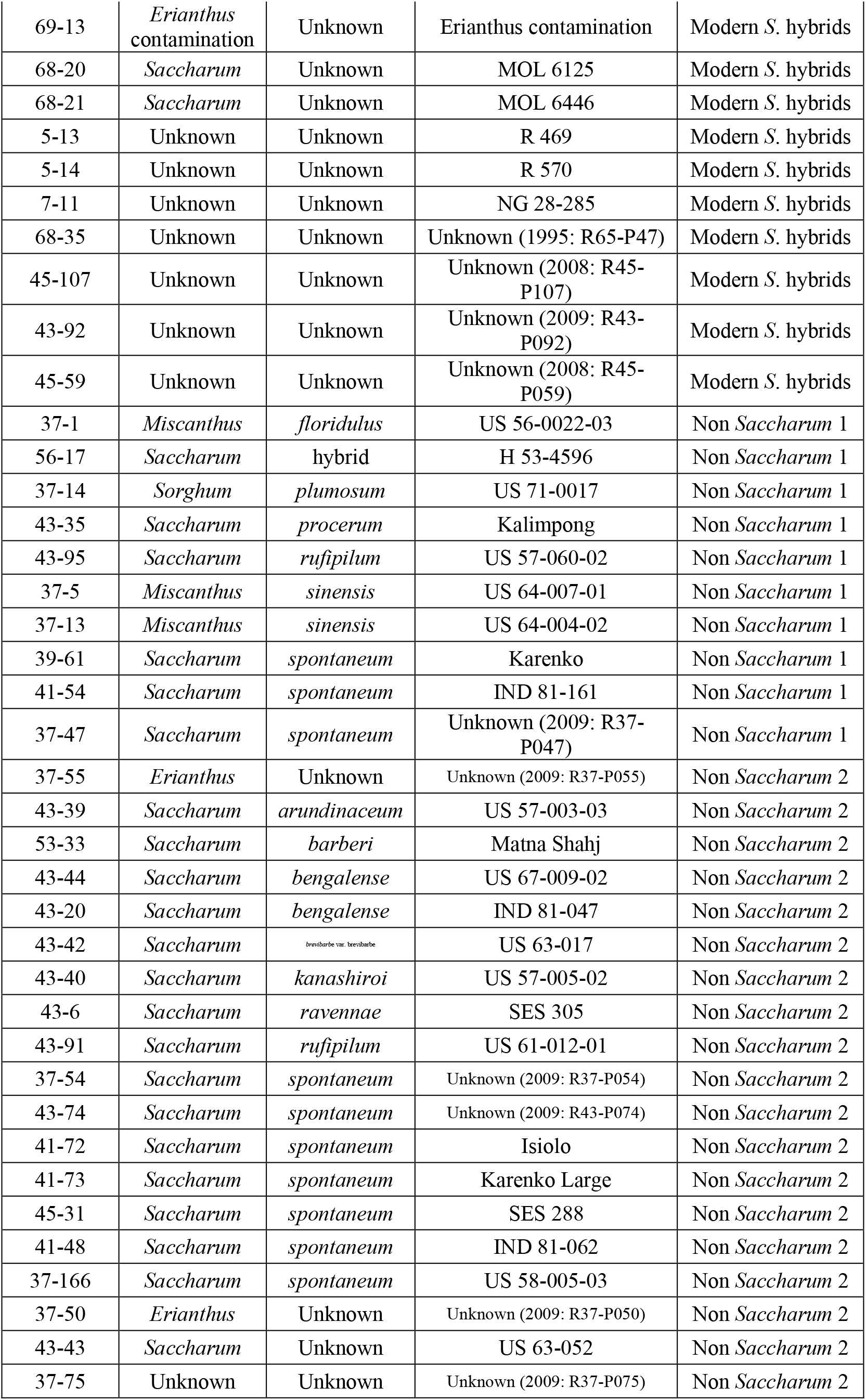
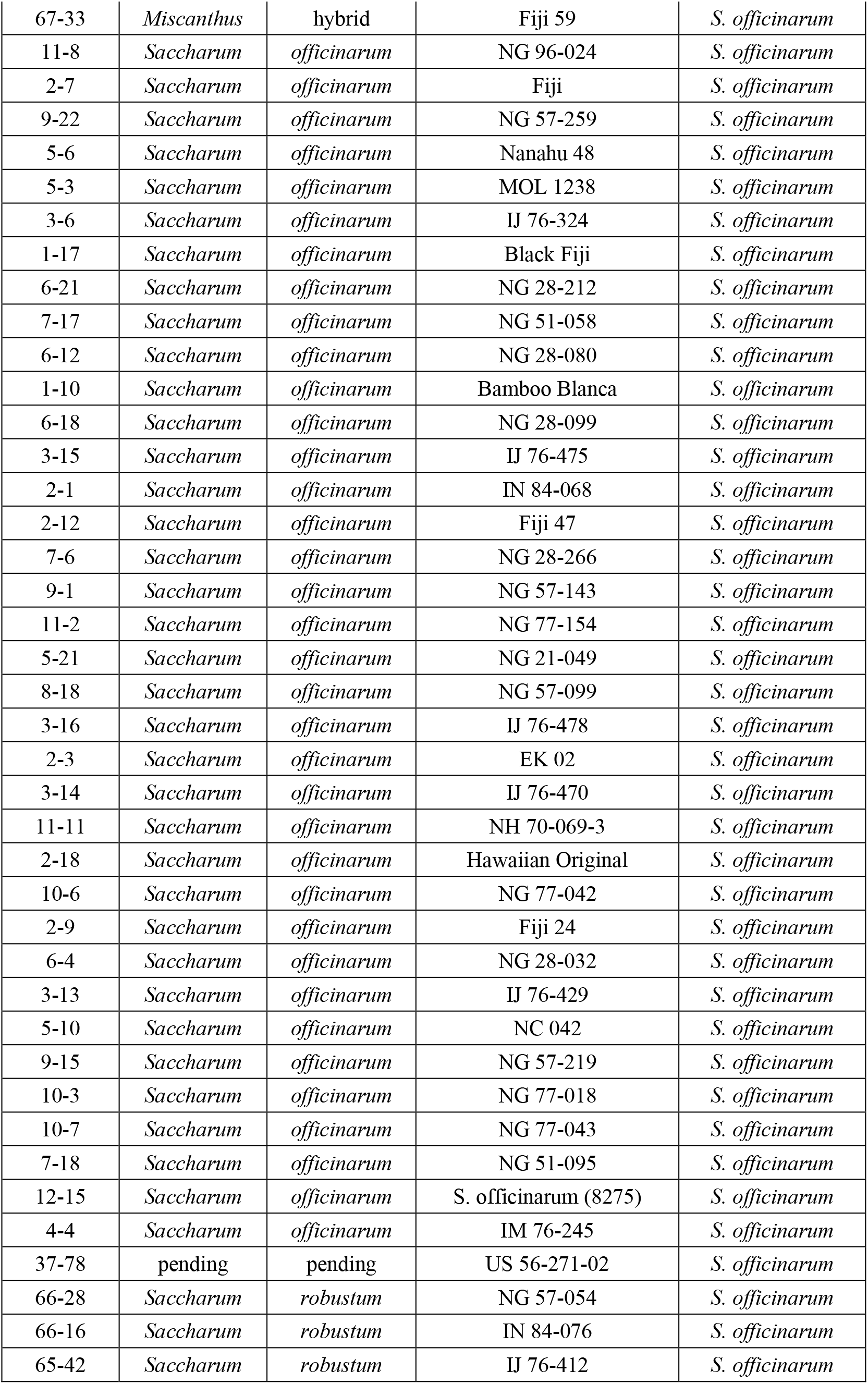
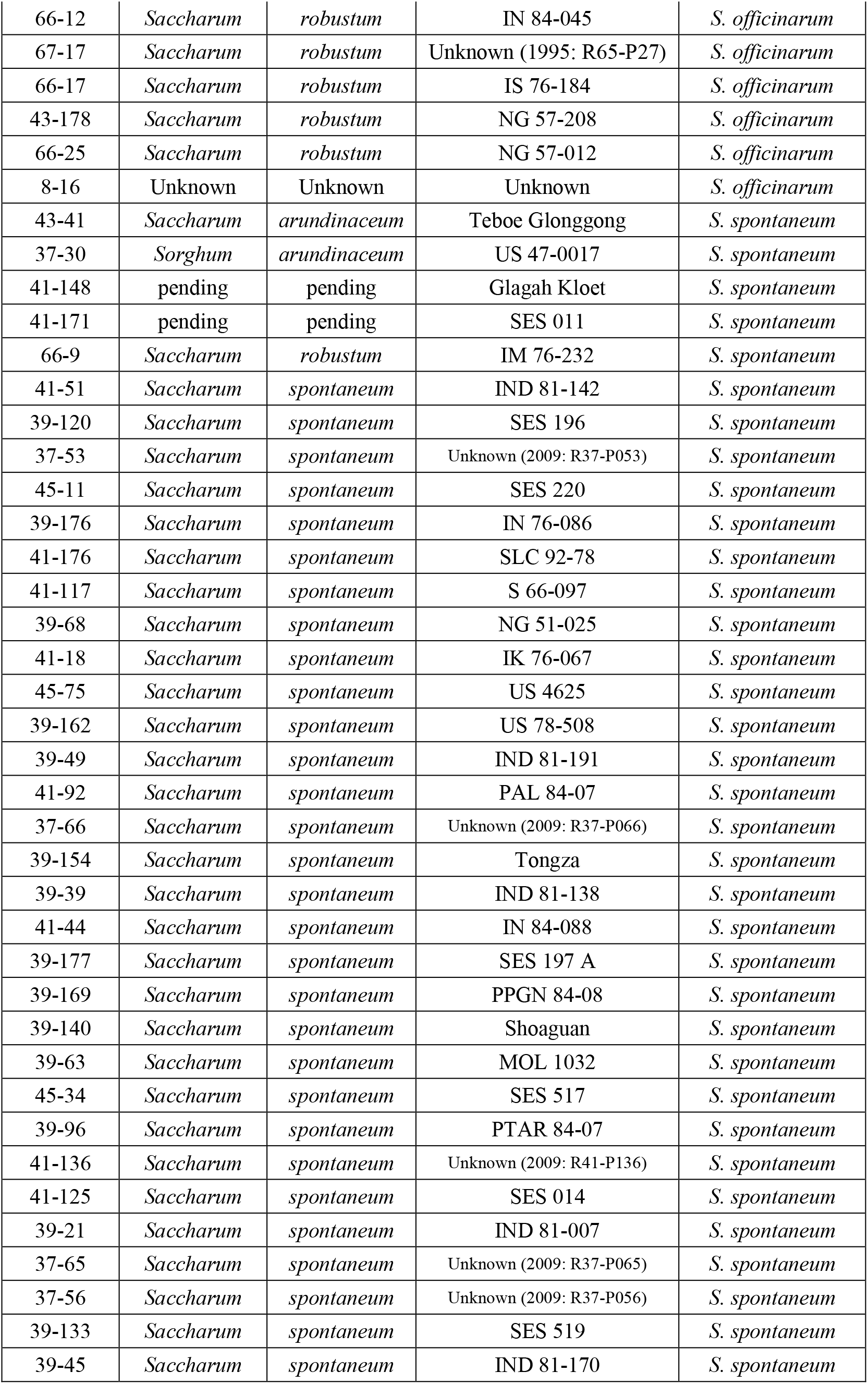
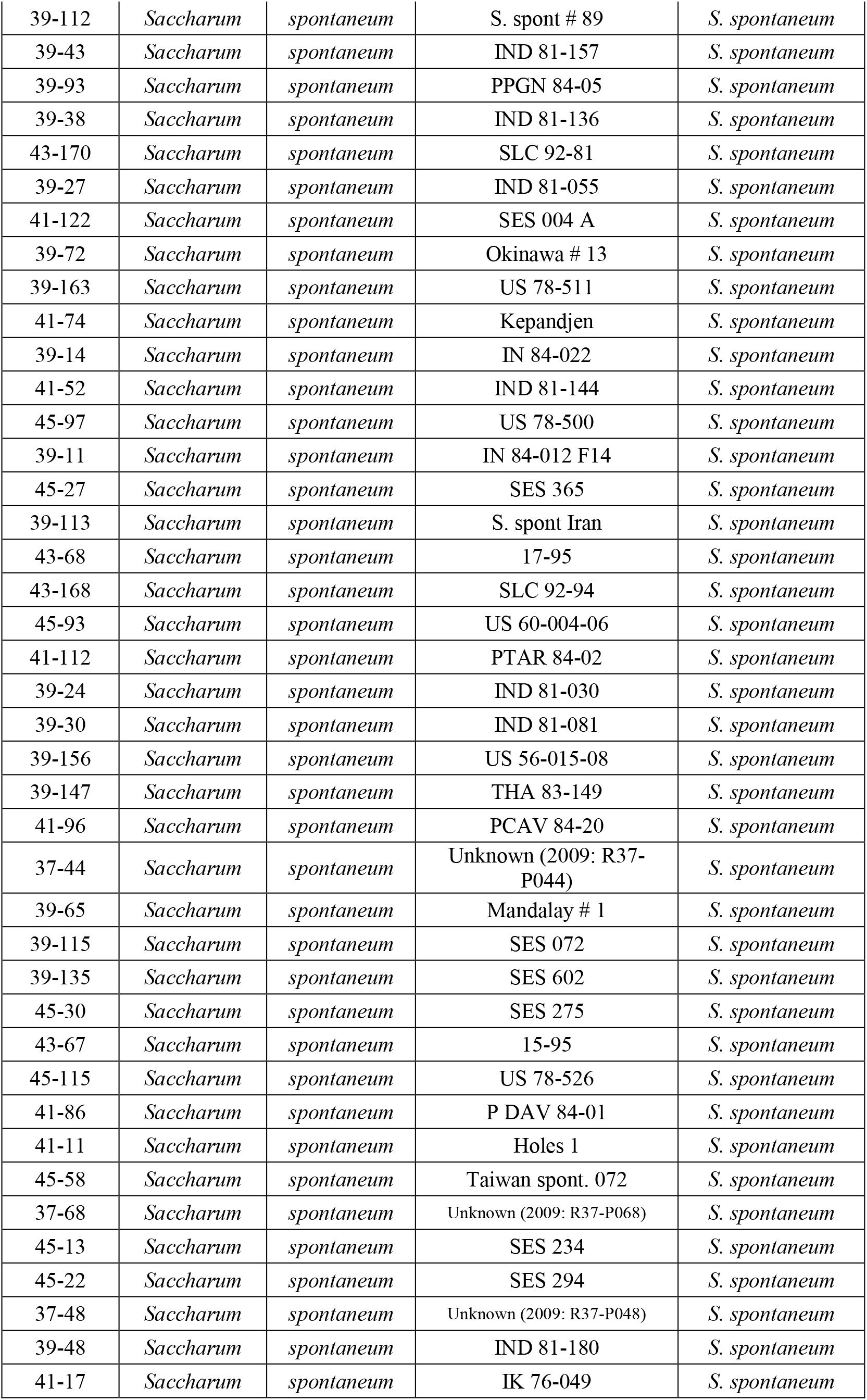
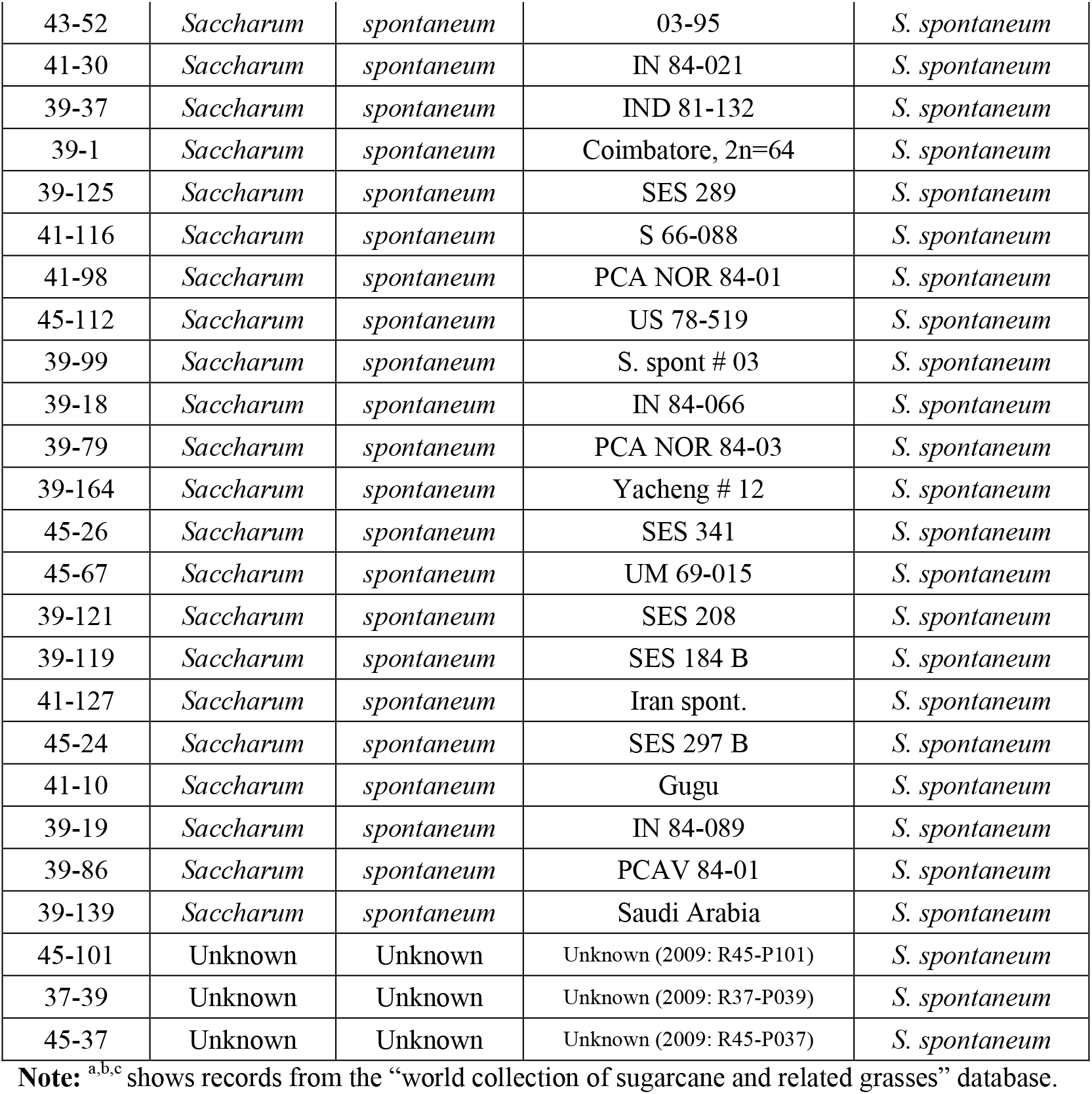
Details of genotypes included in the sugarcane diversity panel.

**Table S2.**
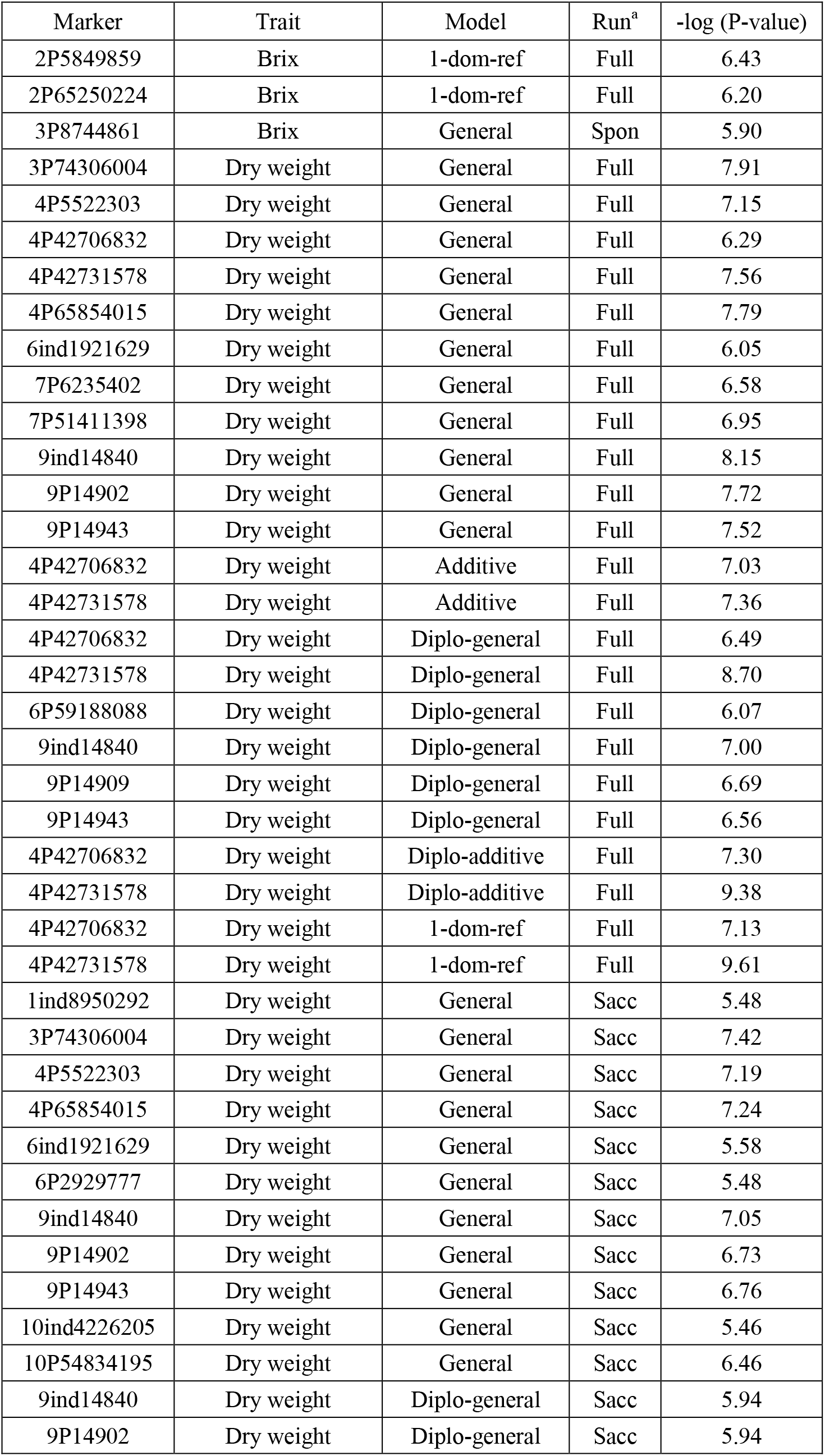
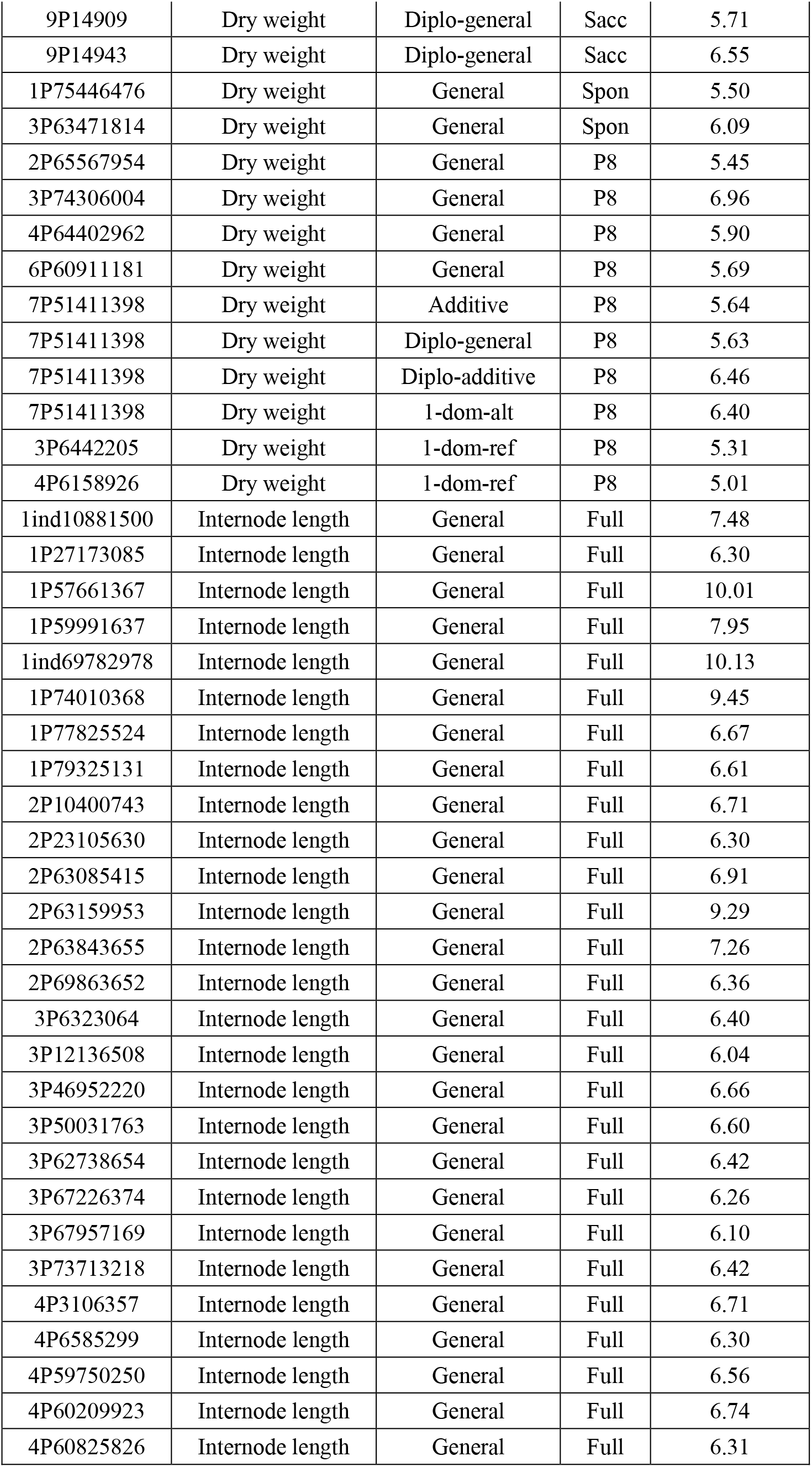
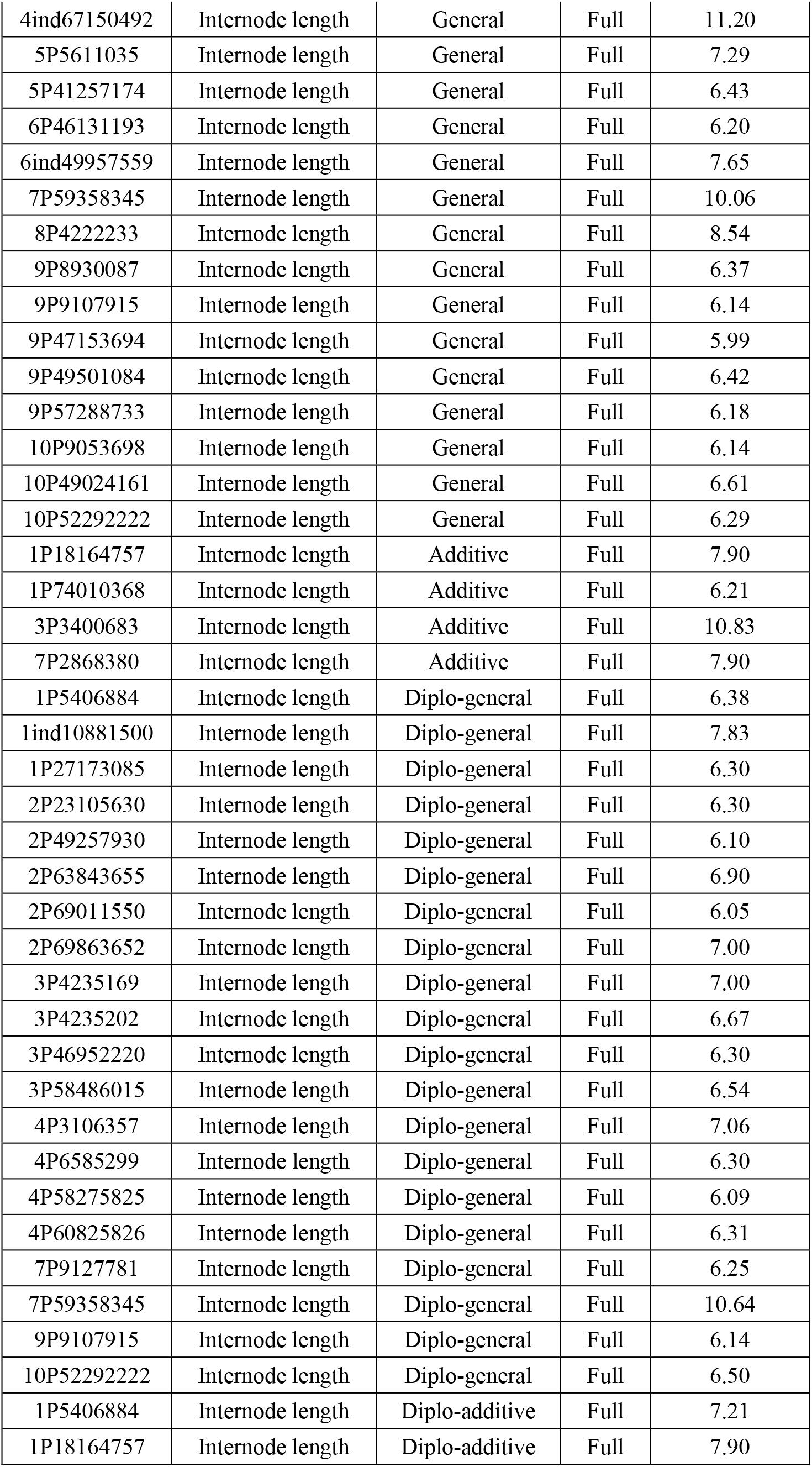
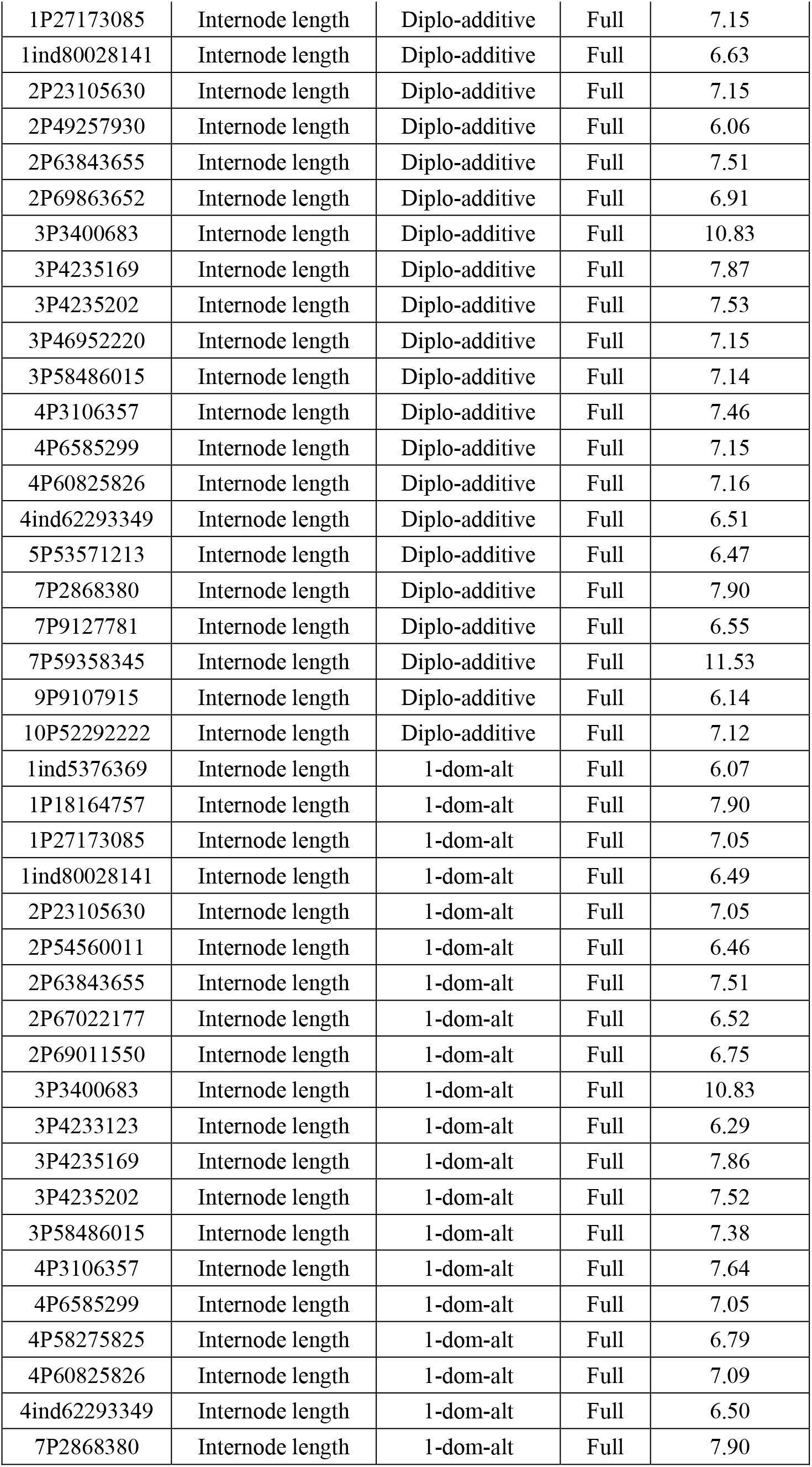
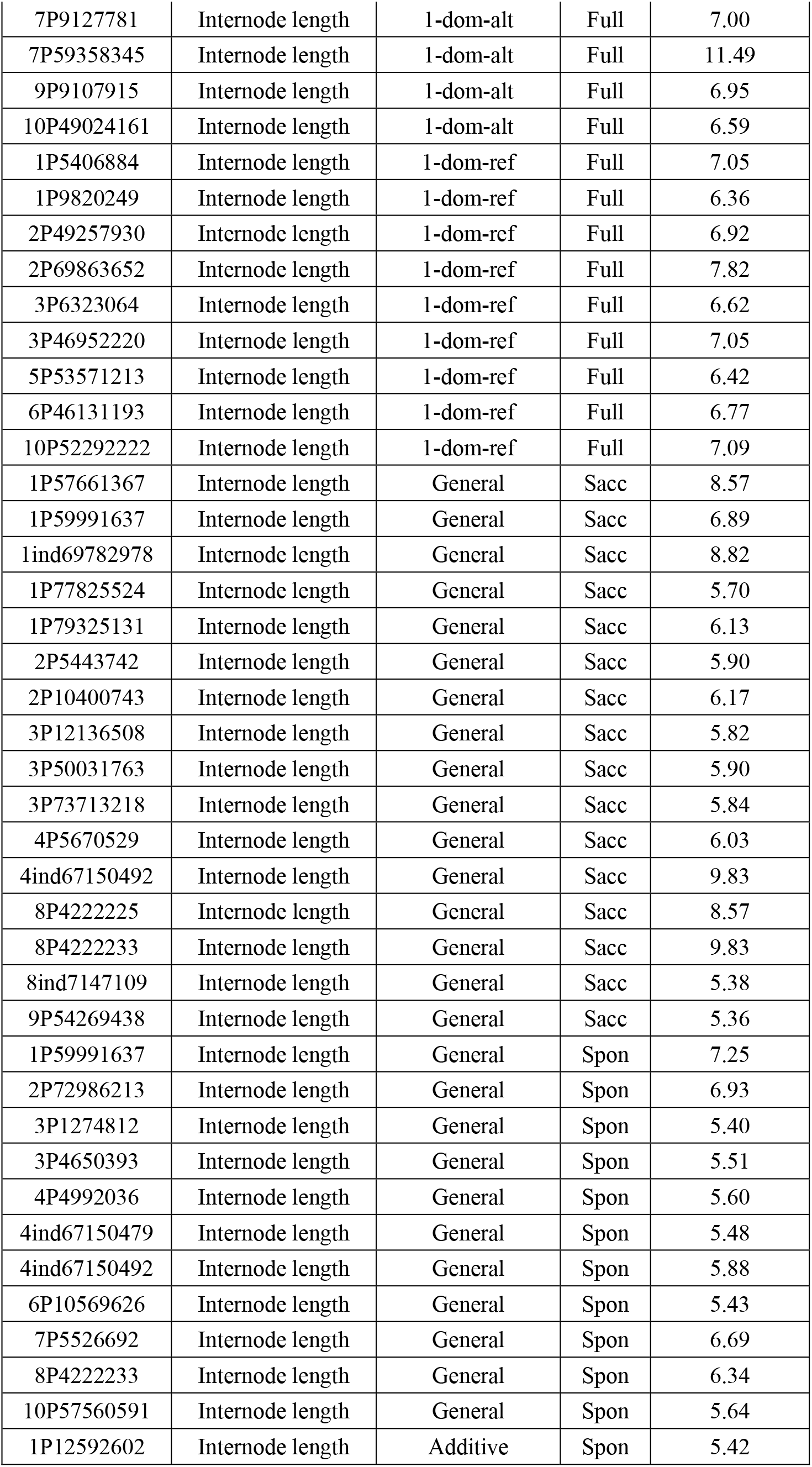
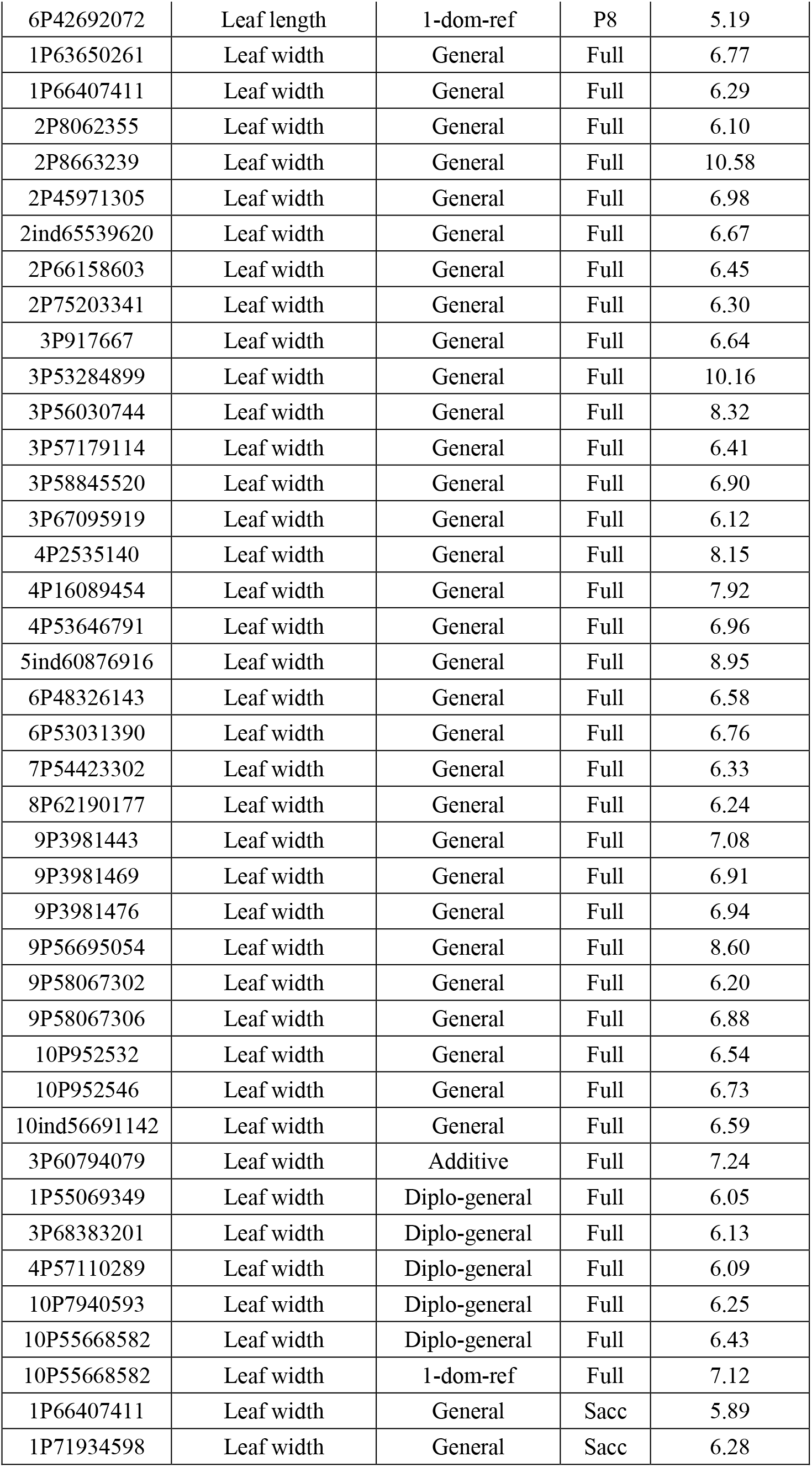
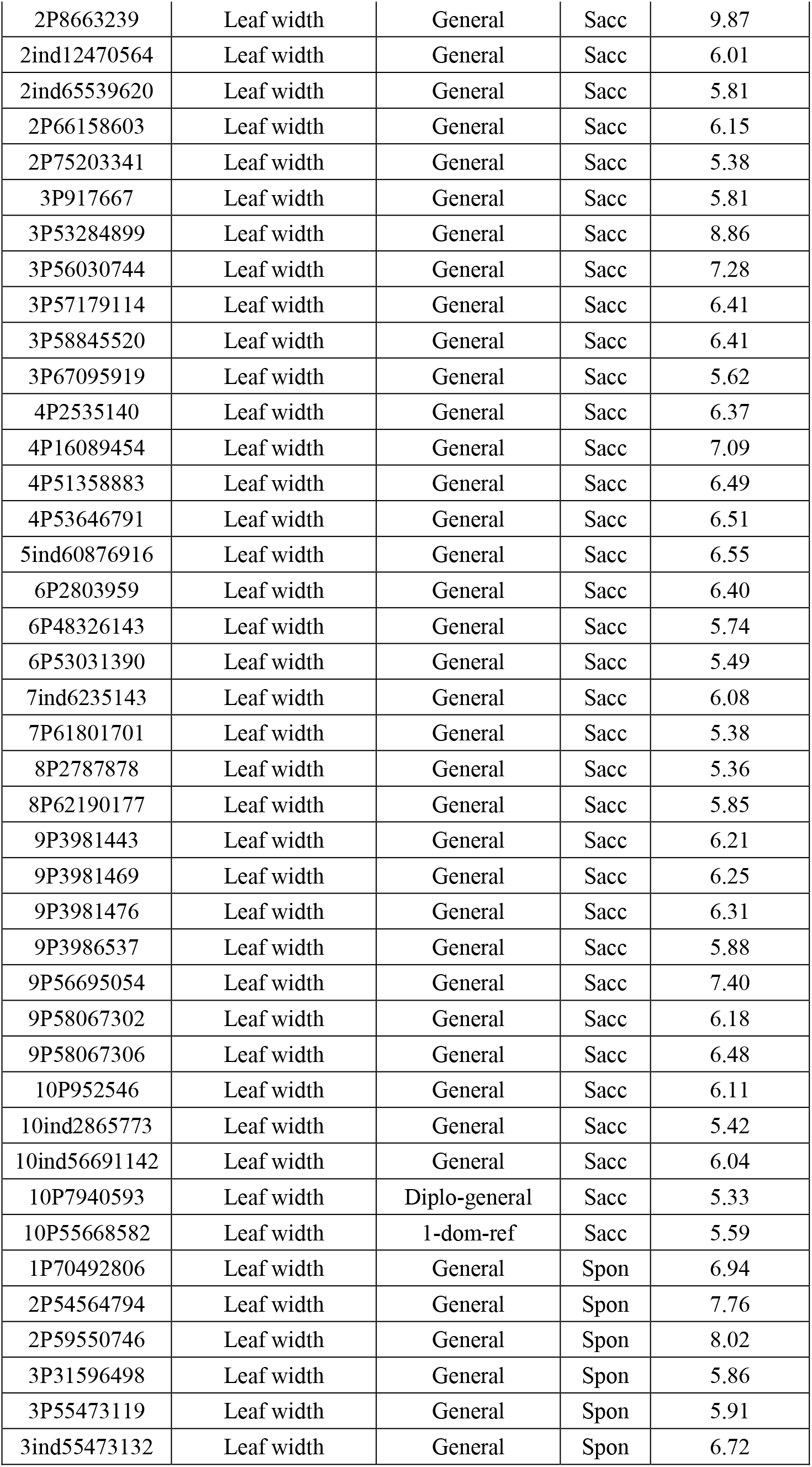
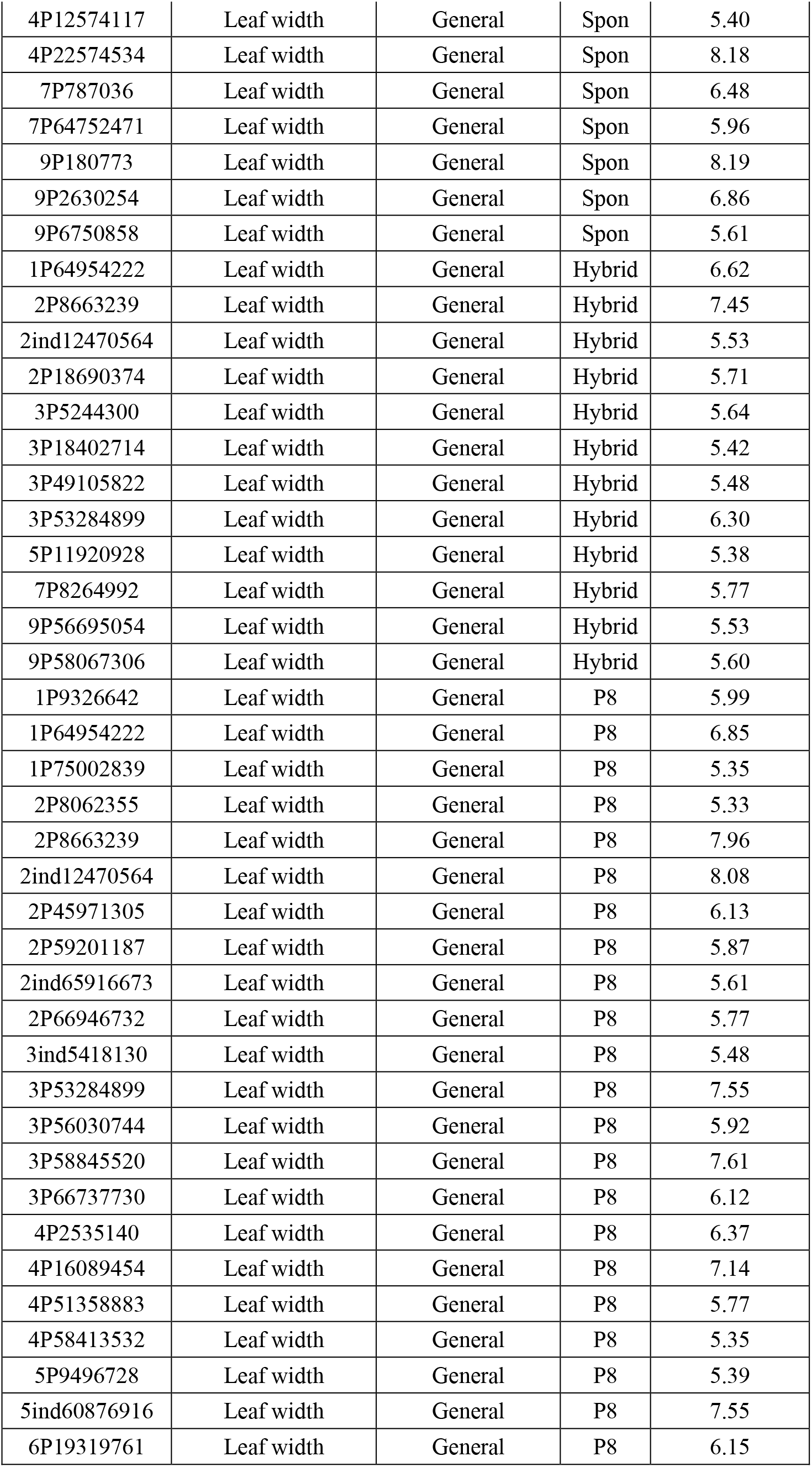
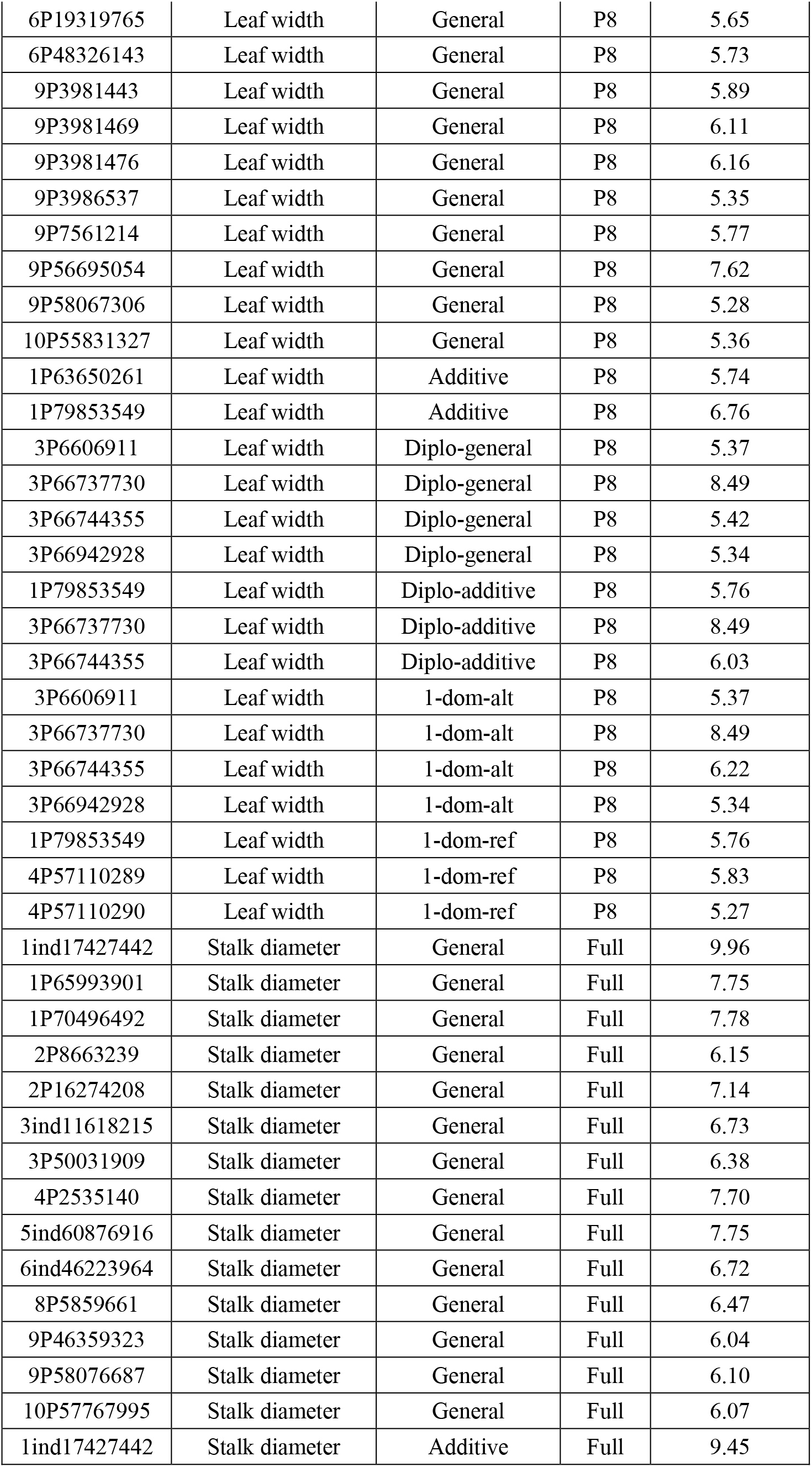
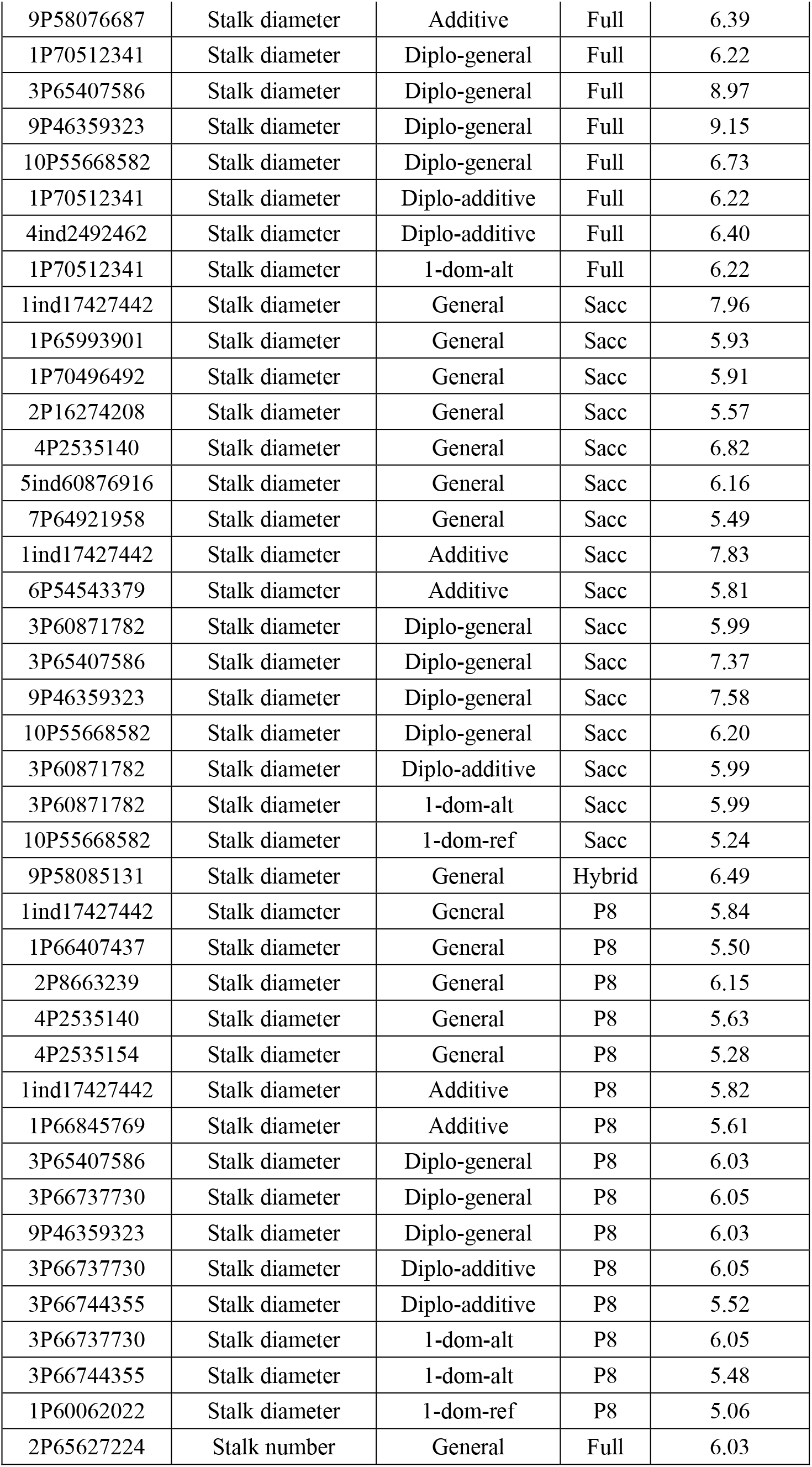
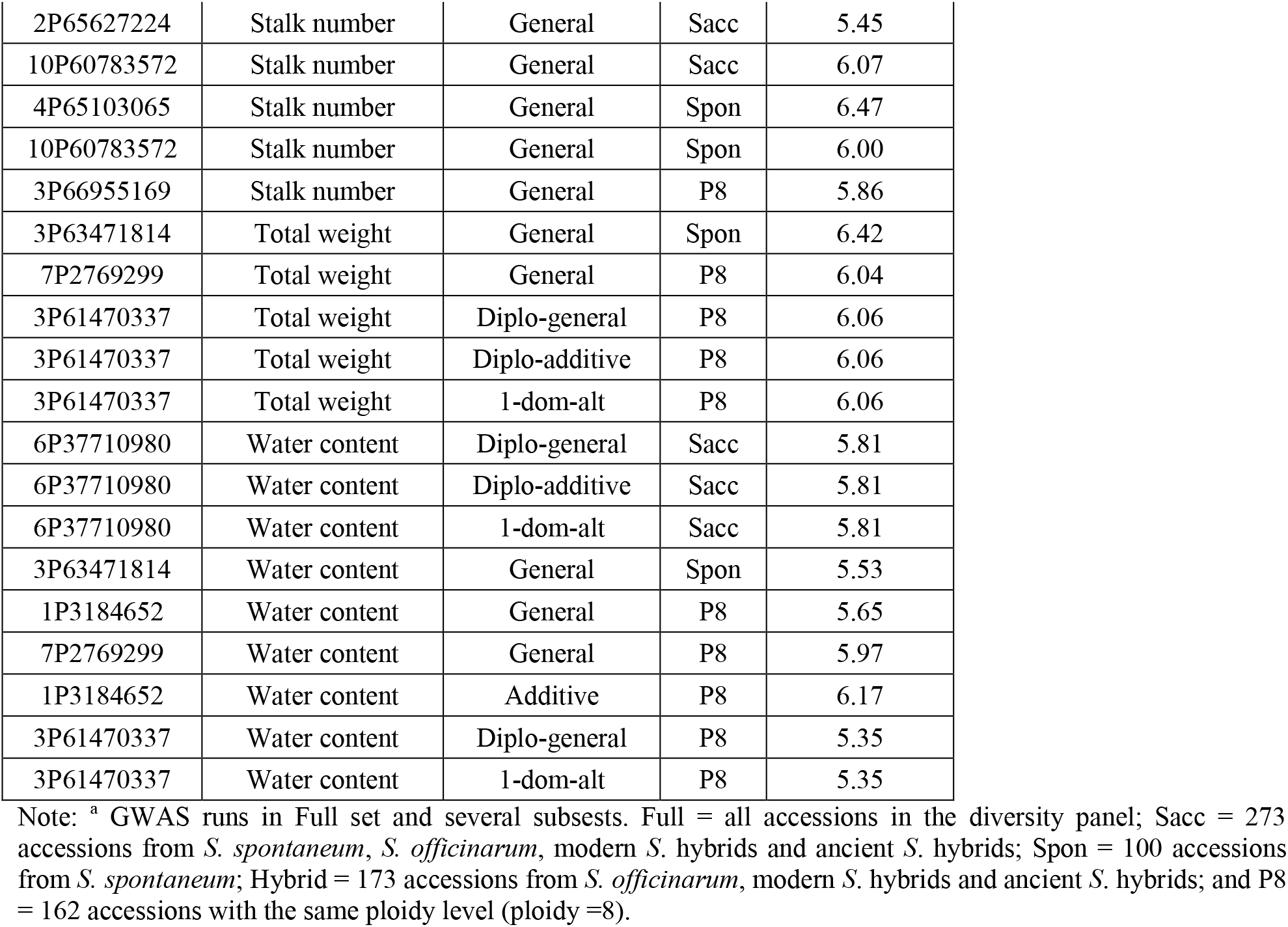
Associated DNA markers identified by genome-wide association study (GWAS) in sugarcane diversity panel.

**Table S3.**
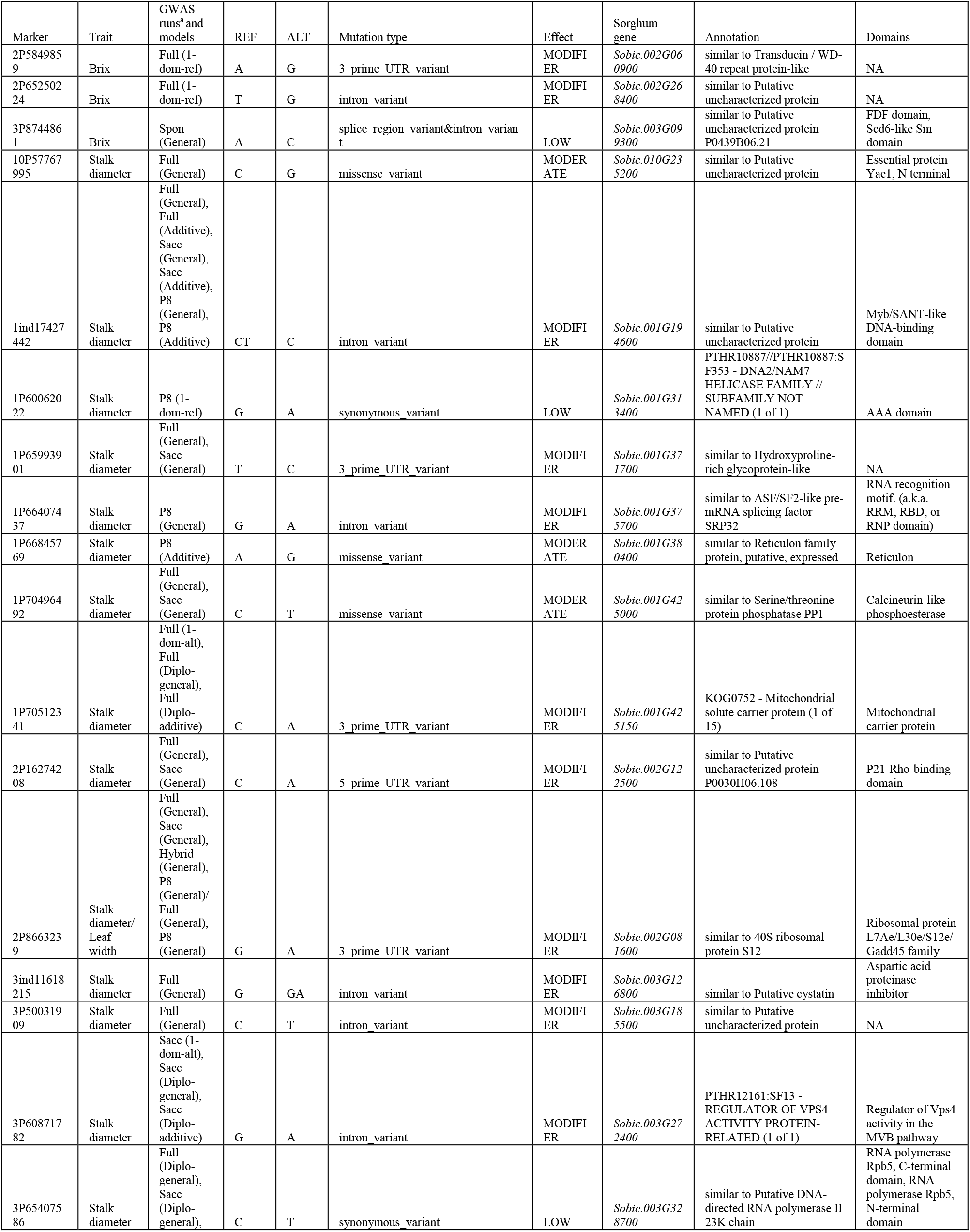
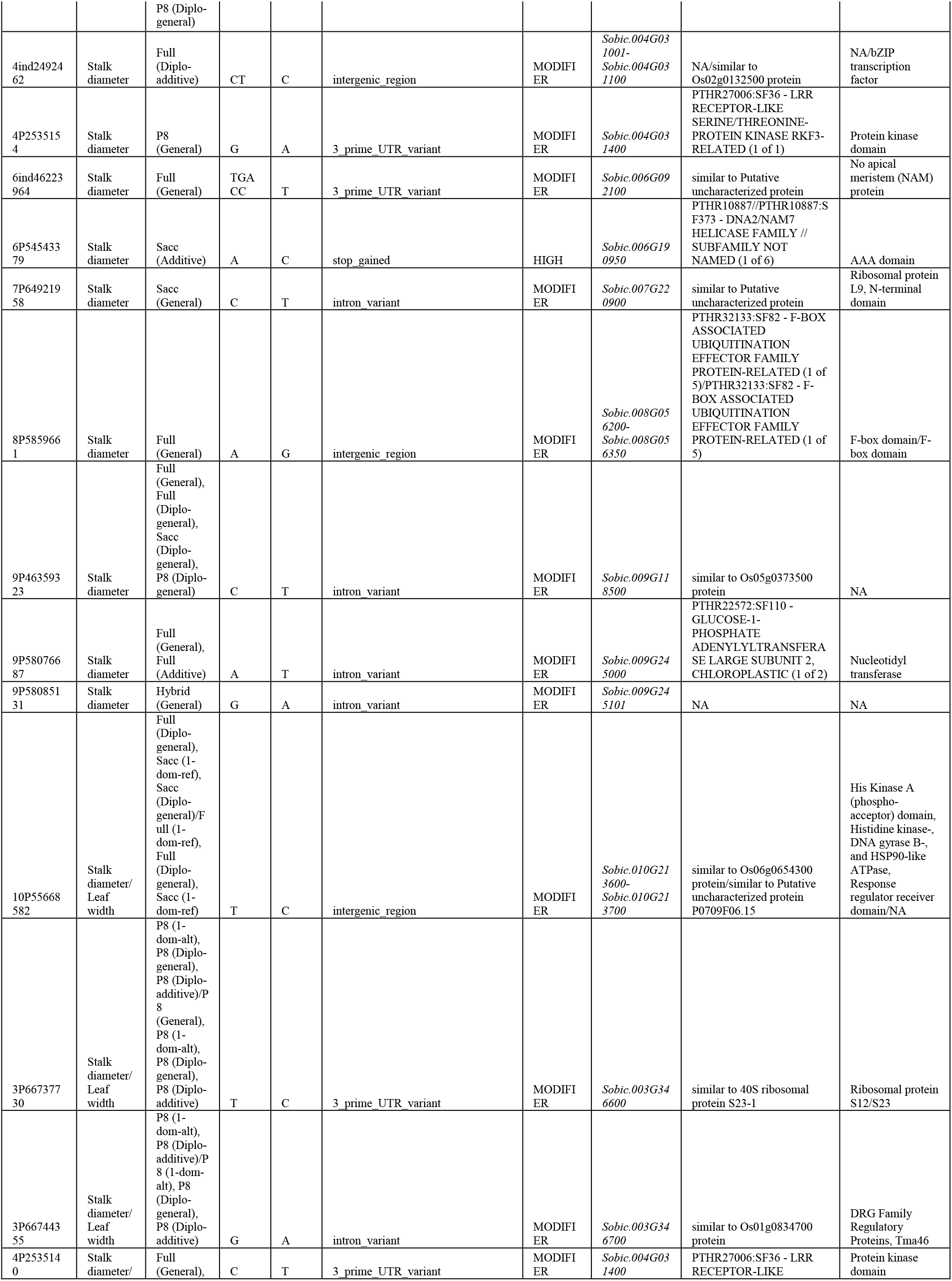
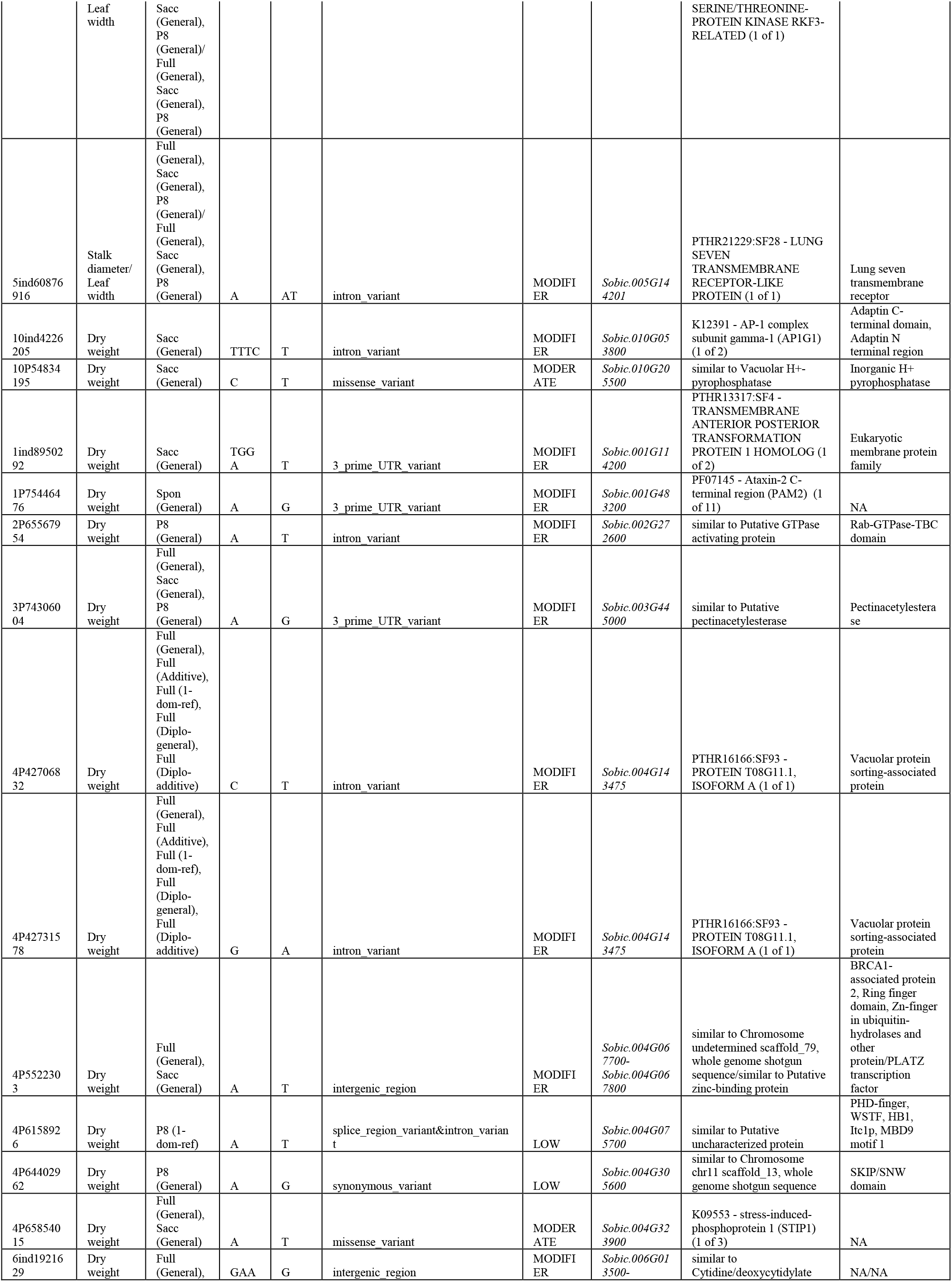
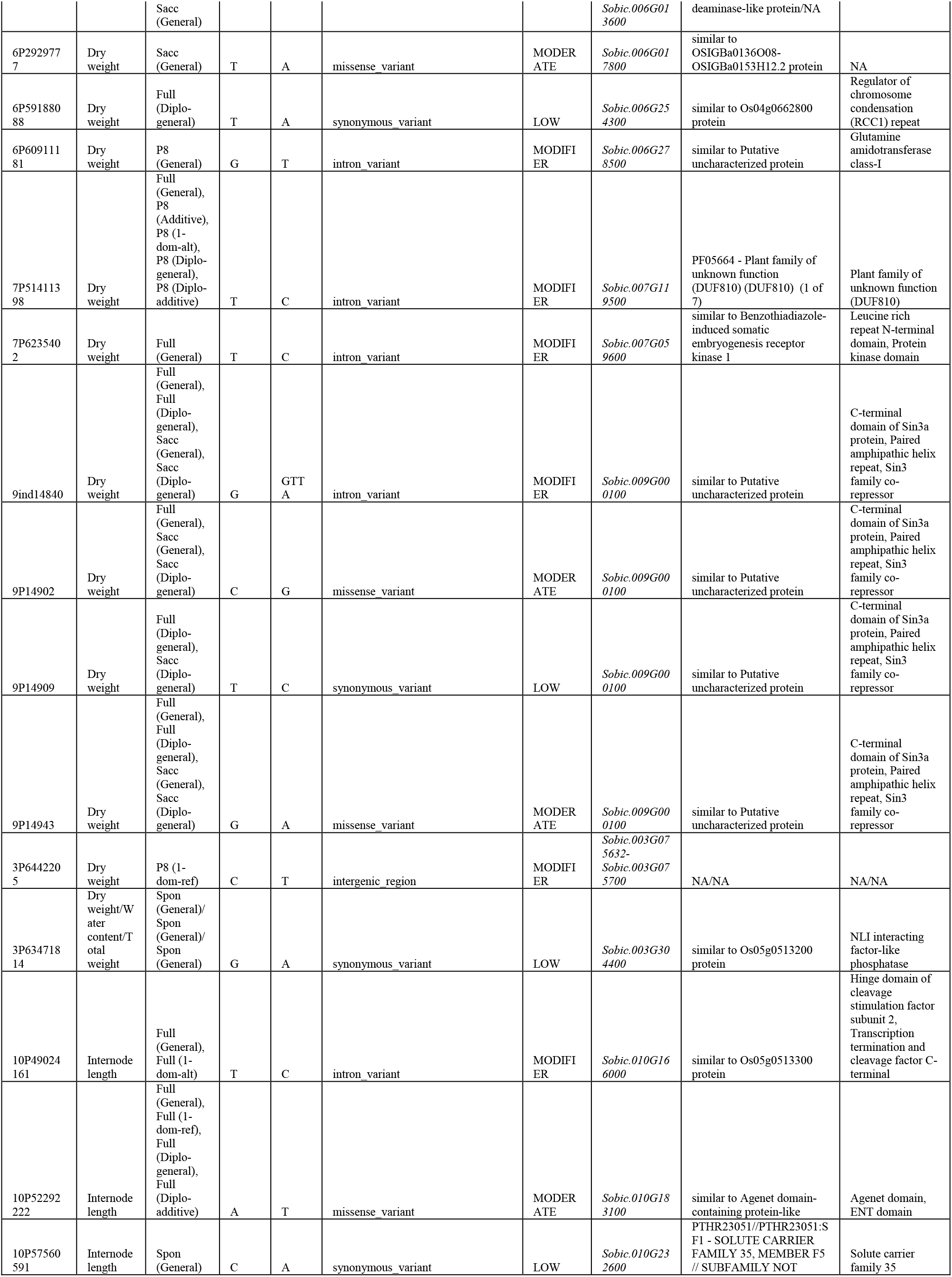
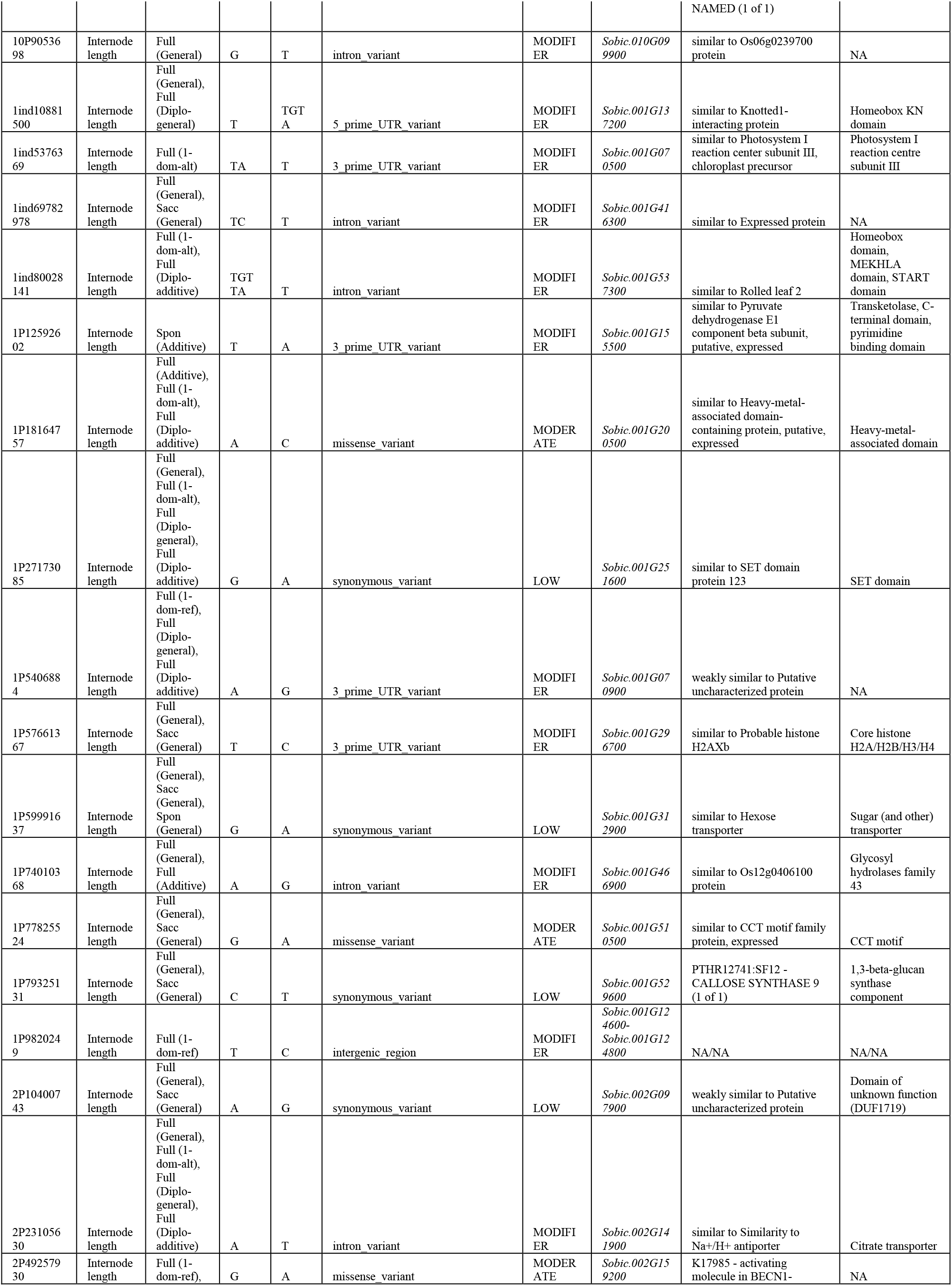
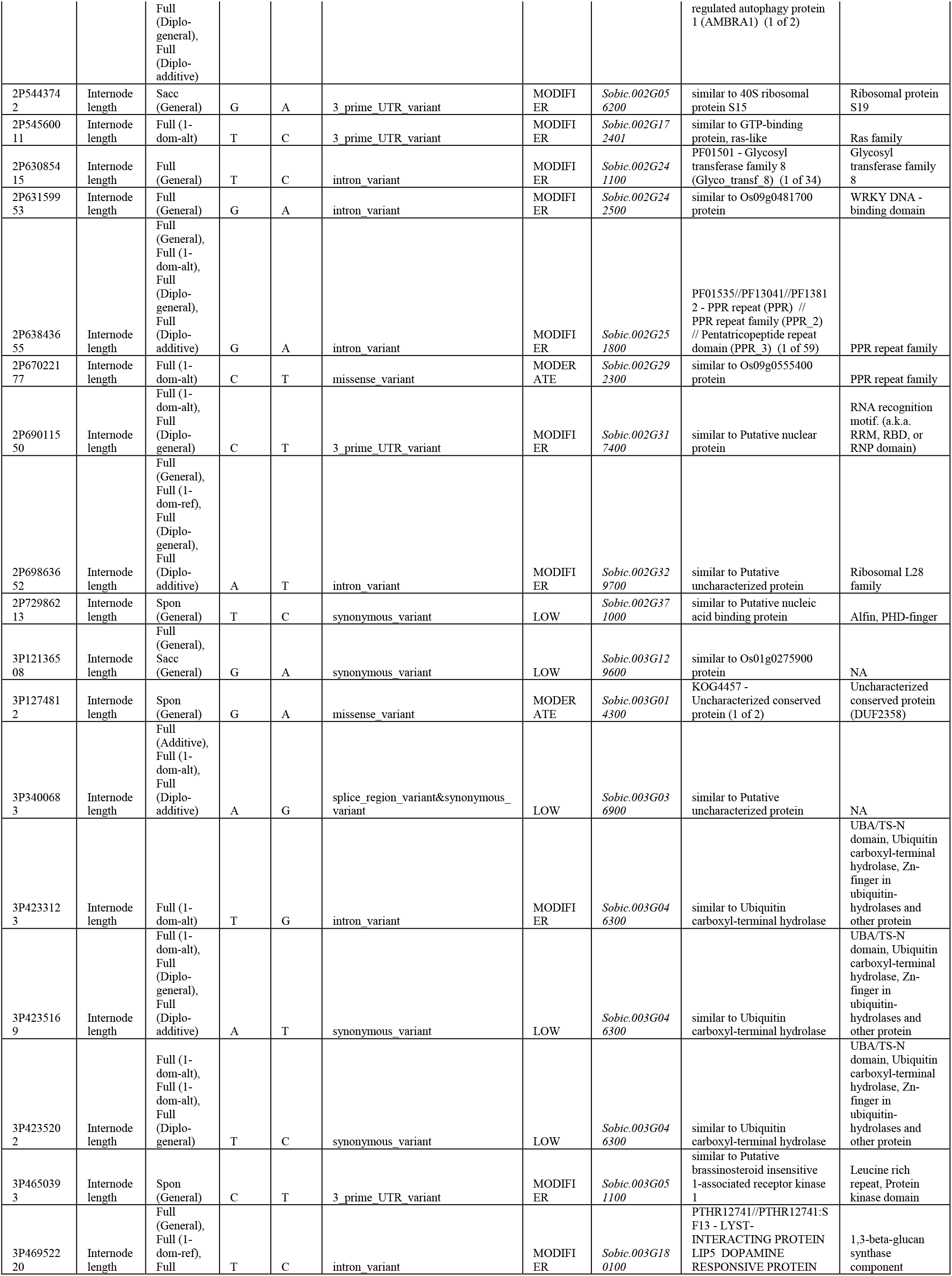
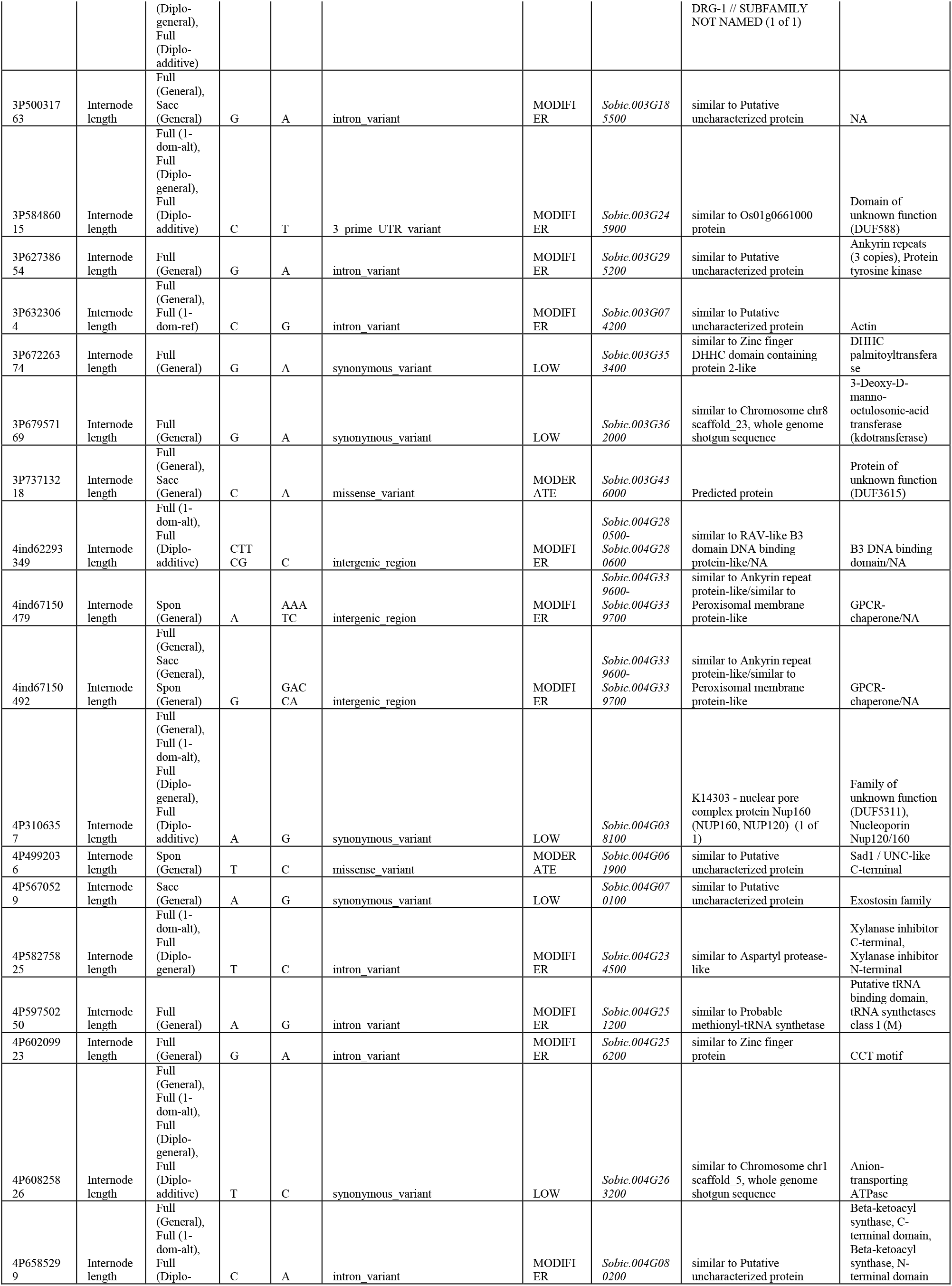
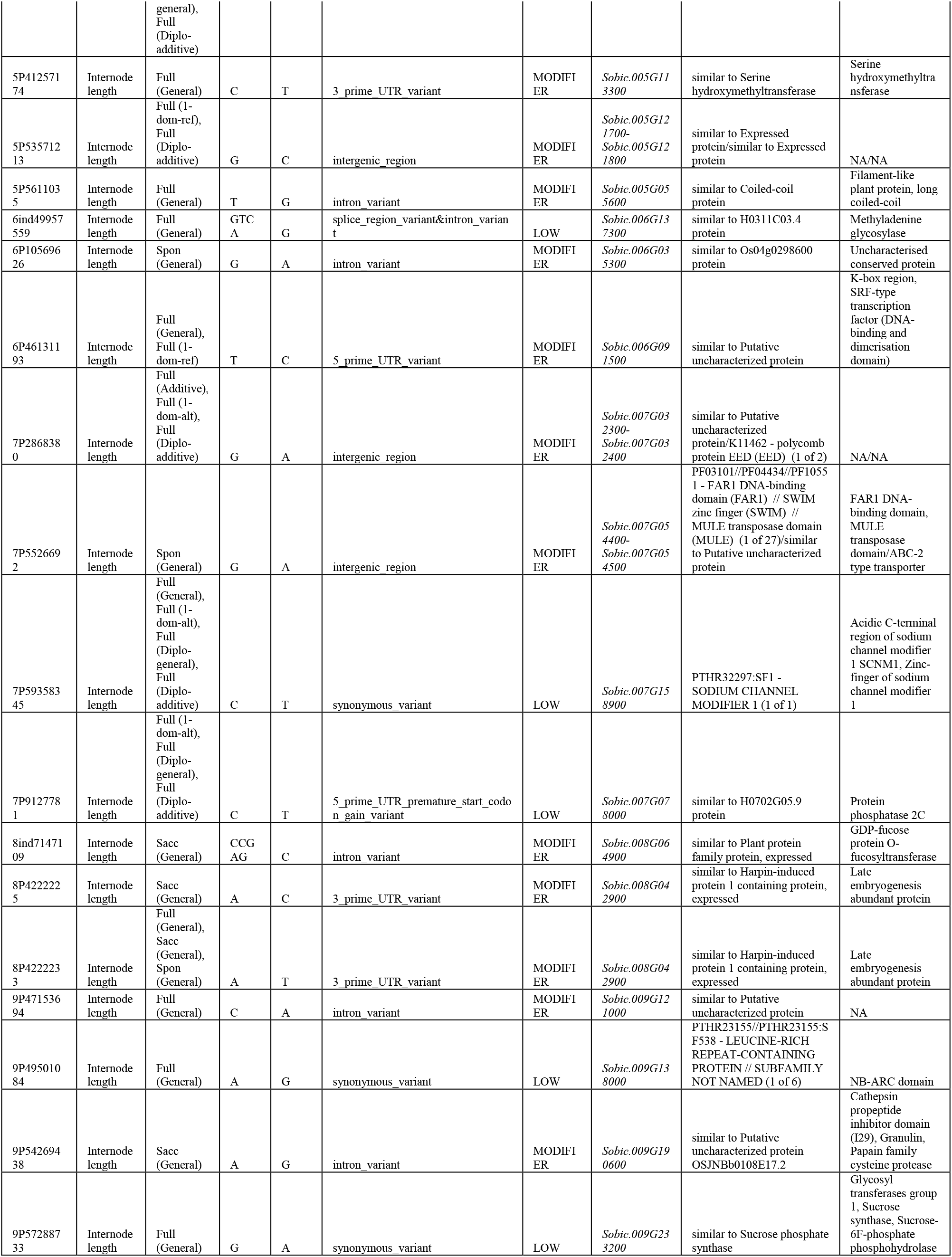
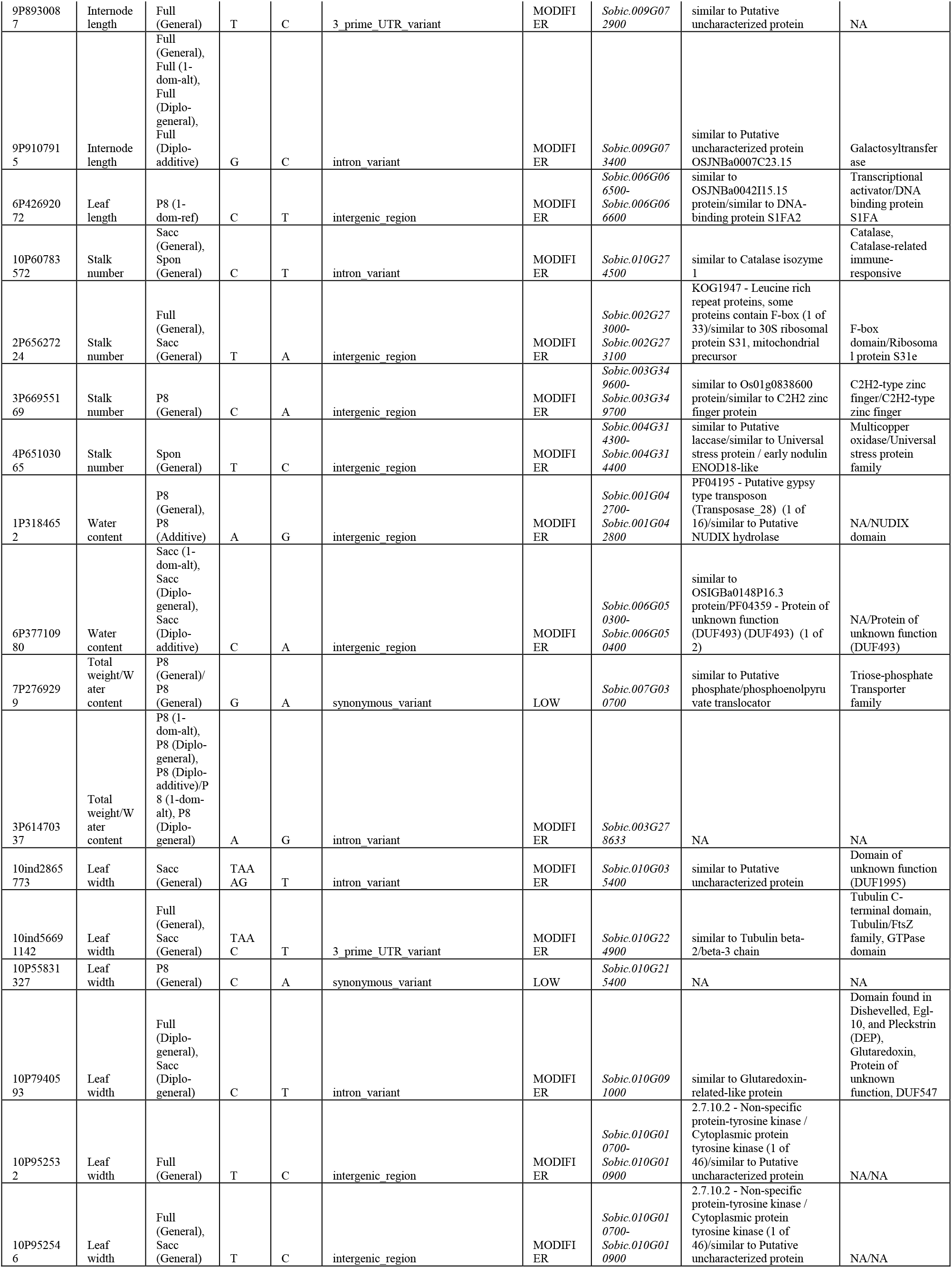
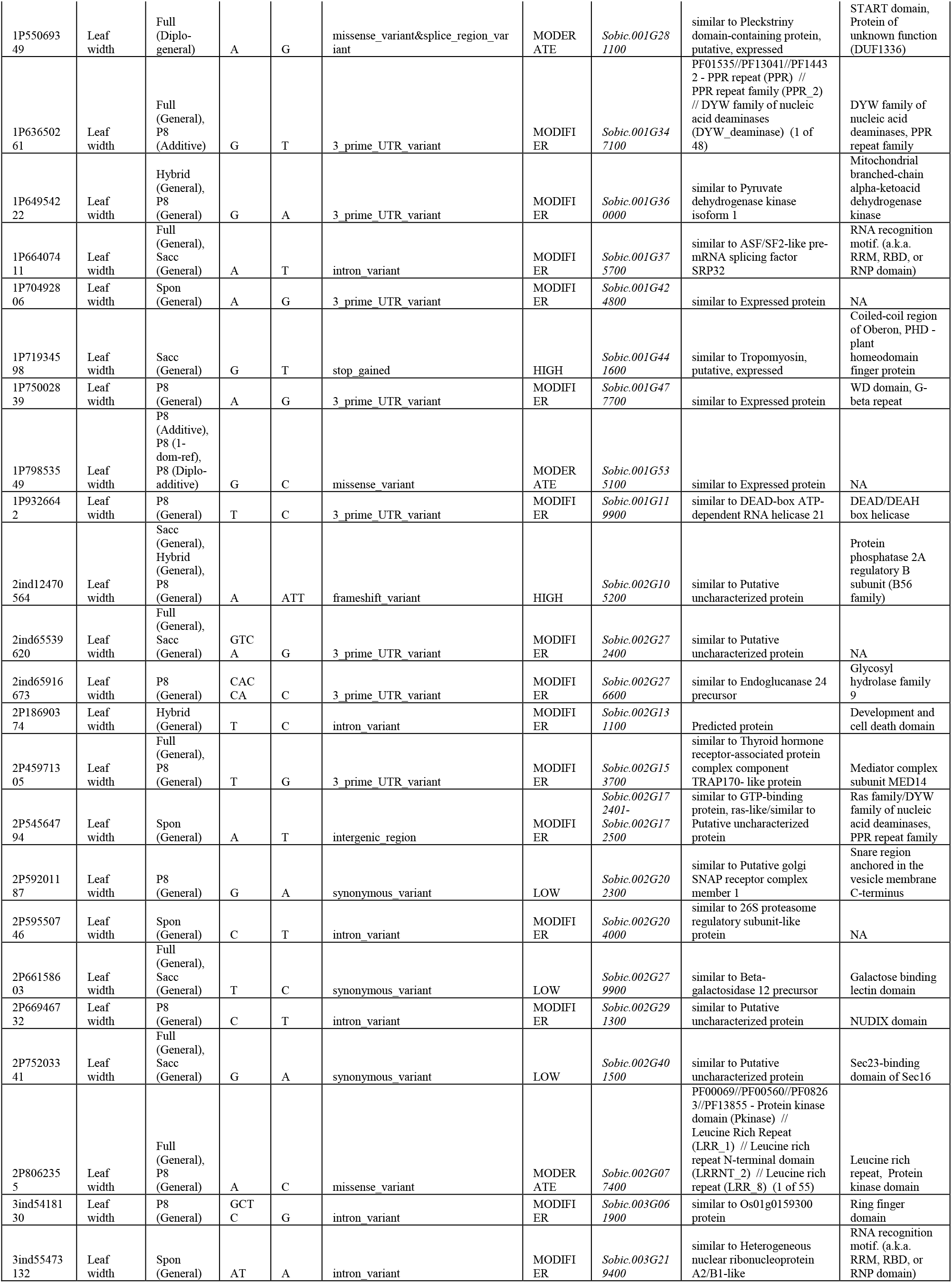
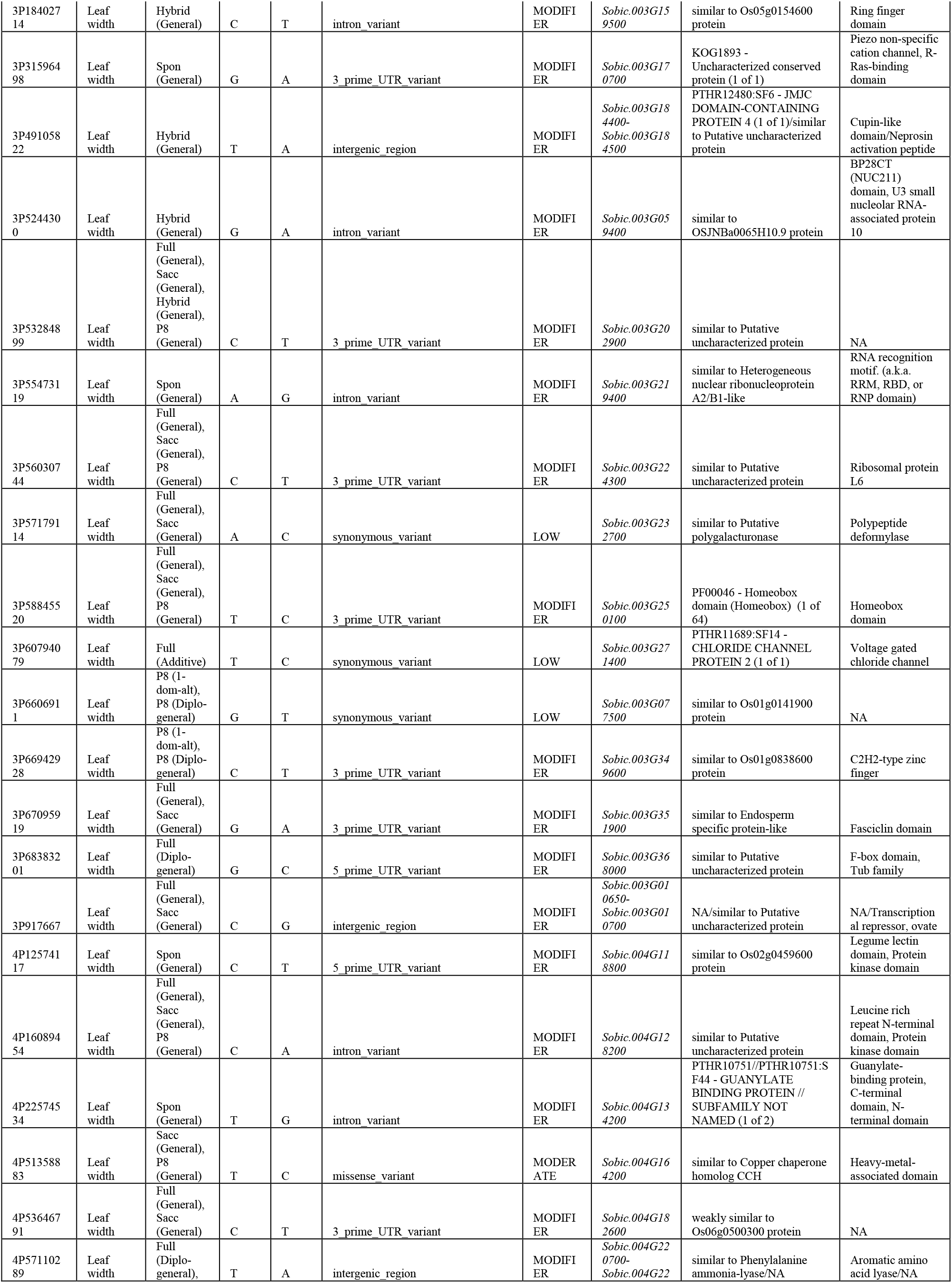
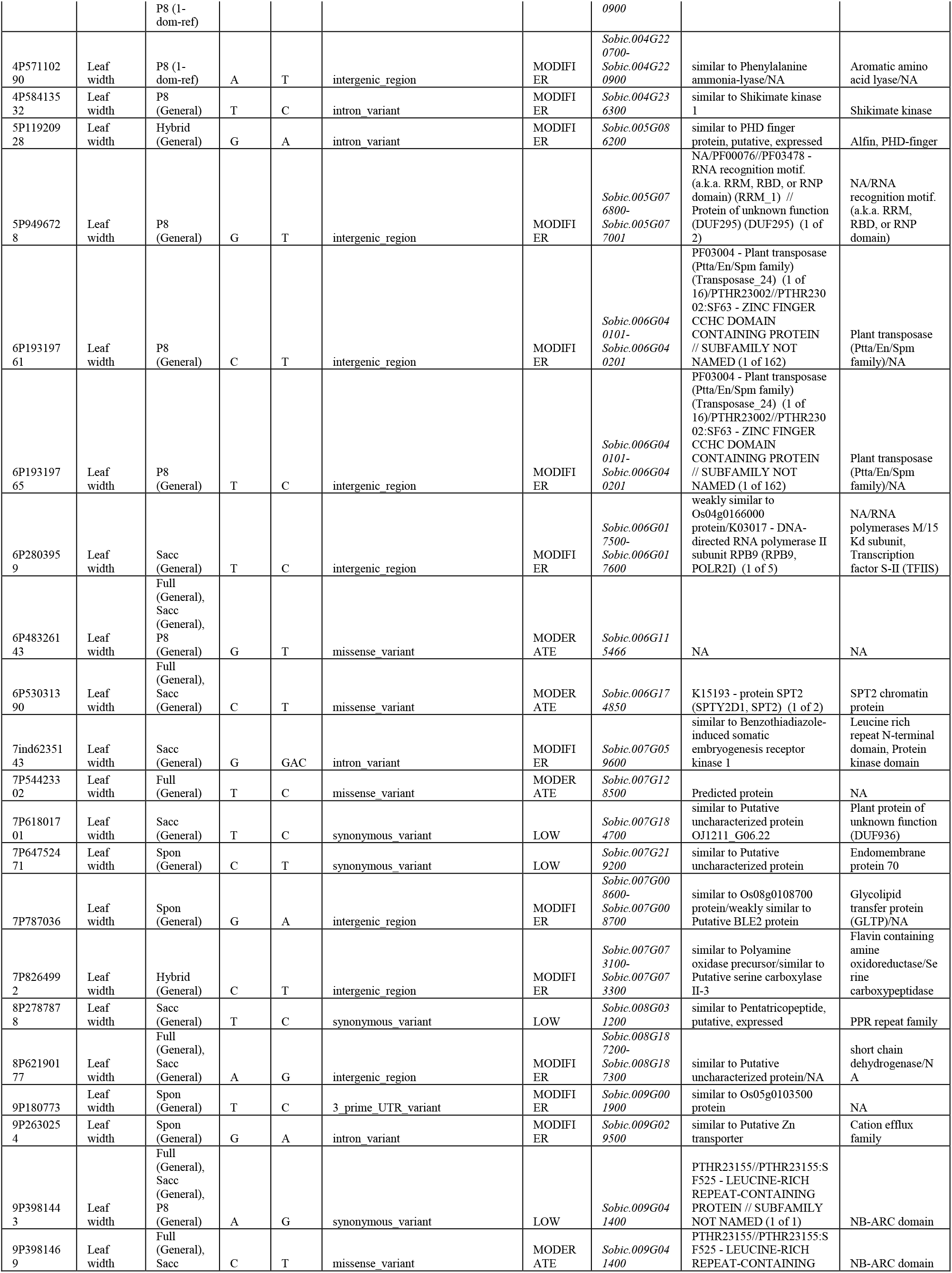
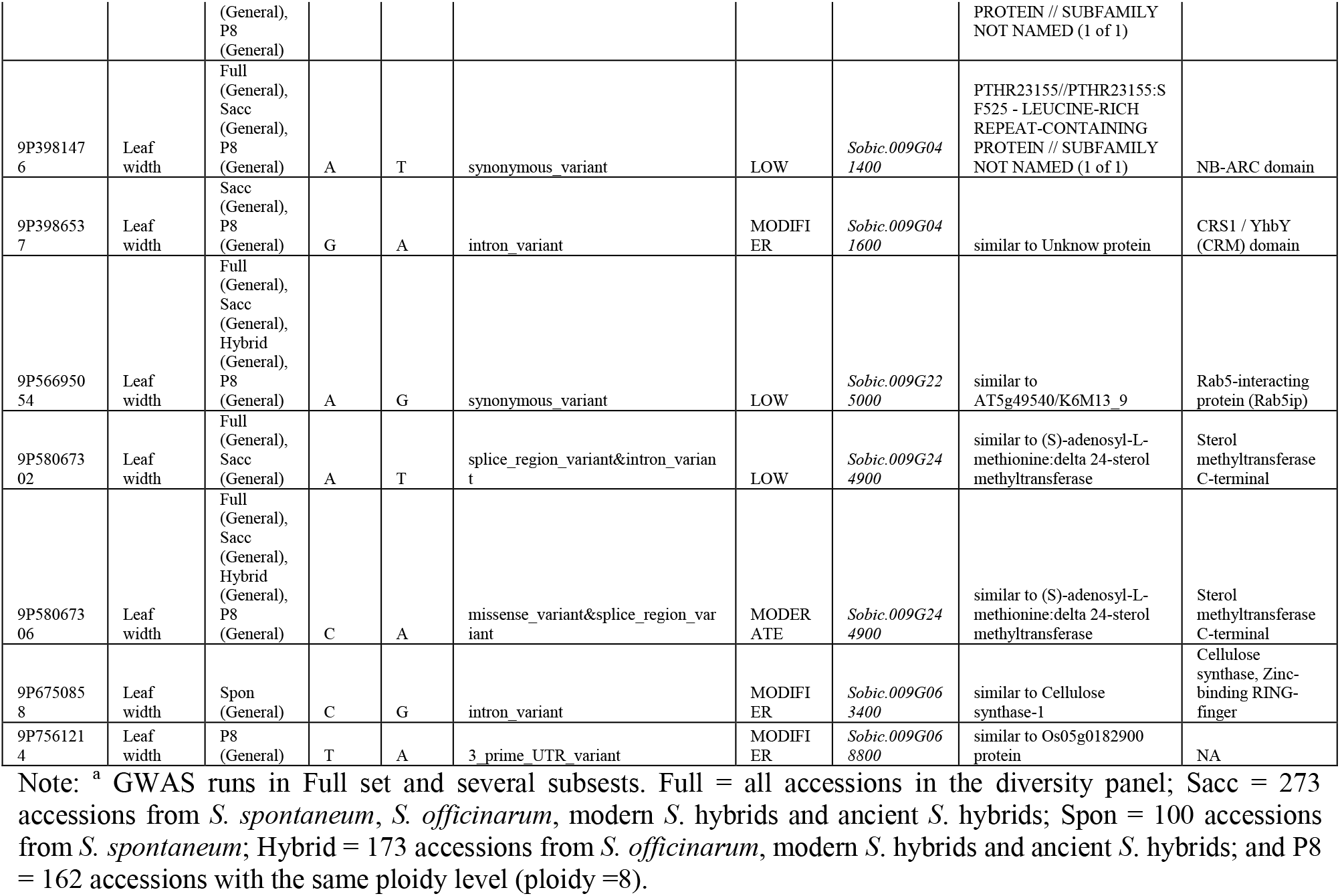
Non-redundant associated markers and candidate genes with genome-wide associations study (GWAS) hits.

